# Feature selection for preserving biological trajectories in single-cell data

**DOI:** 10.1101/2023.05.09.540043

**Authors:** Jolene S. Ranek, Wayne Stallaert, Justin Milner, Natalie Stanley, Jeremy E. Purvis

**Affiliations:** Department of Genetics, University of North Carolina at Chapel Hill, Chapel Hill, NC, USA; Computational Medicine Program, University of North Carolina at Chapel Hill, Chapel Hill, NC, USA; Department of Computational and Systems Biology, University of Pittsburgh, Pittsburgh, PA, USA; Department of Microbiology and Immunology, University of North Carolina at Chapel Hill, Chapel Hill, NC, USA; Department of Computer Science, University of North Carolina at Chapel Hill, Chapel Hill, NC, USA

## Abstract

Single-cell technologies can readily measure the expression of thousands of molecular features from individual cells undergoing dynamic biological processes, such as cellular differentiation, immune response, and disease progression. While examining cells along a computationally ordered pseudotime offers the potential to study how subtle changes in gene or protein expression impact cell fate decision-making, identifying characteristic features that drive continuous biological processes remains difficult to detect from unenriched and noisy single-cell data. Given that all profiled sources of feature variation contribute to the cell-to-cell distances that define an inferred cellular trajectory, including confounding sources of biological variation (e.g. cell cycle or metabolic state) or noisy and irrelevant features (e.g. measurements with low signal-to-noise ratio) can mask the underlying trajectory of study and hinder inference. Here, we present DELVE (dynamic selection of locally covarying features), an unsupervised feature selection method for identifying a representative subset of dynamically-expressed molecular features that recapitulates cellular trajectories. In contrast to previous work, DELVE uses a bottom-up approach to mitigate the effect of unwanted sources of variation confounding inference, and instead models cell states from dynamic feature modules that constitute core regulatory complexes. Using simulations, single-cell RNA sequencing data, and iterative immunofluorescence imaging data in the context of the cell cycle and cellular differentiation, we demonstrate that DELVE selects features that more accurately characterize cell populations and improve the recovery of cell type transitions. This feature selection framework provides an alternative approach for improving trajectory inference and uncovering co-variation amongst features along a biological trajectory. DELVE is implemented as an open-source python package and is publicly available at: https://github.com/jranek/delve.

## Introduction

High-throughput single-cell technologies, such as flow and mass cytometry [1, 2, 3], single-cell RNA sequencing [4, 5, 6, 7], and imaging-based profiling techniques [8, 9, 10, 11] have transformed our ability to study how cell populations respond and dynamically change during processes like development [12, 13, 14, 15] and immune response [16, 17, 18]. By profiling many features (e.g. proteins or genes) for many thousands of cells from a biological sample, these technologies provide high-dimensional snapshot measurements that can be used to gain fundamental insights into the molecular mechanisms that govern phenotypic changes.

Trajectory inference methods [19] have been developed to model dynamic biological processes from snapshot single-cell data. By assuming cells are asynchronously changing over time such that a profiled biological sample from a single experimental time point describes a range of the underlying dynamic process, computational trajectory inference approaches have leveraged minimum spanning tree approaches [20, 21, 22], curve-fitting [23, 24], graph-based techniques [25, 26, 27], probabilistic approaches [28, 29, 30], or optimal transport [31, 32] to order cells based on their similarities in feature expression. Once a trajectory model is fit, regression [33, 34, 35] can be performed along estimated pseudotime (e.g. distance through the inferred trajectory from a start cell) to identify specific cell state changes associated with differentiation or disease trajectories. Moreover, these inferred cellular trajectories have the potential to elucidate higher-order gene interactions [36], gene regulatory networks [37], predict cell fate probabilities [29], or find shared mechanisms of expression dynamics across disease conditions or species [38, 39].

While trajectory analysis has proven useful in the context of single-cell biology, the identification of characteristic genes or proteins that drive continuous biological processes relies on having inferred accurate cellular trajectories, which can be challenging, especially when trajectory inference is performed on the original full unenriched dataset. Single-cell data are noisy measurements that suffer from limitations in detection sensitivity, where dropout [40], low signal-to-noise, or sample degradation [41] can result in spurious signals that can overwhelm true biological differences. Furthermore, all profiled sources of feature variation contribute to the cell-to-cell distances that define the inferred cellular trajectory; thus, including confounding sources of biological variation (e.g. cell cycle, metabolic state) or irrelevant features (e.g. extracted imaging measurements that contain low signal-to-noise ratio) can distort or mask the intended trajectory of study [42, 43]. With the accumulation of large-scale single-cell data and multi-modal measurements [44], appropriate filtering of noisy, information-poor, or irrelevant features can serve as a crucial and necessary step for cell type identification, inference of dynamic phenotypes, and identification of punitive driver features (e.g. genes, proteins).

Feature selection methods [45] are a class of supervised or unsupervised approaches that can remove redundant or information-poor features prior to performing trajectory inference, and therefore, they have great potential for improving the interpretation of downstream analysis, while easing the computational burden by reducing dataset dimension. In the supervised-learning regime, classification-based [46] or information-theoretic approaches [47, 48] have been used to evaluate features according to their discriminative power or association with cell types. Despite having great power to detect biologically-relevant features, these methods rely on expensive or laborious manual annotations (e.g. cell types) which are often unavailable [49] thus precluding them from use. In the unsupervised-learning regime, computational approaches often aim to identify relevant features based on intrinsic properties of the complete dataset; however, these methods have some limitations with respect to retaining features that are useful for defining cellular trajectories. Namely, although unsupervised variance-based approaches [50, 51], which effectively sample features based on their overall variation across cells, have been extensively used to identify features that define cell types without the need for ground truth annotations, (1) they can be overwhelmed by noisy or irrelevant features that dominate data variance, and (2) are insensitive to lineage-specific features (e.g. transcription factors) that have a small variance and gradual progression of expression. Alternatively, unsupervised similarity-based [52, 53, 25] or subspace-learning [54, 55] feature selection methods evaluate features according to their association with a cell-similarity graph defined by all features or the underlying structure of the data (e.g. pairwise similarities defined by uniform manifold approximation and projection (UMAP) [56], eigenvectors of the graph Laplacian matrix [57]). While these approaches have the potential to detect smoothly varying genes or proteins that define cellular transitions, they rely on the cell-similarity graph from the full dataset and can fail to identify relevant features when the number of noisy features outweighs the number of informative ones [58, 59].

To address these limitations, we developed DELVE (dynamic selection of locally covarying features), an unsupervised feature selection method for identifying a representative subset of molecular features that robustly recapitulate cellular trajectories. In contrast to previous work [55, 52, 50, 54, 53, 25], DELVE uses a bottom-up approach to mitigate the effect of unwanted sources of variation confounding feature selection and trajectory inference, and instead models cell states from dynamic feature modules that constitute core regulatory complexes. Features are then ranked for selection according to their association with the underlying cell trajectory graph using data diffusion techniques. We demonstrate the power of our approach for improving inference of cellular trajectories through achieving an increased sensitivity to detect diverse and dynamically expressed features that better delineate cell types and cell type transitions from single-cell RNA sequencing and protein immunofluorescence imaging data. Overall, this feature selection framework provides an alternative approach for uncovering co-variation amongst features along a biological trajectory.

## Results

### Overview of the DELVE algorithm

We propose DELVE, an unsupervised feature selection framework for modeling dynamic cell state transitions using graph neighborhoods (Fig. 1). Our approach extends previous unsupervised similarity-based [52, 53, 25] or subspace-learning feature selection [55] methods by computing the dependence of each gene on the cellular trajectory graph structure using a two-step approach. Inspired by the molecular events that occur during differentiation, where the coordinated spatio-temporal expression of key regulatory genes govern lineage specification [60, 61, 62, 63], we reasoned that we can approximate cell state transitions by identifying groups of features that are temporally co-expressed or co-regulated along the underlying dynamic process.

**Figure 1:**
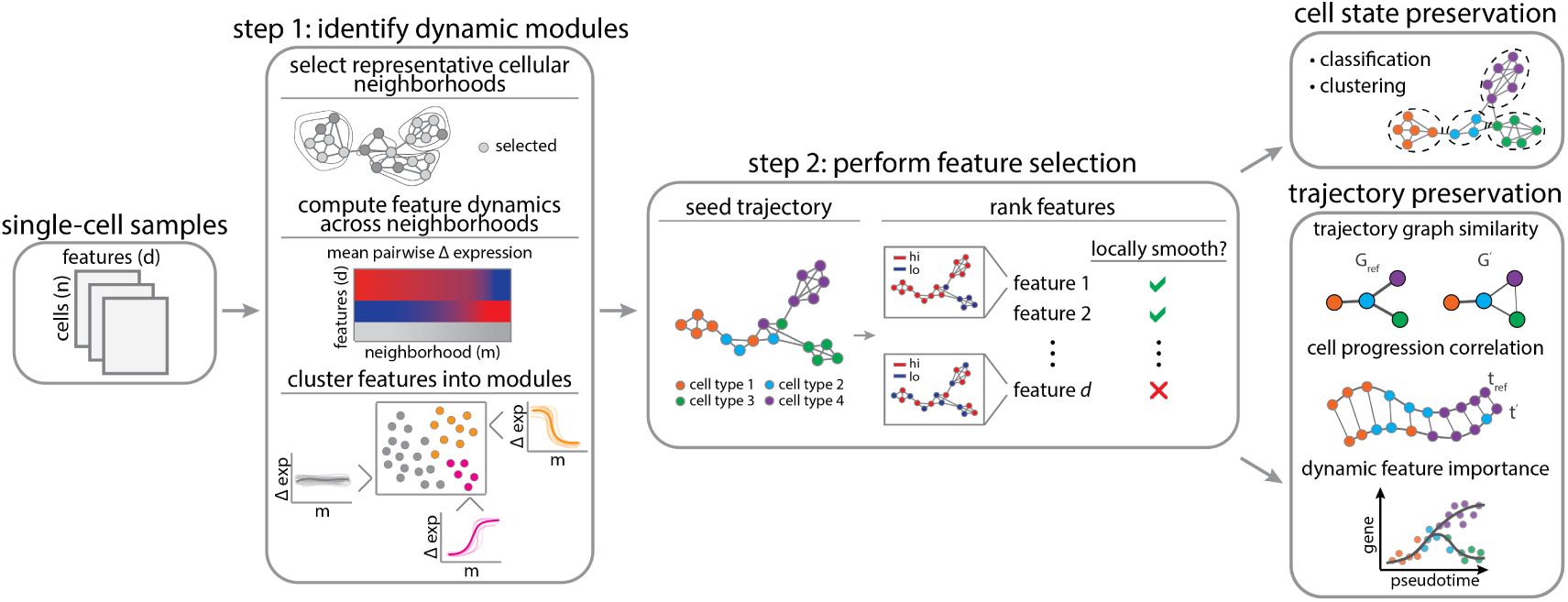
Schematic overview of the DELVE pipeline. Feature selection is performed in a two-step process. In step 1, DELVE clusters features according to their expression dynamics along local representative cellular neighborhoods defined by a weighted *k*-nearest neighbor affinity graph. Neighborhoods are sampled using a distribution-focused sketching algorithm that preserves cell-type frequencies and spectral properties of the original dataset [66]. A permutation test with a variance-based test statistic is used to determine if a set of features are (1) dynamically changing (dynamic) or (2) exhibiting random patterns of variation (static). In step 2, dynamic modules are used to seed or initialize an approximate cell trajectory graph and the trajectory is refined by ranking and selecting features that best preserve the local structure using the Laplacian Score [52]. In this study, we compare DELVE to the alternative unsupervised feature selection approaches on how well selected features preserve cell type and cell type transitions according to several metrics.

In step one, DELVE identifies groups of features that are temporally co-expressed by clustering features according to their average pairwise change in expression across prototypical cellular neighborhoods (Figure 1 step 1). As has been done previously [64, 65], we model cell states using a weighted *k*-nearest neighbor (*k*-NN) affinity graph, where nodes represent cells and edges describe the transcriptomic or proteomic similarity amongst cells according to all profiled features. Here, DELVE leverages a distribution-focused sketching method [66] to effectively sample cellular neighborhoods across all cell types. This sampling approach has three main advantages: (1) cellular neighborhoods are more reflective of the distribution of cell states, (2) redundant cell states are removed, and (3) fewer cellular neighborhoods are required to estimate feature dynamics resulting in increased scalability. Following clustering, each DELVE module contains a set of features with similar local changes in co-variation across prototypical cell states along the cellular trajectory. Feature-wise permutation testing is then used to assess the significance of dynamic expression variation across grouped features as compared to random assignment. By identifying and excluding modules of features that have static, random, or noisy patterns of expression variation, this approach effectively mitigates the effect of unwanted sources of variation confounding feature ranking and selection, and subsequent trajectory inference.

In step two, DELVE approximates the underlying cellular trajectory by constructing an affinity graph between cells, where cell similarity is now redefined according to a core set of dynamically expressed regulators. All profiled features are then ranked according to their association with the underlying cellular trajectory graph using graph signal processing techniques [67, 68] (Fig. 1 step 2). More concretely, a graph signal is any function that has a real defined value on all of the nodes. In this context, we consider all features as graph signals and rank them according to their total variation in expression along the cellular trajectory graph using the Laplacian Score [52]. Intuitively, DELVE retains features that are considered to be globally smooth, or have similar expression values amongst similar cells along the approximate cellular trajectory graph. In contrast, DELVE excludes features that have a high total variation in signal, or expression values that are rapidly oscillating amongst neighboring cells, as these features are likely noisy or not involved in the underlying dynamic process that was seeded. The output of DELVE is a ranked set of features that best preserve the local trajectory structure. For a more detailed description on the problem formulation, the mathematical foundations behind feature ranking, and the impact of nuisance features on trajectory inference, see DELVE in the Methods section.

### DELVE outperforms existing feature selection methods on representing cellular trajectories in the presence of single-cell RNA sequencing noise

Although feature selection is a common preprocessing step in single-cell analysis [69] with the potential to reveal cell-type transitions that would have been masked in the original high-dimensional feature space [42], there has been no systematic evaluation of feature selection method performance on identifying biologically-relevant features for trajectory analysis in single-cell data, especially in the context of noisy data that contain biological or technical challenges (e.g. low total mRNA count, low signal-to-noise ratio, or dropout). In this study, we compared DELVE to eleven other feature selection approaches and evaluated methods on their ability to select features that represent cell types and cell type transitions using simulated data where the ground truth was known. In the sections below, we will describe an overview of the feature selection methods considered and outline the simulation design and evaluation criteria in more detail. We will then provide qualitative and quantitative assessments of how noise impacts feature selection method performance and subsequent inference of cellular trajectories.

#### Overview of feature selection methods

We performed a systematic evaluation of twelve feature selection methods for preserving cellular trajectories in noisy single-cell data. Feature selection methods were grouped into five general categories prior to evaluation: supervised, similarity, subspace-learning, variance, and baseline approaches. For more details on the feature selection methods implemented and hyperparameters, see Benchmarked feature selection methods and Supplementary Table 1.

##### Supervised approaches

To illustrate the performance of *ground-truth* feature selection that could be obtained through supervised learning on expert annotated cell labels, we performed Random Forest classification. Random Forest classification [46] is a supervised ensemble learning algorithm that uses an ensemble of decision trees to partition the feature space such that all of the cells with the same cell type label are grouped together. Here, each decision or split of a tree was chosen by minimizing the Gini Impurity score [70]. This approach was included to provide context for unsupervised feature selection method performance.

##### Similarity approaches

We considered four similarity-based approaches as unsupervised feature selection methods that rank features according to their association with a cell similarity graph defined by all profiled features (e.g. Laplacian Score, neighborhood variance, hotspot) or dynamically-expressed features (e.g. DELVE). First, the Laplacian Score (LS) [52] is an unsupervised locality-preserving feature selection method that ranks and selects features according to (1) the total variation in feature expression across neighboring cells using a cell similarity graph defined by all features and (2) a feature’s global variance. Next, neighborhood variance [25] is an unsupervised feature selection method that selects features with gradual changes in expression for building biological trajectories. Here, features are selected if their variance in expression across local cellular neighborhoods is smaller than their global variance. Hotspot [53] performs unsupervised feature selection through a local autocorrelation test statistic that measures the association of a gene’s expression with a cell similarity graph defined by all features. Lastly, DELVE (dynamic selection of locally covarying features) is an unsupervised feature selection method that ranks features according to their association with the underlying cellular trajectory graph. First, features are clustered into modules according to changes in expression across local representative cellular neighborhoods. Next, modules of features with dynamic expression patterns (denoted as *dynamic seed*) are used to construct an approximate cellular trajectory graph. Features are then ranked according to their association with the approximate cell trajectory graph using the Laplacian Score [52].

##### Subspace learning approaches

We considered two subspace-learning feature selection methods as unsupervised methods that rank features according to how well they preserve the overall cluster structure (e.g. MCFS) or manifold structure (e.g. SCMER) of the data. First, multi-cluster feature selection (MCFS) [55] is an unsupervised feature selection method that selects features that best preserve the multi-cluster structure of data by solving an L1 regularized least squares regression problem on the spectral embedding defined by all profiled features. The optimization is solved using the least angles regression algorithm [71]. Next, single-cell manifold-preserving feature selection (SCMER) [54] is an unsupervised feature selection method that selects a subset of features that best preserves the pairwise similarity matrix between cells defined in uniform manifold approximation and projection [56] based on all profiled features. To do so, it uses elastic net regression to find a sparse solution that minimizes the KL divergence between a pairwise similarity matrix between cells defined by all features and one defined using only the selected features.

##### Variance approaches

We considered two variance-based feature selection approaches (e.g. highly variable genes [50], max variance) as unsupervised methods that use global expression variance as a metric for ranking feature importance. First, highly variable gene selection (HVG) [50] is an unsupervised feature selection method that selects features according to a normalized dispersion measure. Here, features are binned based on their average expression. Within each bin, genes are then z-score normalized to identify features that have a large variance, yet a similar mean expression. Next, max variance is an unsupervised feature selection method that ranks and selects features that have a large global variance in expression.

##### Baseline approaches

We considered three baseline strategies (e.g. all, random, dynamic seed) that provide context for the overall performance of feature selection. First, all features illustrates the performance when feature selection is not performed and all features are included for analysis. Second, random features represents the performance quality when a random subset of features are sampled. Lastly, dynamic seed features indicate the performance from dynamically-expressed features identified in step 1 of the DELVE algorithm prior to feature ranking and selection.

#### Single-cell RNA sequencing simulation design

To validate our approach and benchmark feature selection methods on representing cellular trajectories, we simulated 90 single-cell RNA sequencing datasets with 1500 cells and 500 genes using Splatter. Splatter [72] simulates single-cell RNA sequencing data with various trajectory structures (e.g. linear, bifurcation, tree) using a gamma-Poisson hierarchical model. Importantly, this approach provides ground truth reference information (e.g. cell type annotations, differentially expressed genes per cell type and trajectory, and a latent vector that describes an individual cell’s progression through the trajectory) that we can use to robustly assess feature selection method performance, as well as quantitatively evaluate the limitations of feature selection strategies for trajectory analysis. Moreover, to comprehensively evaluate feature selection methods under common biological and technical challenges associated with single-cell RNA sequencing data, we added relevant sources of single-cell noise to the simulated data. First, we simulated low signal-to-noise ratio by enforcing a mean-variance relationship amongst genes; this ensures that lowly expressed genes are more variable than highly expressed genes. Next, we modified the total number of profiled mRNA transcripts, or library size. This has been shown previously to vary amongst cells within a single-cell experiment and can influence both the detection of differentially expressed genes [73], as well as impact the reproducibility of the inferred lower-dimensional embedding [74]. Lastly, we simulated the inefficient capture of mRNA molecules, or dropout, by undersampling gene expression from a binomial distribution; this increases the amount of sparsity present within the data. For more details on the splatter simulation, see *Splatter simulation*. For each simulated trajectory, we performed feature selection according to all described feature selection strategies, and considered the top 100 ranked features for downstream analysis and evaluation.

#### Qualitative assessment of feature selection method performance

Prior to evaluating feature selection method performance quantitatively, we began our analysis with a qualitative assessment of the importance of feature selection for representing cellular trajectories when the data contain irrelevant or noisy genes. First, we visually compared the cellular trajectories generated from a feature selection strategy with PHATE (potential of heat-diffusion for affinity-based transition embedding). PHATE [75] is a nonlinear dimensionality reduction method that has been shown to effectively learn and represent the geometry of complex continuous and branched biological trajectories. As an illustrative example, Fig. 2a shows the PHATE embeddings for simulated linear differentiation trajectories generated from four feature selection approaches (all, DELVE, Laplacian Score (LS), and random) when subjected to a decrease in the signal-to-noise ratio. Here, we simulated a reduction in the signal-to-noise ratio and stochastic gene expression by modifying the biological coefficient of variation (BCV) parameter within Splatter [72]. This scaling factor controls the mean-variance relationship between genes, where lowly expressed genes are more variable than highly expressed genes (See *Splatter simulation*). Under low noise conditions where the data contained a high signal-to-noise ratio, we observed that excluding irrelevant features with DELVE or the Laplacian Score (LS) produced a much smoother, denoised visualization of the linear trajectory, where cells were more tightly clustered according to cell type. This was compared to the more diffuse presentation of cell states obtained based on all genes. We then examined how noise influences the quality of selected features from a feature selection strategy. As the signal-to-noise ratio decreased (high, medium, low), we observed that the linear trajectory became increasingly harder to distinguish, whereby including both irrelevant and noisy genes often masked the underlying trajectory structure (Fig. 2a all genes, medium to low signal-to-noise ratio). Furthermore, we found that unsupervised similarity-based or subspace learning feature selection methods that initially define a cell similarity graph according to all irrelevant, noisy, and informative genes often selected genes that produced noisier embeddings as the amount of noise increased (e.g. Fig. 2a LS: medium signal-to-noise ratio), as compared to DELVE (e.g. Fig. 2a DELVE medium signal-to-noise ratio). We reason that this is due to spurious similarities amongst cells, reduced clusterability, and increased diffusion times. These qualitative observations were consistent across different noise conditions (e.g. decreased signal-to-noise, decreased library size, increased dropout) and trajectory types (e.g. linear, bifurcation, tree) (See Supplementary Figs. 1 - 9). Although a qualitative comparison, this example illustrates how including irrelevant or noisy genes can define spurious similarities amongst cells, which can (1) influence a feature selection method ability to identify biologically-relevant genes and (2) impact the overall quality of an inferred lower dimensional embedding following selection. Given that many trajectory inference methods use lower dimensional representations in order to infer a cell’s progression through a differentiation trajectory, it is crucial to remove information-poor features prior to performing trajectory inference in order to obtain high quality embeddings, clustering assignments, or cellular orderings that are reproducible for both qualitative interpretation and downstream trajectory analysis.

**Figure 2:**
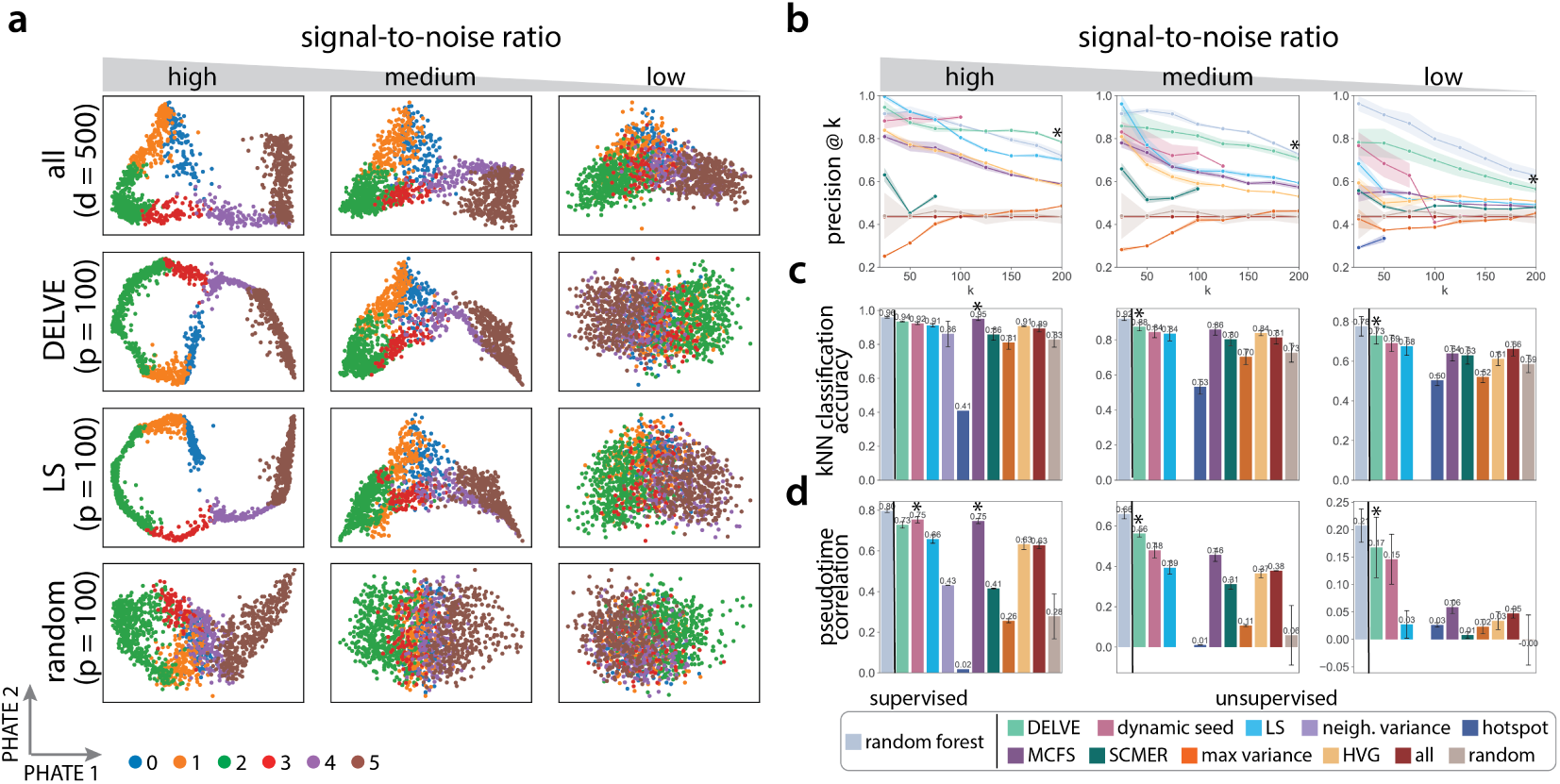
Comparison of feature selection methods on preserving linear trajectories when subjected to a reduction in the signal-to-noise ratio. (a) Example PHATE [75] visualizations of simulated linear trajectories using four feature selection approaches (all features, DELVE, Laplacian Score (LS) [52], and random selection) when subjected to a reduction in the signal-to-noise ratio (high, medium, low). Here, we simulated a reduction in the signal-to-noise ratio and stochastic gene expression by modifying the biological coefficient of variation (bcv) parameter within Splatter [72] that controls the mean-variance relationship between genes, where lowly expressed genes are more variable than highly expressed genes (high: bcv = 0.1, medium: bcv = 0.25, low: bcv = 0.5). *d* indicates the total number of genes (*d* = 500) and *p* indicates the number of selected genes following feature selection (*p* = 100). (b-d) Performance of twelve different feature selection methods: random forest [46], DELVE, dynamic seed features, LS [52], neighborhood variance [25], hotspot [53], multi-cluster feature selection (MCFS) [55], single-cell manifold preserving feature selection (SCMER) [54], max variance, highly variable gene selection (HVG) [50], all features, random features. Following feature selection, trajectory preservation was quantitatively assessed according to several metrics: (b) the precision of differentially expressed genes at *k* selected genes, (c) *k*-NN classification accuracy, and (d) pseudotime correlation to the ground truth cell progression across 10 random trails. Error bars/bands represent the standard deviation. * indicates the method with the highest median score. For further details across other trajectory types and noise conditions, see Supplementary Figs. 1 - 9.

#### Quantitative assessment of feature selection method performance

We next quantitatively examined how biological or technical challenges associated with single-cell RNA sequencing data may influence a feature selection method’s ability to detect the particular genes that define cell types or cell type transitions. To do so, we systematically benchmarked the 12 described feature selection strategies on their capacity to preserve trajectories according to three sets of quantitative comparisons. Method performance was assessed by evaluating if selected genes from an approach were (1) differentially expressed within a cell type or along a lineage, (2) could be used to classify cell types, and (3) could accurately estimate individual cell progression through the cellular trajectory. Fig. 2b-d shows feature selection method performance for simulated linear differentiation trajectories when subjected to the technical challenge of having a reduction in the signal-to-noise ratio.

First, we assessed the biological relevancy of selected genes, as well as the overall recovery of relevant genes as the signal-to-noise ratio decreased by computing a precision score. Precision@k is a metric that defines the proportion of selected genes (*k*) that are known to be differentially-expressed within a cell type or along a lineage (See Precision@k). Overall, we found that DELVE achieved the highest precision@k score between selected genes and the ground truth, validating that our approach was able to select genes that are differentially expressed and was the strongest in defining cell types and cell type transitions (See Fig. 2b). Importantly, DELVE’s ability to recover informative genes was robust to the number of genes selected (*k*) and to the amount of noise present in the data. In contrast, variance-based, similarity-based, or subspace-learning approaches exhibited comparatively worse recovery of cell type and lineage-specific differentially expressed genes.

Given that a key application of single-cell profiling technologies is the ability to identify cell types or cell states that are predictive of sample disease status, responsiveness to drug therapy, or are correlated with patient clinical outcomes [76, 77, 78, 79, 65], we then assessed whether selected genes from a feature selection strategy can correctly classify cells according to cell type along the underlying cellular trajectory; this is a crucial and necessary step of trajectory analysis. Therefore, we trained a *k*-nearest neighbor (*k*-NN) classifier on the selected feature set (see *k-nearest neighbor classification*) and compared the predictions to the ground truth cell type annotations by computing a cell type classification accuracy score. Across all simulated trajectories, we found that DELVE selected genes that often achieved the highest median *k*-NN classification accuracy score (high signal-to-noise ratio: 0.937, medium signal-to-noise ratio: 0.882, low signal-to-noise ratio: 0.734) and produced *k*-NN graphs that were more faithful to the underlying biology (See Fig. 2c). Moreover, we observed a few results that were consistent with the qualitative interpretations. First, removing irrelevant genes with DELVE, LS, or MCFS achieved higher *k*-NN classification accuracy scores (e.g. high signal-to-noise ratio; DELVE = 0.937, LS = 0.915, and MCFS = 0.955, respectively) than was achieved by retaining all genes (all = 0.900). Next, DELVE outperformed the Laplacian Score, suggesting that using a bottom-up framework and excluding nuisance features prior to performing ranking and selection is crucial for recovering cell-type specific genes that would have been missed if the cell similarity graph was initially defined based on all genes. Lastly, when comparing the percent change in performance as the amount of noise corruption increased (e.g. high signal-to-noise ratio to medium signal-to-noise ratio) for linear trajectories, we found that DELVE often achieved the highest average classification accuracy score (0.905) and lowest percent decrease in performance (−6.398%), indicating that DELVE was the most robust unsupervised feature selection method to noise corruption (See Supplementary Fig. 10a). In contrast, the existing unsupervised similarity-based or subspace learning feature selection methods that achieved high to moderate average *k*-NN classification accuracy scores (e.g. MCFS = 0.905, LS = 0.874) had larger decreases in performance (e.g. MCFS = -9.673%, LS = -8.390%) as the amount of noise increased. This further highlights the limitations of current feature selection methods on identifying cell type-specific genes from noisy single-cell omics data.

Lastly, when undergoing dynamic biological processes such as differentiation, cells exhibit a continuum of cell states with fate transitions marked by linear and nonlinear changes in gene expression [80, 81, 82]. Therefore, we evaluated how well feature selection methods could identify genes that define complex differentiation trajectories and correctly order cells along the cellular trajectory in the presence of noise. To infer cellular trajectories and to estimate cell progression, we used the diffusion pseudotime algorithm [83] on the selected gene set from each feature selection strategy, as this approach has been shown previously [19] to perform reasonably well for inference of simple or branched trajectory types (See Trajectory inference and analysis). Method performance was then assessed by computing the Spearman rank correlation between estimated pseudotime and the ground truth cell progression. We found that DELVE approaches more accurately inferred cellular trajectories and achieved the highest median pseudotime correlation to the ground truth measurements, as compared to alternative methods or all features (See Fig. 2d). Furthermore, similar to the percent change in classification performance, we found that DELVE was the most robust unsupervised feature selection method in estimating cell progression, as it often achieved the highest average pseudotime correlation (0.645) and lowest percent decrease in performance (−22.761%) as the amount of noise increased (See Supplementary Fig. 10b high to medium signal-to-noise ratio). In contrast, the alternative methods incorrectly estimated cellular progression and achieved lower average pseudotime correlation scores (e.g. MCFS = 0.602, LS = 0.526) and higher decreases in performance as the signal-to-noise ratio decreased (MCFS = -38.884%, LS = -40.208%).

We performed this same systematic evaluation across a range of trajectory types (e.g. linear, bifurcation, tree) and biological or technical challenges associated with single-cell data (See Supplementary Figs. 1 - 12). Fig. 3 displays the overall ranked method performance of feature selection methods on preserving cellular trajectories when subjected to different sources of single-cell noise (pink: decreased signal-to-noise ratio, green: decreased library size, and blue: increased dropout). Ranked aggregate scores were computed by averaging results across all datasets within a condition; therefore, this metric quantifies how well a feature selection strategy can recover genes that define cell types or cell type transitions underlying a cellular trajectory when subjected to that biological or technical challenge (See Aggregate scores). Across all conditions, we found that DELVE often achieved an increased recovery of differentially expressed genes, higher cell type classification accuracy, higher correlation of estimated cell progression, and lower percent change in performance in noisy data. While feature selection method performance varied across biological or technical challenges, we found that the Laplacian score (LS) and multi-cluster feature selection (MCFS) performed reasonably well under low amounts of noise corruption and are often the second and third ranked unsupervised methods. Altogether, this simulation study demonstrates that DELVE more accurately recapitulates cellular dynamics and can be used to effectively interrogate cell identity and lineage-specific gene expression dynamics from noisy single-cell data.

**Figure 3:**
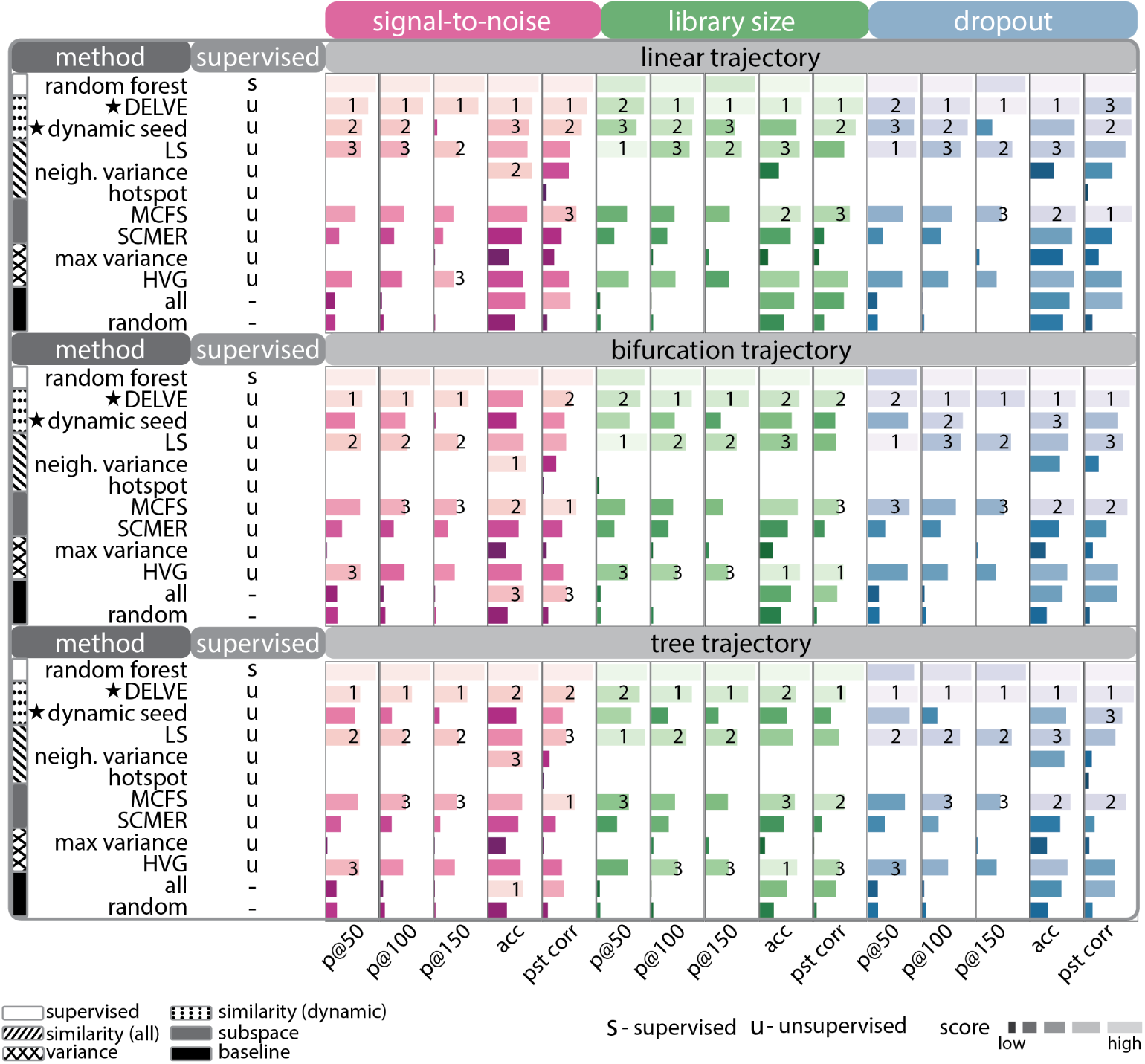
DELVE outperforms existing feature selection methods on representing trajectories in the presence of single-cell RNA sequencing noise. Feature selection methods were ranked by averaging their overall performance across datasets from different trajectory types (e.g. linear, bifurcation, tree) when subjected to noise corruption (e.g. decreased signal-to-noise ratio, decreased library size, and increased dropout). Several metrics were used to quantify trajectory preservation, including, precision of dynamically-expressed genes with 50 selected genes (*p*@50), precision at 100 selected genes (*p*@100), precision at 150 selected genes (*p*@150), *k*-NN classification accuracy of cell type labels (acc), and pseudotime correlation (pst). Here, higher-ranked methods are indicated by a longer lighter bar, and the star illustrates our approach (DELVE) as well as the performance from dynamic seed features of step 1 of the algorithm. DELVE often achieves the highest precision of lineage-specific differentially expressed genes, highest classification accuracy, and highest pseudotime correlation across noise conditions and trajectory types. Of note, random forest was included as a baseline representation to illustrate feature selection method performance when trained on ground truth cell type annotations; however, it was not ranked, as this study is focused on unsupervised feature selection method performance on trajectory preservation.

### Revealing molecular trajectories of proliferation and cell cycle arrest

Recent advances in spatial single-cell profiling technologies [8, 9, 84, 10, 11, 85, 86, 87, 88] have enabled the simultaneous measurement of transcriptomic or proteomic signatures of cells, while also retaining additional imaging or array-derived features that describe the spatial positioning or morphological properties of cells. These spatial single-cell modalities have provided fundamental insights into mammalian organogenesis [88, 89] and complex immune responses linked to disease progression [90, 91]. By leveraging imaging data to define cell-to-cell similarity, DELVE can identify smoothly varying spatial features that are strongly associated with cellular progression, such as changes in cell morphology or protein localization.

To demonstrate this, we applied DELVE to an integrated live cell imaging and protein iterative indirect immunofluorescence imaging (4i) dataset consisting of 2759 human retinal pigmented epithelial cells (RPE) undergoing the cell cycle (See *RPE analysis*). In a recent study [92], we performed time-lapse imaging on an asynchronous population of non-transformed RPE cells expressing a PCNA-mTurquoise2 reporter to record the cell cycle phase (G0/G1, S, G2, M) and age (time since last mitosis) of each cell. We then fixed the cells and profiled them with 4i to obtain measurements of 48 core cell cycle effectors. The resultant dataset consisted of 241 imaging-derived features describing the expression and localization of different protein markers (e.g. nucleus, cytoplasm, perinuclear region – denoted as ring), as well as morphological measurements from the images (e.g. size and shape of the nucleus). Given that time-lapse imaging was performed prior to cell fixation, this dataset provides the unique opportunity to rigorously evaluate feature selection methods on a real biological system (cell cycle) with technical challenges (e.g. many features with low signal-to-noise ratio, autofluorescence, sample degradation).

We first tested whether DELVE can identify a set of dynamically-expressed cell cycle-specific features to construct an approximate cellular trajectory graph for feature selection. Overall, we found that DELVE successfully identified dynamically-expressed seed features (*p* = 13) that are known to be associated with cell cycle proliferation (e.g. increase in DNA content and area of the nucleus) and captured key mechanisms previously shown to drive cell cycle progression (Fig. 4a right), including molecular events that regulate the G1/S and G2/M transitions. For example, the G1/S transition is governed by the phosphorylation of RB by cyclin:CDK complexes (e.g. cyclinA/CDK2 and cyclinE/CDK2), which control the expression of E2F transcription factors that regulate S phase genes [93]. We also observed an increase in expression of Skp2, which reduces p27-mediated inhibition of E2F1 target genes [94, 95]. In addition, our approach identified S phase events that are known to be associated with DNA replication, including an accumulation of PCNA foci at sites of active replication [96] and a DNA damage marker, pH2AX, which becomes phosphorylated in response to double-stranded DNA breaks in areas of stalled replication [97, 98]. Lastly, we observed an increase in expression of cyclin B localized to different regions of the cell, which is a primary regulator of G2/M transition alongside CDK1 [99, 100]. Of note, phosphorylation of RB also controls cell cycle re-entry and is an important biomarker that is often used for distinguishing proliferating from arrested cells [101, 102]. Furthermore, by ordering the average pairwise change in expression of features across ground truth phase annotations, we observed that DELVE dynamically-expressed seed features exhibited non-random patterns of expression variation that gradually increased throughout the canonical phases of the cell cycle (Fig. 4a), and were amongst the top ranked features that were biologically predictive of cell cycle phase and age measurements using a random forest classification and regression framework, respectively (See *Random forest*, Figure 4a right, Supplementary Fig. 13). Collectively, these results illustrate that the dynamic feature module identified by DELVE represents a minimum cell cycle feature set (Fig. 4b dynamic seed) that precisely distinguishes individual cells according to their cell cycle progression status and can be used to construct an approximate cellular trajectory for ranking feature importance.

**Figure 4:**
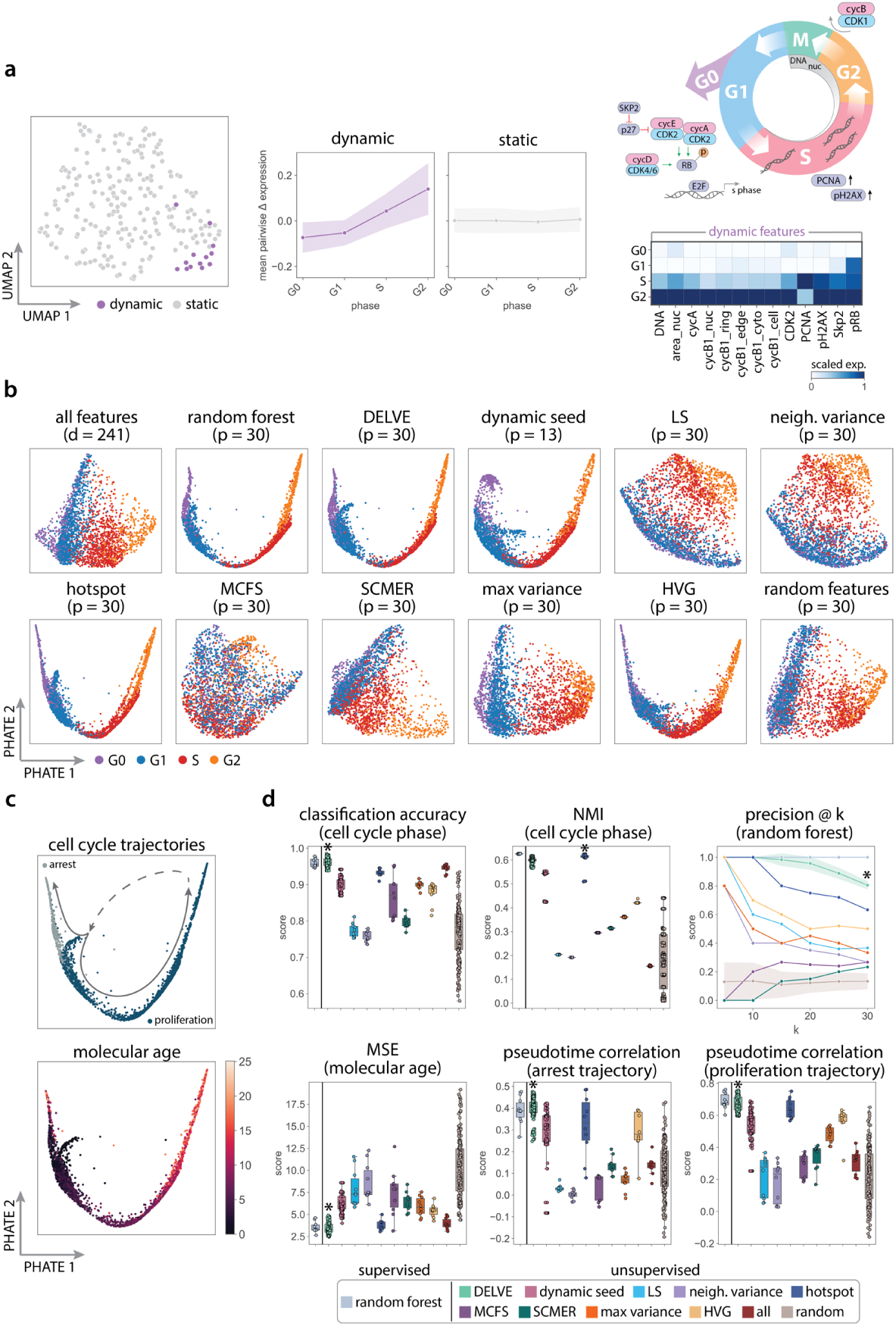
DELVE recovered signatures of proliferation and arrest in noisy protein immunofluorescence imaging data. (a) DELVE identified one dynamic module consisting of 13 seed features that represented a minimum cell cycle. (a left) UMAP visualization of image-derived features where each point indicates a dynamic or static feature identified by the model. (a middle) The average pairwise change in expression of features within DELVE modules ordered across ground truth cell cycle phase annotations. (a right top) Simplified signaling schematic of the cell cycle highlighting the role of DELVE dynamic seed features within cell cycle progression. (a right bottom) Heatmap of the standardized average expression of dynamic seed features across cell cycle phases. (b) Feature selection was performed to select the top (*p* = 30) ranked features from the original (*d* = 241) feature set according to a feature selection strategy. PHATE visualizations illustrating the overall quality of low-dimensional retinal pigmented epithelial cell cycle trajectories following feature selection. Cells were labeled according to ground truth cell cycle phase annotations from time-lapse imaging. Each panel represents a different feature selection strategy. (c) PHATE visualizations following DELVE feature selection, where cells were labeled according to cell cycle trajectory (top) or ground truth age measurements (bottom). (d) Performance of twelve feature selection methods on representing the cell cycle according to several metrics, including classification accuracy between predicted and ground truth phase annotations using a support vector machine classifier on selected features, normalized mutual information (NMI) between predicted and ground truth phase labels to indicate clustering performance, precision of phase-specific features determined by a random forest classifier trained on cell cycle phase annotations, mean-squared error between predicted and ground truth molecular age measurements using a support vector machine regression framework on selected features, and the correlation between estimated pseudotime to the ground truth molecular age measurements following trajectory inference on selected features. Error bands represent the standard deviation. * indicates the method with the highest median score. DELVE achieved the highest classification accuracy, highest p@k score, and high NMI clustering score indicating robust prediction of cell cycle phase. Furthermore, DELVE achieved the lowest mean squared error and highest correlation between arrest and proliferation estimated pseudotime and ground truth age measurements indicating robust prediction of cell cycle transitions.

We then comprehensively evaluated feature selection methods on their ability to retain imaging-derived features that define cell cycle phases and resolve proliferation and arrest cell cycle trajectories. We reasoned that cells in similar stages of the cell cycle (as defined by the cell cycle reporter) should have similar cell cycle signatures (4i features) and should be located near one another in a low dimensional projection. Fig. 4b shows the PHATE embeddings from each feature selection strategy. Using the DELVE feature set, we obtained a continuous PHATE trajectory structure that successfully captured the smooth progression of cells through the canonical phases of the cell cycle, where cells were tightly grouped together according to ground truth cell cycle phase annotations (Fig. 4b). Moreover, we observed that the two DELVE approaches (i.e. DELVE and dynamic seed), in addition to hotspot and HVG selection, produced qualitatively similar denoised lower-dimensional visualizations comparable to the supervised random forest approach that was trained on ground truth cell cycle phase annotations. In contrast, similarity-based approaches such as Laplacian score and neighborhood variance, which define a cell similarity graph according to all features, showed more diffuse presentations of cell states. Variance-based (max variance) or subspace-learning approaches (SCMER, MCFS) produced qualitatively similar embeddings to that produced using all features.

To quantitatively assess if selected features from a feature selection strategy were biologically predictive of cell cycle phases, we performed three complementary analyses. We first focused on the task of cell state classification, where our goal was to learn the ground truth cell cycle phase annotations from the selected feature set. To do so, we trained a support vector machine (SVM) classifier and compared the accuracy of predictions to their ground truth phase annotations (See *Support Vector Machine*). We performed nested 10-fold cross validation to obtain a distribution of predictions for each method. Overall, we found that DELVE achieved the highest median classification accuracy (DELVE = 0.960) obtaining a similar performance to the random forest classifier trained on cell cycle phase annotations (random forest = 0.957), and outperforming existing unsupervised approaches (e.g. hotspot = 0.935, max variance = 0.902, HVG = 0.889, MCFS = 0.870, SCMER = 0.797, LS = 0.770), as well as all features (0.946), suggesting that selected features with DELVE were more biologically predictive of cell cycle phases (Fig. 4d). We next aimed to assess how well a feature selection method could identify and rank cell cycle phase-specific features according to their representative power. To test this, we trained a random forest classifier on the ground truth phase annotations using nested 10-fold cross validation (See *Random forest*). We then compared the average ranked feature importance scores from the random forest to the selected features from a feature selection strategy using the precision@k metric. Strikingly, we found that DELVE achieved the highest median precision@k score (DELVE *p*@30 = 0.800) and appropriately ranked features according to their discriminative power of cell cycle phases despite being a completely unsupervised approach (Fig. 4d). This was followed by hotspot with a precision@k score of (hotspot *p*@30 = 0.633) and highly variable gene selection (HVG *p*@30 = 0.500). In contrast, the Laplacian Score and max variance obtained low precision scores (*p*@30 = 0.367 and 0.333 respectively), whereas neighborhood variance and subspace-learning feature selection methods MCFS and SCMER were unable to identify cell cycle phase-specific features from noisy 4i data and exhibited precision scores near random (*p*@30 = 0.267, 0.267, and 0.233, respectively). Lastly, we assessed if selected image-derived features could be used for downstream analysis tasks like unsupervised cell population discovery. To do so, we clustered cells using the KMeans++ algorithm [103] on the selected feature set and compared the predicted labels to the ground truth annotations using a normalized mutual information (NMI) score over 25 random initializations (See Unsupervised clustering). We found that hotspot, DELVE, and dynamic seed features were better able to cluster cells according to cell cycle phases and achieved considerably higher median NMI scores (0.615, 0.599, 0.543, respectively), as compared to retaining all features (0.155) (Fig. 4d). Moreover, we found that clustering performance was similar to that of the random forest trained on cell cycle phase annotations (0.626). In contrast, variance-based approaches achieved moderate NMI clustering scores (HVG: 0.421, max variance: 0.361) and alternative similarity-based and subspace learning approaches obtained low median NMI scores (*∼* 0.2) and were unable to cluster cells into biologically-cohesive cell populations. Of note, many trajectory inference methods use clusters when fitting trajectory models [24, 104, 105, 23], thus accurate cell-to-cluster assignments following feature selection is crucial for both cell type annotation and discovery, as well as for accurate downstream trajectory analysis interpretation. Collectively, these results highlight that feature selection with DELVE identifies imaging-derived features from noisy protein immunofluorescence imaging data that are more biologically predictive of cell cycle phases.

We then focused on the much harder task of predicting an individual cell’s progression through the cell cycle. A central challenge in trajectory inference is the destructive nature of single-cell technologies, where only a static snapshot of cell states is profiled. To move toward a quantitative evaluation of cell cycle trajectory reconstruction following feature selection, we leveraged the ground truth age measurements determined from time-lapse imaging of the RPE-PCNA reporter cell line. We first evaluated whether selected features could be used to accurately predict cell cycle age by training a support vector machine (SVM) regression framework using nested ten fold cross validation (See *Support Vector Machine*). Method performance was subsequently assessed by computing the mean squared error (MSE) between the predictions and the ground truth age measurements. Overall, we found that DELVE achieved the lowest median MSE (3.261), outperforming both supervised (random forest = 3.296) and unsupervised approaches (e.g. second best performer hotspot = 3.654) suggesting that selected features more accurately estimate the time following mitosis (Fig. 4c). Crucially, this highlights DELVE’s ability to learn new biologically-relevant features that might be missed when performing a supervised or unsupervised approach. Lastly, we assessed whether selected imaging features could be used to accurately infer proliferation and arrest cell cycle trajectories using common trajectory inference approaches (Fig. 4d). Briefly, we constructed predicted cell cycle trajectories using the diffusion pseudotime algorithm [83] under each feature selection strategy (See Trajectory inference and analysis). Cells were separated into proliferation or arrest lineages according to their average expression of pRB, and cellular progression was estimated using ten random root cells that had the youngest age. Feature selection method performance on trajectory inference was then quantitatively assessed by computing the Spearman rank correlation between estimated pseudotime and the ground truth age measurements. We found that DELVE achieved the highest median correlation of estimated pseudotime to the ground truth age measurements (proliferation: 0.656, arrest: 0.405) as compared to alternative methods (second best performer hotspot; proliferation: 0.632, arrest: 0.333) or all features (proliferation: 0.330, arrest: 0.135), indicating that our approach was better able to resolve both proliferation and cell cycle arrest trajectories where other approaches failed (Fig. 4d). Of note, DELVE was robust to the choice in hyperparameters and obtained reproducible results across a range of hyperparameter choices (See Supplementary Fig. 14).

As a secondary validation, we applied DELVE to 9 pancreatic adenocarcinoma (PDAC) cell lines (e.g. BxPC3, CFPAC, MiaPaCa, HPAC, Pa01C, Pa02C, PANC1, UM53) profiled with 4i (See *PDAC analysis*) and performed a similar evaluation of cell cycle phase and phase transition preservation (See Supplementary Figs. 16 - 24). Across all cell lines and metrics, we found that DELVE approaches and hotspot considerably outperformed alternative methods on recovering the cell cycle from noisy 4i data and often achieved the highest classification accuracy scores, clustering scores, and the highest correlation of cellular progression along proliferative and arrested cell cycle trajectories (See Supplementary Fig. 15). Notably, DELVE was particularly useful in resolving cell cycle trajectories from the PDAC cell lines that had numerous imaging measurements with low signal-to-noise ratio (e.g. CFPAC, MiaPaCa, PANC1, and UM53), whereas the alternative strategies were unable to resolve cell cycle phases and achieved scores near random (See Supplementary Figs. 17, 19, 23, 24).

### Identifying molecular drivers of CD8+ T cell effector and memory formation

To demonstrate the utility of our approach in a complex differentiation setting consisting of heterogeneous cell subtypes and shared and distinct molecular pathways, we applied DELVE to a single-cell RNA sequencing time series dataset consisting of 29,893 mouse splenic CD8+ T cells responding to acute viral infection [106]. Here, CD8+ T cells were profiled over 12 time points following infection with the Armstrong strain of lymphocytic choriomeningitis virus (LCMV): Naive, d3-, d4-, d5-, d6-, d7-, d10-, d14-, d21-, d32-, d60-, and d90-post-infection (See *CD8 T cell differentiation analysis*). During an immune response to acute viral infection, naive CD8+ T cells undergo a rapid activation and proliferation phase, giving rise to effector cells that can serve in a cytotoxic role to mediate immediate host defense, followed by a contraction phase giving rise to self-renewing memory cells that provide long-lasting protection and are maintained by antigen-dependent homeostatic proliferation [107, 108, 109]. Despite numerous studies detailing the molecular mechanisms of CD8+ T cell effector and memory fate specification, the molecular mechanisms driving activation, fate commitment, or T cell dysfunction continue to remain unclear due to the complex intra- and inter-temporal heterogeneity of the CD8+ T cell response during infection. Therefore, we applied DELVE to the CD8+ T cell dataset to resolve the differentiation trajectory and investigate transcriptional changes that are involved in effector and memory formation during acute viral infection with LCMV.

Following unsupervised seed selection, we found that DELVE successfully identified three gene modules constituting core regulatory complexes involved in CD8+ T cell viral response and had dynamic expression patterns that varied across experimental time following viral infection (Fig. 5a-c). Namely, dynamic module 0 contained genes involved in early activation and interferon response (e.g. Ly6a, Bst2, Ifi27l2a) [110, 111], and proliferation (e.g. Cenpa, Cenpf, Ccnb2, Ube2c, Top2a, Tubb4b, Birc5, Cks2, Cks1b, Nusap1, Hmgb2, Rrm2, H2afx, Pclaf, Stmn1) [112]. Dynamic module 1 contained genes involved in effector formation, including interferon-*γ* cytotoxic molecules, such as perforin/granzyme pathway (e.g. Gzma, Gzmk), integrins (e.g. Itgb1), killer cell lectin-like receptor family (e.g. Klrg1, Klrd1, Klrk1, Klrc1), chemokine receptors (e.g. Cxcr3, Cxcr6, Ccr2), and a canonical transcription factor involved in terminal effector formation (e.g. Id2) [113, 114, 115, 116]. Lastly, dynamic module 2 contained genes involved in long-term memory formation (e.g. Sell, Bcl2, Il7r, Ltb) [117, 118, 119, 120]. To quantitatively examine if genes within a dynamic module were meaningfully associated with one another, or had experimental evidence of co-regulation, we constructed gene association networks using experimentally-derived association scores from the STRING database [121]. Here, a permutation test was performed to assess the statistical significance of the observed experimental association amongst genes within a DELVE module as compared to random gene assignment (See Protein-protein interaction networks). Notably, across all three dynamic modules, DELVE identified groups of genes that had statistically significant experimental evidence of co-regulation (*p*-value = 0.001), where DELVE networks had a larger average degree of experimentally-derived edges than the null distribution (Fig. 5b: dynamic modules). Degree centrality is a simple measurement of the number of edges (e.g. experimentally derived associations between genes) connected to a node (e.g. gene); therefore, in this context, networks with a high average degree may contain complexes of genes that are essential for regulating a biological process. In contrast, genes identified by DELVE that exhibited random or noisy patterns of expression variation (static module) had little to no evidence of co-regulation (*p*-value = 1.0) and achieved a much lower average degree than networks defined by random gene assignment (Fig. 5b).

**Figure 5:**
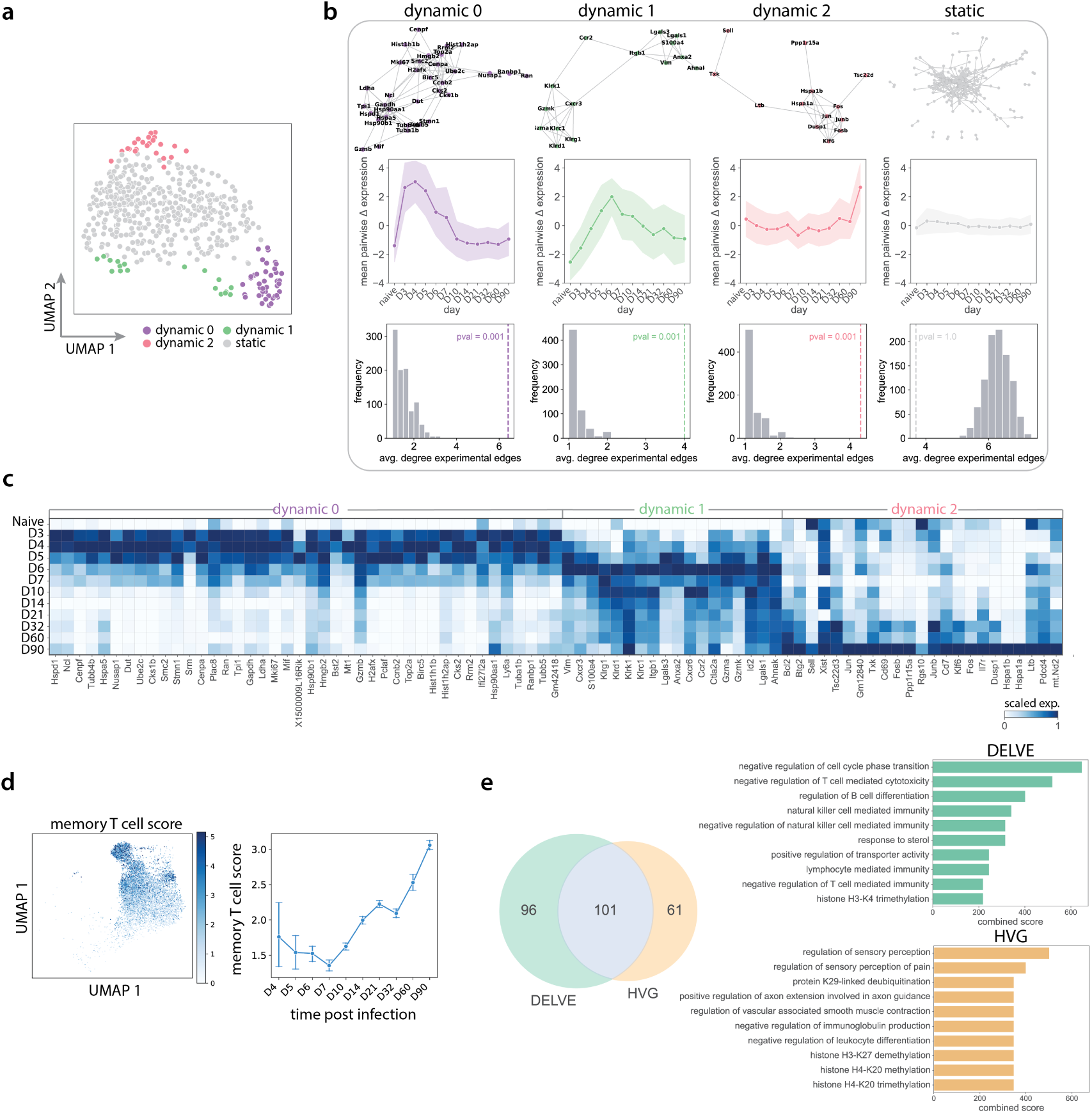
DELVE identified molecular drivers of CD8+ T cell effector and memory formation. (a) DELVE identified three dynamic modules representing cell cycle and early activation (dynamic 0), effector formation and cytokine signaling (dynamic 1), and long-term memory formation (dynamic 2) during CD8+ T cell differentiation response to viral infection with lymphocytic choriomeningitis virus (LCMV). UMAP visualization of (*d* = 500) genes where each point indicates a dynamic or static gene identified by the model. (b) A permutation test was performed using experimentally-derived association scores from the STRING protein-protein interaction database [121] to assess whether genes within DELVE dynamic modules had experimental evidence of co-regulation as compared to random assignment. (b top) STRING association networks, where nodes represent genes from a DELVE module and edges represent experimental evidence of association. (b middle) Average pairwise change in expression amongst genes within a module ordered by time following infection. (b bottom) Histograms showing the distribution of the average degree of experimentally-derived edges of gene networks from *R* = 1000 random permutations. The dotted line indicates the observed average degree from genes within a DELVE module. *p*-values were computed using a one-sided permutation test. (c) Heatmap visualization of the standardized average expression of dynamically-expressed genes identified by DELVE ordered across time following infection. (d) Trajectory inference was performed along the memory lineage using the diffusion pseudotime algorithm [83], where cell similarity was determined by DELVE selected genes or highly variable genes (HVG). UMAP visualization of memory T cell scores according to the average expression of known memory markers (Bcl-2, Sell, Il7r). Line plot indicates the onset of expression and cellular commitment to the memory lineage following infection. (e) Genes from the full dataset were regressed along estimated pseudotime using a generalized additive model to determine lineage-specific significant genes. The venn diagram illustrates the quantification and overlap of memory lineage-specific genes across feature selection strategies. The barplots show the top ten gene ontology terms associated with the temporally-expressed gene lists specific to each feature selection strategy.

Next, we examined if DELVE could be used to improve the identification of genes associated with long-term CD8+ T cell memory formation following trajectory inference. To do so, we first prioritized cells along the memory lineage by computing a memory T cell score according to the average expression of three known memory markers (Bcl2, Sell, and Il7r) (See *CD8 T cell differentiation analysis*, Fig. 5d). We then reconstructed the memory CD8+ T cell differentiation trajectory from middle-late stage cellular commitment using the diffusion pseudotime algorithm [83] on the top 250 ranked genes following DELVE feature selection or highly variable gene selection. Therefore, in this context, cell ordering was reflective of the differences in cell state along the memory lineage according to (1) dynamically-expressed genes that had experimental evidence of co-regulation or (2) variance-based selection. Lastly, we performed a regression analysis for each gene (*d* = 500) in the original dataset along estimated pseudotime using a negative binomial generalized additive model (GAM). Genes were considered to be differentially expressed along the memory lineage if they had a *q−*value *<* 0.05 following Benjamini-Hochberg false discovery rate correction [122] (See Trajectory inference and analysis). Overall, we found that ordering cells according to similarities in selected gene expression using DELVE was more reflective of long-term memory formation and achieved an increased recovery of memory lineage-specific genes, as directly compared to the standard approach of highly variable gene selection (Fig. 5e). To determine the biological relevance of these memory lineage-specific genes, we performed gene set enrichment analysis on the temporally-expressed genes specific to each feature selection strategy using EnrichR [123]. Here, DELVE obtained higher combined scores and identified more terms involved in immune regulation and memory CD8+ T cell formation, including negative regulation of cell cycle phase transition, negative regulation of T cell mediated cytotoxicity, lymphocyte mediated immunity, and negative regulation of T cell mediated immunity (Fig. 5e).

## Discussion

Computational trajectory inference methods have transformed our ability to study the continuum of cellular states associated with dynamic phenotypes; however, current approaches for reconstructing cellular trajectories can be hindered by biological or technical noise inherent to single-cell data [42, 43]. To mitigate the effect of unwanted sources of variation confounding trajectory inference, we designed a bottom-up unsupervised feature selection method that ranks and selects features that best approximate cell state transitions from dynamic feature modules that constitute core regulatory complexes. The key innovation of this work is the ability to parse temporally co-expressed features from noisy information-poor features prior to performing feature selection; in doing so, DELVE constructs cell similarity graphs that are more reflective of cell state progression for ranking feature importance.

In this study, we benchmarked twelve feature selection methods [46, 52, 25, 53, 55, 54, 50] on their ability to identify biologically relevant features for trajectory analysis from simulated single-cell RNA sequencing data where the ground truth was known. We found that DELVE achieved the highest recovery of differentially expressed genes within a cell type or along a cellular lineage, highest cell type classification accuracy, and most accurately estimated individual cell progression across a variety of trajectory topologies and biological or technical challenges. Furthermore, through a series or qualitative and quantitative comparisons, we illustrated how noise (e.g. stochasticity, sparsity, low library size) and information-poor features create spurious similarities amongst cells and considerably impact the performance of existing unsupervised similarity-based or subspace learning-based feature selection methods on identifying biologically-relevant features.

Next, we applied DELVE to a variety of biological contexts and demonstrated improved recovery of cellular trajectories over existing unsupervised feature selection strategies. In the context of studying the cell cycle from protein imaging-derived features [92], we illustrated how DELVE identified molecular features that were strongly associated with cell cycle progression and were more biologically predictive of cell cycle phase and age, as compared to the alternative unsupervised feature selection methods. Importantly, DELVE often achieved similar or better performance to the supervised Random Forest classification approach without the need for training on ground truth cell cycle labels. Lastly, in the context of studying heterogeneous CD8+ T cell response to viral infection from single-cell RNA sequencing data [106], we showed how DELVE identified gene complexes that had experimental evidence of co-regulation and were strongly associated with CD8+ T cell differentiation. Furthermore, we showed how performing feature selection with DELVE prior to performing trajectory inference improved the identification and resolution of gene programs associated with long-term memory formation that would have been missed by the standard unsupervised feature selection approach.

This study highlights how DELVE can be used to improve inference of cellular trajectories in the context of noisy single-cell omics data; however, it is important to note that feature selection can greatly bias the interpretation of the underlying cellular trajectory [42], thus careful consideration should be made when performing feature selection for trajectory analysis. Furthermore, we provided an unsupervised framework for ranking features according to their association with temporally co-expressed genes, although we note that DELVE can be improved by using a set of previously established regulators (See Step 2: feature ranking). Future work could focus on extending this framework for applications such as (1) deconvolving cellular trajectories using biological system-specific seed graphs or (2) studying complex biological systems such as organoid models or spatial microenvironments.

## Methods

### DELVE

DELVE identifies a subset of dynamically-changing features that preserve the local structure of the underlying cellular trajectory. In this section, we will (1) describe computational methods for the identification and ranking of features that have non-random patterns of dynamic variation, (2) explain DELVE’s relation to previous work, and (3) provide context for the mathematical foundations behind discarding information-poor features prior to performing trajectory inference.

#### Problem formulation

Let 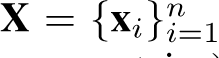 denote a single-cell dataset, where **x***_i_ ∈* R*^d^* represents the vector of *d* measured features (e.g. genes or proteins) measured in cell *i*. We assume that the data have an inherent trajectory structure, or biologically-meaningful ordering, that can be directly inferred by a limited subset of *p* features where *p ≪ d*. Therefore, our goal is to identify this limited set of *p* features from the original high-dimensional feature set that best approximate the transitions of cells through each stage of the underlying dynamic process.

#### Step 1: dynamic seed selection

##### Graph construction

Our approach DELVE extends previous similarity-based [52, 25, 53] or subspace-learning [55] feature selection methods by computing the dependence of each gene on the underlying cellular trajectory. In step 1, DELVE models cell states using a weighted k-nearest neighbor affinity graph of cells (*k* = 10), where nodes represent cells and edges describe the transcriptomic or proteomic similarity amongst cells according to the *d* profiled features encoded in **X**. More specifically, let *G* = (*V, E*) denote a between-cell affinity graph, where *V* represents the cells and the edges, *E*, are weighted according to a Gaussian kernel as,

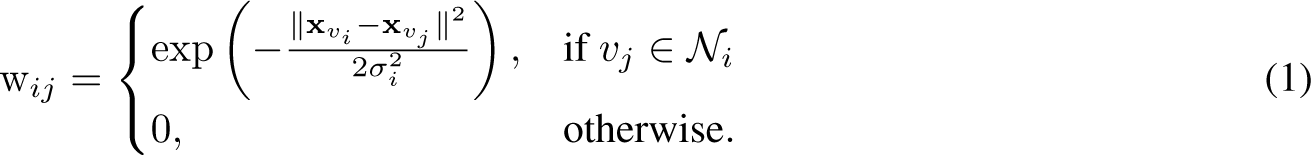

Here, **W** is a *n × n* between-cell similarity matrix, where cells *v_i_* and *v_j_* are connected with an edge with edge weight *w_ij_* if the cell *v_j_* is within the set of *v_i_’*s neighbors, as denoted by notation *N_i_*. Moreover, σ*_i_*, specific for a particular cell *i*, represents the Gaussian kernel bandwidth parameter that controls the decay of cell similarity edge weights. We chose a bandwidth parameter as the distance to the 3rd nearest neighbor as this has been shown previously in refs. [53] and [124] to provide reasonable decay in similarity weights.

##### Identification of feature modules

To identify groups of features with similar co-expression variation, DELVE clusters features according to changes in expression across prototypical cell neighborhoods. First, cellular neighborhoods are defined according to the average expression of each set of *k* nearest neighbors (*N_i_*) as, 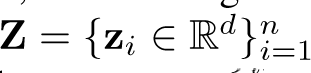, where each 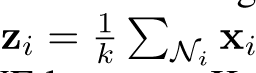 represents the center of the *k* nearest neighbors for cell i=1 i across all measured features.Next, DELVE leverages Kernel Herding sketching [66] to effectively sample m representative cell neighborhoods, or rows, from the per-cell neighbor averaged feature matrix, **Z**, as 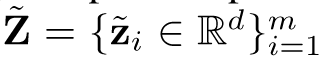. This sampling approach ensures that cellular neighborhoods are more reflective of the original distribution of cell states, while removing redundant cell states to aid in the scalability of estimating expression dynamics. DELVE then computes the average pairwise change in expression of features across representative cellular neighborhoods, **Δ**, as,

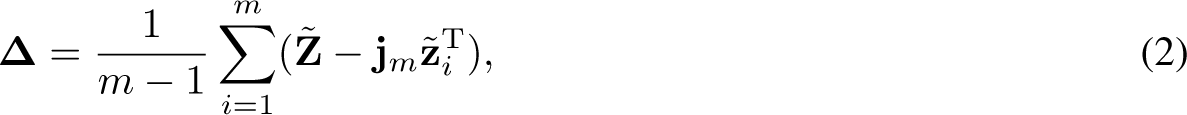

where **j**_*m*_ is a column vector of ones with length *m*, such that **j**_*m*_ ∈ R^*m*^. Lastly, features are clustered according to the transpose of their average pairwise change in expression across the representative cellular neighborhoods, **Δ**^T^, using the KMeans++ algorithm [103]. In this context, each DELVE module contains a set of features with similar local changes in co-variation across cell states along the cellular trajectory.

##### Dynamic expression variation permutation testing

To assess the significance of dynamic expression variation across grouped features within a DELVE module, we perform a permutation test as follows. Let 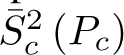 denote the average sample variance of the average pairwise change in expression across m cell neighborhoods for the set of *p* features (a set of features denoted as *P_c_*) within a DELVE cluster *c* as,

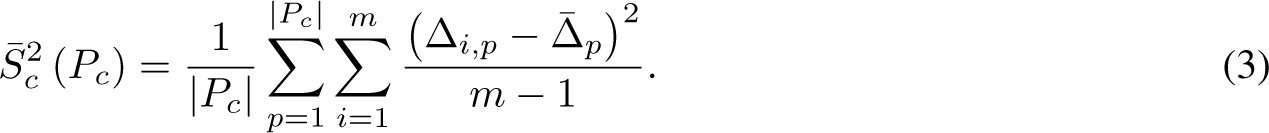

Moreover, let *R_q_* denote a set of randomly selected features sampled without replacement from the full feature space *d*, such that 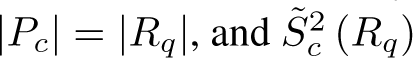 denote the average sample variance of randomly selected feature sets averaged across *t* random permutations as,

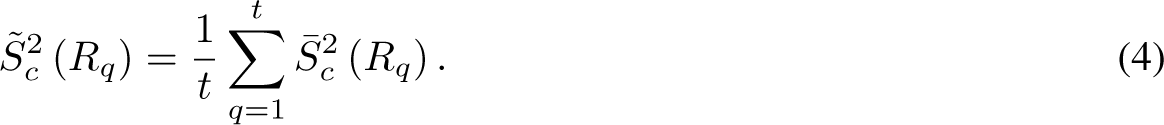

Here, DELVE considers a module of features as being dynamically-expressed if the average sample variance of the change in expression of the set of features within a DELVE cluster (or specifically feature set *P_c_*), is greater than random assignment, *R_q_*, across randomly permuted trials as,

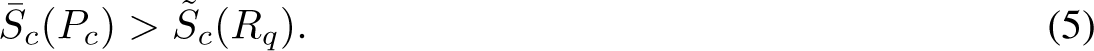

In doing so, this approach is able to identify and exclude modules of features that have static, random, or noisy patterns of expression variation, while retaining dynamically expressed features for ranking feature importance. Of note, given that KMeans++ clustering is used to initially assign features to a group, feature-to-cluster assignments can tend to vary due to algorithm stochasticity. Therefore, to reduce the variability and find a core set of features that are consistently dynamically-expressed, this process is repeated across ten random clustering initializations and the set of dynamically-expressed features are defined as the intersection across runs.

#### Step 2: feature ranking

Following dynamic seed selection, in step two, DELVE ranks features according to their association with the underlying cellular trajectory graph. First, DELVE approximates the underlying cellular trajectory by constructing a between-cell affinity graph, where the nodes represent the cells and edges are now re-weighted according to a Gaussian kernel between all cells based on the core subset of dynamically expressed regulators from step 1, such that 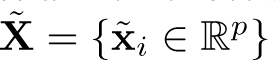 where *p ≪ d* as,

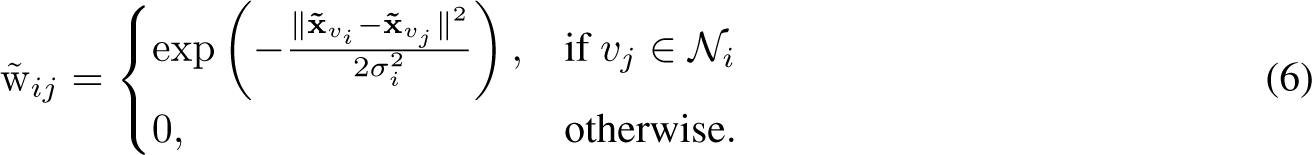

Here, *W̃* is a *n × n* between-cell similarity matrix, where cells *v_i_* and *v_j_* are connected with an edge with edge weight *w_ij_* if the cell is within the set of *v_i_*’s neighbors, denoted as **N_i_**. Moreover, as previously mentioned, *σ _i_* represents the Gaussian kernel bandwidth parameter for a particular cell *i* as the distance to the 3rd nearest neighbor.

Features are then ranked according to their association with the underlying cellular trajectory graph using graph signal processing techniques [52, 67, 68]. A graph signal *f* is any function that has a real defined value on all of the nodes, such that ***f*** ϵ R^*n*^ and *f_i_* gives the signal at the *i*th node. Intuitively, we can consider all features as graph signals and rank them according to their variation in expression along the approximate cell trajectory graph to see if they should be included or excluded from downstream analysis. Let **L** denote the unnormalized graph Laplacian, with **L** = **D** *−* **W̃**, where **D** is a diagonal degree matrix with each element as 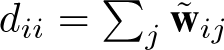. The local variation in the expression of feature signal **f** can then be defined as the weighted sum of differences in signals around a particular cell *i* as,

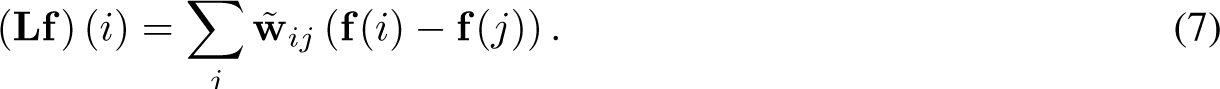

This metric effectively measures the similarity in expression of a particular node’s graph signal, denoted by the feature vector, **f**, around its *k* nearest neighbors. By summing the local variation in expression across all neighbors along the cellular trajectory, we can define the total variation in expression of feature graph signal **f** as,

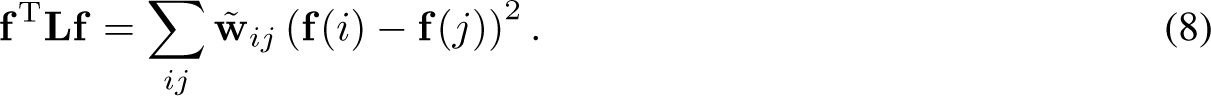

Otherwise known as the Laplacian quadratic form, in this context, the total variation represents the global smoothness of the particular graph signal encoded in **f** (e.g. expression of a particular gene or protein) along the approximate cellular trajectory graph. Intuitively, DELVE ultimately retains features that have a low total variation in expression, or have similar expression values amongst similar cells along the approximate cellular trajectory graph. In contrast, DELVE excludes features that have a high total variation in expression, or those which have expression values that are rapidly oscillating amongst neighboring cells, as these features are likely noisy or not involved in the underlying dynamic process that was initially seeded.

In this work, we ranked features according to their association with the cell-to-cell affinity graph defined by dynamically expressed features from DELVE dynamic modules using the Laplacian score [52]. This measure takes into account both the total variation in expression, as well as the overall global variance. For each of the original *d* measured features, or *graph signals* encoded in **f** with **f** *∈* R*^n^*, the Laplacian score *L_f_* is computed as,

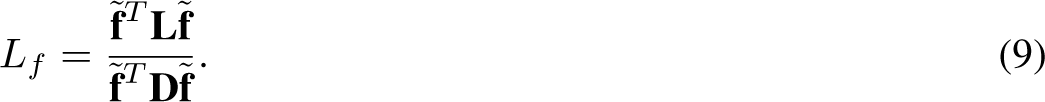

Here, **L** represents the unnormalized graph Laplacian, such that **L** = **D** *−* **W̃**, **D** is a diagonal degree matrix with the *i*th element of the diagonal *d_ii_* as 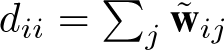, **f̃** represents the mean centered expression of feature **f** as 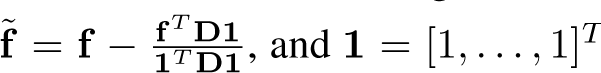. By sorting features in ascending order according to their Laplacian score, DELVE effectively ranks features that best preserve the local trajectory structure (e.g. an ideal numerator has a small local variation in expression along neighboring cells), as well as best preserve cell types (e.g. an ideal denominator has large variance in expression for discriminitive power).

### Benchmarked feature selection methods

In this section, we describe the twelve feature selection methods evaluated for representing biological trajectories. For more details on implementation and hyperparameters, see Supplementary Table 1.

#### Random forest

To quantitatively compare feature selection approaches on preserving biologically relevant genes or proteins, we aimed to implement an approach that would leverage ground truth cell type labels to determine feature importance. Random forest classification [46] is a supervised ensemble learning algorithm that uses an ensemble of decision trees to partition the feature space such that all of the samples (cells) with the same class (cell type labels) are grouped together. Each decision or split of a tree was chosen by minimizing the Gini impurity score as,

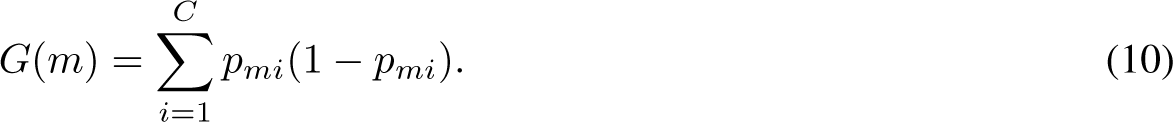

Here, *p_mi_* is the proportion of cells that belong to class *i* for a feature node *m*, and *C* is the total number of classes (e.g. cell types). We performed random forest classification using the sklearn v0.23.2 package in python. Nested 10-fold cross-validation was performed using stratified random sampling to assign cells to either a training or test set. The number of trees was tuned over a grid search within each fold prior to training the model. Feature importance scores were subsequently determined by the average Gini importance across folds.

#### Max variance

Max variance is an unsupervised feature selection approach that uses sample variance as a criterion for retaining discriminative features, where 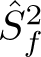 represents the sample variance for feature **f** *∈* R*^n^* as,

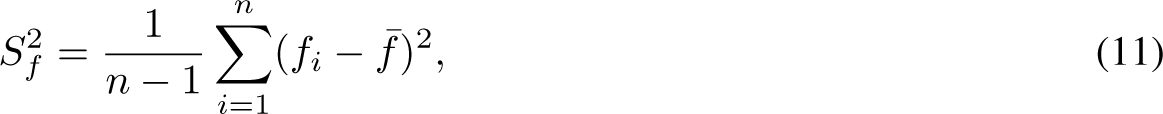

where *f_i_*indicates the expression value of feature **f** in cell *i*. We performed max variance feature selection by sorting features in descending order according to their variance score and selecting the top *p* maximally varying features.

#### Neighborhood variance

Neighborhood variance [25] is an unsupervised feature selection approach that uses a local neighborhood variance metric to select gradually-changing features for building biological trajectories. Namely, the neighborhood variance metric 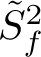 quantifies how much feature *f* varies across neighboring cells as,

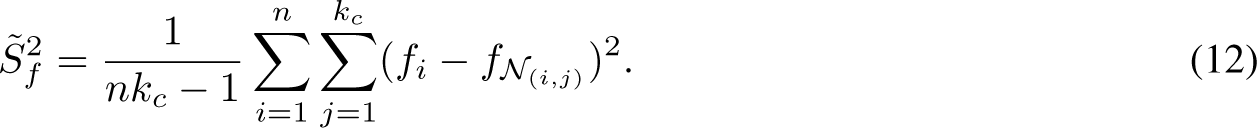

Here, *f_i_* represents the expression value of feature *f* for cell *i*, *N*_(*i,j*)_ indicates the *j* nearest neighbor of cell *i*, and *k_c_* is the minimum number of *k*-nearest neighbors required to form a fully connected graph. Features were subsequently selected if they had a smaller neighborhood variance 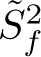 than global variance 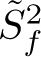,

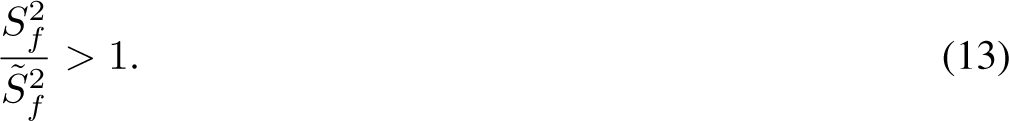

#### Highly variable genes

Highly variable gene selection [50] is an unsupervised feature selection approach that selects features according to a normalized dispersion measure. First, features are binned based on their average expression. Within each bin, genes are then z-score normalized to identify features that have a large variance, yet a similar mean expression. We selected the top *p* features using the highly variable genes function in Scanpy v1.8.1 (flavor = Seurat, bins = 20, n_top_genes = *p*).

#### Laplacian score

Laplacian score (LS) [52] is a locality-preserving unsupervised feature selection method that ranks features according to (1) how well a feature’s expression is consistent across neighboring cells defined by a between-cell similarity graph define by all profiled features and (2) the feature’s global variance. First, a weighted *k*-nearest neighbor affinity graph of cells (*k* = 10) is constructed according to pairwise Euclidean distances between cells based on all features, **X**. More specifically, let *G* = (*V, E*), where *V* represents the cells and edges, *E*, are weighted using a Gaussian as follows. Specifically, edge weights between cells *i* and *j* can be defined as,

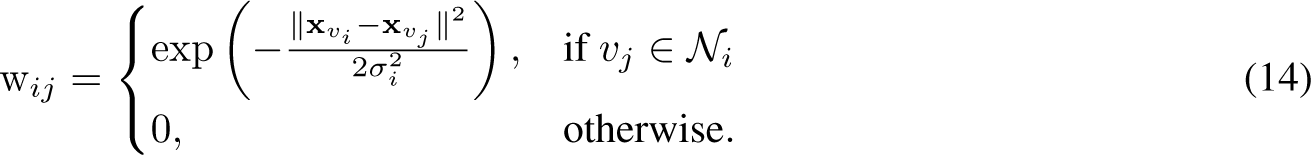

Here **W** is a *n × n* between-cell similarity matrix, where cells *v_i_* and *v_j_* are connected with an edge with edge weight *w_ij_* if the cell *v_j_* is within the set of *v_i_*’s neighbors, **N_i_**. Moreover, as previously described, *σ_i_* represents the bandwidth parameter for cell *i* defined as the distance to the 3rd nearest neighbor. For each feature **f**, where **f** *∈* R*^n^* represents the value of the feature across all *n* cells, we compute the Laplacian score, *L_f_* as,

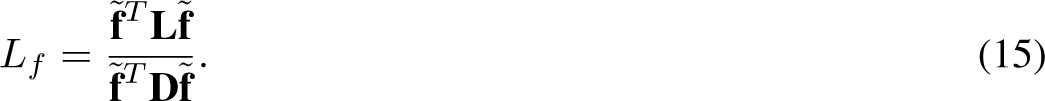

Here, **L** represents the unnormalized graph Laplacian, with **L** = **D** *−* **W**, **D** is a diagonal degree matrix with the *i*th element of the diagonal *d_ii_* as 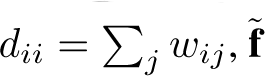 represents the mean centered expression of feature **f** as 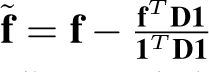, and **1** = [1, . . ., 1]*^T^* . We performed feature selection by sorting features in ascending order according to their Laplacian score and selecting the top *p* features.

#### MCFS

Multi-cluster feature selection (MCFS) [55] is an unsupervised feature selection method that selects for features that best preserve the multi-cluster structure of data by solving an L1 regularized least squares regression problem on the spectral embedding. Similar to the Laplacian score, first *k*-nearest neighbor affinity graph of cells (*k* = 10) is computed to encode the similarities in feature expression between cells *i* and *j* using a Gaussian kernel as,

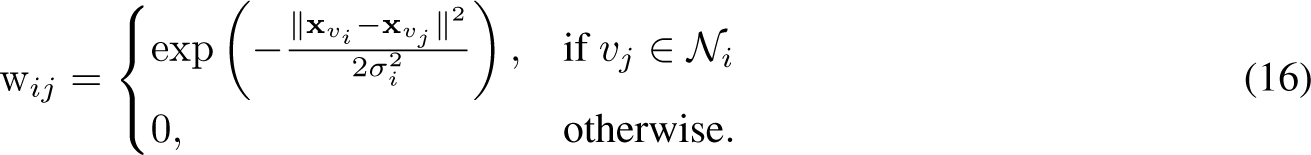

Similar to previous formulations above, **W** is an *n × n* between cell affinity matrix, where a pair of cells *v_i_* and *v_j_* are connected with an edge with weight *w_ij_* if cell *v_j_* is within the set of *v_i_*’s neighbors, **N_i_**. Further, *σ_i_*represents the kernel bandwidth parameter chosen to be the distance to the third nearest neighbor from cell *i*. Next, to represent the intrinsic dimensionality of the data, the spectral embedding [125] is computed through eigendecomposition of the unnormalized graph Laplacian **L**, where **L** = **D** *−* **W** as,

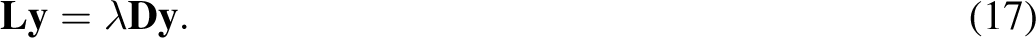

Here, 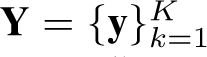 are the eigenvectors corresponding to the *K* smallest eigenvalues, **W** is a symmetric affinity matrix encoding cell similarity weights, and **D** represents a diagonal degree matrix with each element as *d_ii_* = ∑*_j_ w_ij_*. Given that eigenvectors of the graph Laplacian represent frequency harmonics [68] and low frequency eigenvectors are considered to capture the informative structure of the data, MCFS computes the importance of each feature along each intrinsic dimension **y***_k_* by finding a relevant subset of features by minimizing the error using an L1 norm penalty as,

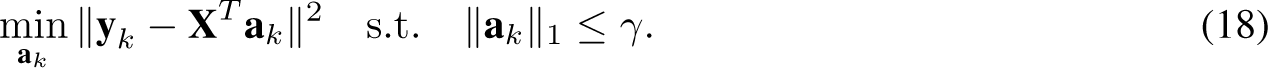

Here, the non-zero coefficients, **a***_k_*, indicate the most relevant features for distinguishing clusters from the embedding space, **y***_k_* and *γ* controls the sparsity and ensures the least relevant coefficients are shrunk to zero. The optimization is solved using the least angles regression algorithm [71], where for every feature, the MCFS score is defined as,

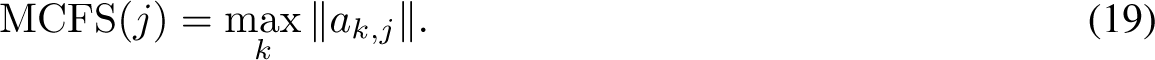

Here, *j* and *k* index feature and eigenvector, respectively. We performed multi-cluster feature selection with the number of eigenvectors *K* chosen to be the number of ground truth cell types present within the data, as this is the traditional convention in spectral clustering [57] and the number of nonzero coefficients was set to the number of selected features, *p*.

#### SCMER

Single-cell manifold-preserving feature selection (SCMER) [54] selects a subset of *p* features that represent the embedding structure of the data by learning a sparse weight vector **w** by formulating an elastic net regression problem that minimizes the KL divergence between a cell similarity matrix defined by all features and one defined by a reduced subset of features. More specifically, let **P** denote a between-cell pairwise similarity matrix defined in UMAP [56] computed with the full data matrix **X** *∈* R*^n×d^* and **Q** denote a between-cell pairwise similarity matrix defined in UMAP computed with the dataset following feature selection **Y** *∈* R*^n×p^*, where **Y** = **Xw** and *p ≪ d*. Here, elastic net regression is used to find a sparse and robust solution of **w** that minimizes the KL divergence as,

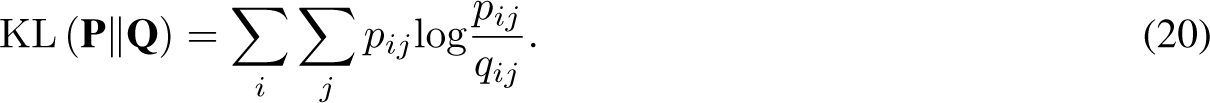

Features with non-zero weights in **w** are considered useful for preserving the embedding structure and selected for downstream analysis. We performed SCMER feature selection using the scmer v.0.1.0a3 package in python by constructing a *k*-nearest neighbor graph (*k* = 10) according to pairwise Euclidean distances of cells based on their first 50 principal components and using the default regression penalty weight parameters (lasso = 3.87*e −* 4, ridge = 0).

#### Hotspot

Hotspot [53] is an unsupervised gene module identification approach that performs feature selection through a test statistic that measures the association of a gene’s expression with the between-cell similarity graph defined based on the full feature matrix, **X**. More specifically, first, a *k*-nearest neighbor cell affinity graph (*k* = 10) is defined based on pairwise Euclidean distances between all pairs of cells using a Gaussian kernel as,

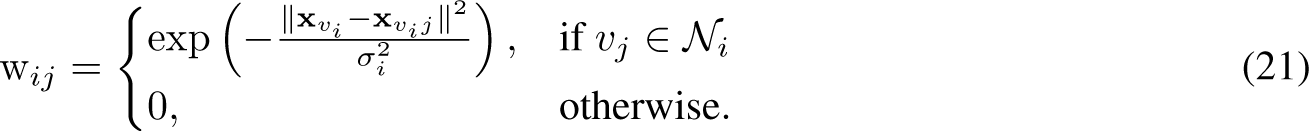

Here, cells *v_i_*and *v_j_* are connected with an edge with edge weight *w_ij_* if the cell *v_j_* is within the set of *v_i_*’s neighbors such that *_j_ w_ij_* = 1 for each cell and *σ_i_* represents the bandwidth parameter for cell *i* defined as the distance to the 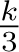 neighbor. For a given feature **f** *∈* R*^n^*, representing expression across all *n* cells where *f_i_* is the mean-centered and standardized expression of feature **f** in cell *i* according to a null distribution model of gene expression, the local autocorrelation test statistic representing the dependence of each gene on the graph structure is defined as,

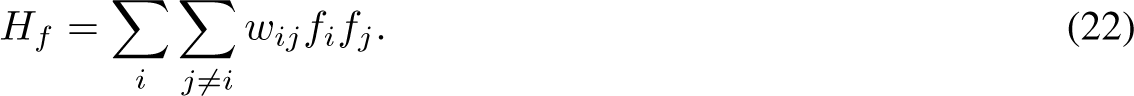

Hotspot was implemented using the hotspot v1.1.1 package in python, where we selected the top *p* features by sorting features in ascending order according to their significance with respect to a null model defined by a negative binomial distribution with the mean dependent on library size.

#### All features

To consider a baseline representation without feature selection, we evaluated performance using all features from each dataset following quality control preprocessing.

#### Random features

As a second baseline strategy, we simply selected a subset of random features without replacement. Results were computed across twenty random initializations for each dataset.

#### DELVE

DELVE was run as previously described. Here, we constructed a weighted *k*-nearest neighbor affinity graph of cells (*k* = 10), and 1000 neighborhoods were sketched to identify dynamic seed feature clusters (*c* = 3 for the simulated dataset, *c* = 5 for the RPE cell cycle dataset, *c* = 5 for the CD8 T cell differentiation dataset, and *c* = 10 for PDAC cell cycle datasets). Results were computed across twenty random initializations for each dataset.

### Datasets

We evaluated feature selection methods based on how well retained features could adequately recover biological trajectories under various noise conditions, biological contexts, and single-cell technologies.

#### Splatter simulation

Splatter [72] is a single-cell RNA sequencing simulation software that generates count data using a gamma-Poisson hierarchical model with modifications to alter the mean-variance relationship amongst genes, the library size, or sparsity. We used splatter to simulate a total of 90 ground truth datasets with different trajectory structures (e.g. linear, bifurcation, and tree topologies). First, we estimated simulation parameters by fitting the model to a real single-cell RNA sequencing dataset consisting of human pluripotent stem cells differentiating into mesoderm progenitors [126]. We then used the estimated parameters (mean_rate = 0.0173, mean_shape = 0.54, lib_loc = 12.6, lib_scale = 0.423, out_prob = 0.000342, out_fac_loc = 0.1, out_fac_scale = 0.4, bcv = 0.1, bcv_df = 90.2, dropout = None) to simulate a diverse set of ground truth reference trajectory datasets with the splatter paths function (python wrapper scprep SplatSimulate v1.1.0 of splatter v1.18.2). Here, a reference trajectory structure (e.g. bifurcation) was used to simulate linear and nonlinear changes in the mean expression of genes along each step of the specified differentiation path. We simulated ten differentiation datasets (1500 cells, 500 genes, 6 clusters) for each trajectory type (linear, bifurcation, tree) by modifying (1) the probability of a cell belonging to a cluster by randomly sampling from a Dirichelet distribution with six categories and a uniform concentration of one and (2) the path skew by randomly sampling from a beta distribution (*α* = 10, *β* = 10). The output of each simulation is a ground truth reference consisting of cell-to-cluster membership, differentially expressed genes per cluster or path, as well as a latent *step* vector that indicates the progression of each cell within a cluster. Lastly, we modified the step vector to be monotonically increasing across clusters within the specified differentiation path to obtain a reference pseudotime measurement.

To estimate how well feature selection methods can identify genes that represent cell populations and are differentially expressed along a differentiation path in noisy single-cell RNA sequencing data, we added relevant sources of biological and technical noise to the reference datasets.

1. *Biological Coefficient of Variation (BCV)*: To simulate the effect of stochastic gene expression, we modified the biological coefficient of variation parameter within splatter (BCV = 0.1, 0.25, 0.5). This scaling factor controls the mean-variance relationship between genes, where lowly expressed genes are more variable than highly expressed genes, following a *γ* distribution.
2. *Library size*: The total number of profiled mRNA transcripts per cell, or library size, can vary between cells within a single-cell RNA sequencing experiment and can influence the detection of differentially expressed genes [73], as well as impact the reproducibility of the lower-dimensional representation of the data [74]. To simulate the effect of differences in sequencing depth, we proportionally adjusted the gene means for each cell by modifying the location parameter (lib_loc = 12, 11, 10) of the log-normal distribution within splatter that estimates the library size scaling factors.
3. *Technical dropout*: Single-cell RNA sequencing data contain a large proportion of zeros, where only a small fraction of total transcripts are detected due to capture inefficiency and amplification noise [127]. To simulate the inefficient capture of mRNA molecules and account for the trend that lowly expressed genes are more likely to be affected by dropout, we undersampled mRNA counts by sampling from a binomial distribution with the scale parameter or dropout rate proportional to the mean expression of each gene as previously described in ref. [128] as,

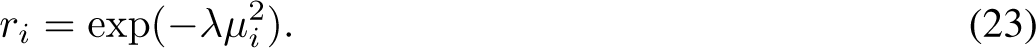

Here, *µ_i_* represents the log mean expression of gene *i*, and *λ* is a hyperparameter that controls the magnitude of dropout (*λ* = 0, 0.05, 0.1).

In our subsequent feature selection method analyses, we selected the top *p* = 100 features under each feature selection approach.

#### RPE analysis

The retinal pigmented epithelial (RPE) dataset [92] is an iterative indirect immunofluorescence imaging (4i) dataset consisting of RPE cells undergoing the cell cycle. Here, time-lapse imaging was performed on an asynchronous population of non-transformed human retinal pigmented epithelial cells expressing a PCNA-mTurquoise2 reporter in order to record the cell cycle phase (G0/G1, S, G2, M) and molecular age (time since last mitosis) of each cell. Following time-lapse imaging, cells were fixed and 48 core cell cycle effectors were profiled using 4i [8]. For preprocessing, we min-max normalized the data and performed batch effect correction on the replicates using ComBat [129]. Lastly, to refine phase annotations and distinguish G0 from G1 cells, we selected cycling G1 cells according to the bimodal distribution of 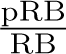 expression as described in ref. [92]. Of note, cells were excluded if they did not have ground truth phase or age annotations. The resultant dataset consisted of 2759 cells *×* 241 features describing the expression and localization of different protein markers. For our subsequent analysis, we selected the top *p* = 30 features for each feature selection approach.

#### PDAC analysis

The pancreatic ductal adenocarcinoma (PDAC) dataset is an iterative indirect immunofluorescence imaging dataset consisting of 9 human PDAC cell lines: BxPC3, CFPAC, HPAC, MiaPaCa, Pa01C, Pa02C, Pa16C, PANC1, UM53. For each cell line (e.g. BxPC3) under control conditions, we min-max normalized the data. Cell cycle phases (G0, G1, S, G2, M) were annotated *a priori* based on manual gating cells according to the abundance of core cell cycle markers. Phospho-RB (pRB) was used to distinguish proliferative cells (G1/S/G2/M, high pRB) from arrested cells (G0, low pRB). DNA content, E2F1, cyclin A (cycA), and phospho-p21 (p-p21) were used to distinguish G1 (DNA content = 2C, low cycA), S (DNA content = 2-4C, high E2F1), G2 (DNA content = 4C, high cycA), and M (DNA content = 4C, high p-p21). For our subsequent analysis, we selected the top *p* = 30 features for each feature selection approach.

#### CD8 T cell differentiation analysis

The CD8 T cell differentiation dataset [106] is a single-cell RNA sequencing dataset consisting of mouse splenic CD8+ T cells profiled over 12-time points (*d* = day) following infection with the Armstrong strain of the lymphocytic choriomeningitis virus: Naive, d3-, d4-, d5-, d6-, d7-, d10-, d14-, d21-, d32-, d60-, d90-post-infection. Spleen single-cell RNA sequencing data were accessed from the Gene Expression Omnibus using the accession code GSE131847 and concatenated into a single matrix. The dataset was subsequently quality control filtered according to the distribution of molecular counts. To remove dead or dying cells, we filtered cells that had more than twenty percent of their total reads mapped to mitochondrial transcripts. Genes that were observed in less than three cells or had less than 400 counts were also removed. Following cell and gene filtering, the data were transcripts-per-million normalized, log+1 transformed, and variance filtered using highly variable gene selection, such that the resulting dataset consisted of 29893 cells *×* 500 genes (See *Highly variable genes*). Lastly, to obtain lineage labels for trajectory analysis, cells were scored and gated according to their average expression of known memory markers (Bcl2, Sell, Il7r) using the score_genes function in Scanpy v1.8.1. When evaluating feature selection methods, we selected the top *p* = 250 features for each feature selection approach.

### Evaluation

#### Classification and regression

##### k-nearest neighbor classification

To quantitatively compare feature selection methods on retaining features that are representative of cell types, we aimed to implement an approach that would assess the quality of the graph structure. *k−*nearest neighbors classification is a supervised learning algorithm that classifies data based on labels of the *k*-most similar cells according to their gene or protein expression, where the output of this algorithm is a set of labels for every cell. We performed *k*-nearest neighbors classification to predict cell type labels from the simulated single-cell RNA sequencing datasets as follows. First, 3-fold cross-validation was performed using stratified random sampling to assign cells to either a training or a test set. Stratified random sampling was chosen to mitigate the effect of cell type class imbalance. Within each fold, feature selection was then performed on the training data to identify the top *p* = 250 relevant features according to a feature selection strategy. Next, a *k*-nearest neighbor classifier (*k* = 3) was fit on the feature selected training data to predict the cell type labels of the feature selected test query points. Here, labels are predicted as the mode of the cell type labels from the closest training data points according to Euclidean distance. Classification performance was subsequently assessed according to the median classification accuracy with respect to the ground truth cell type labels across folds.

##### Support Vector Machine

The Support Vector Machines (SVM) [130] is a supervised learning algorithm that constructs hyperplanes in the high-dimensional feature space to separate classes. We implemented SVM classification or regression using the sklearn v0.23.4 package in python. SVM classification was used to predict cell cycle phase labels for both RPE and PDAC 4i datasets, whereas regression was used to predict age measurements from time lapse imaging for the RPE dataset. Here, Nested 10-fold cross-validation was performed using stratified random sampling to assign cells to either a training set or a test set. Within each fold, feature selection was performed to identify the *p* most relevant features according to a feature selection strategy. SVM hyperparameters were then tuned over a grid search and phase labels were subsequently predicted from the test data according to those *p* features. Classification performance was assessed according to the median classification accuracy with respect to the ground truth cell type labels across folds. Regression performance was assessed according to the average mean squared error with respect to ground truth age measurements across folds.

#### Precision@k

To evaluate the biological relevance of selected features from each method, we computed precision@k (*p*@*k*) as the proportion of top *k* selected features that were considered to be biologically relevant according to a ground truth reference as,\

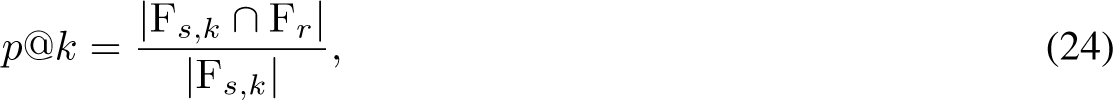

where F*_s,k_* indicates the set of selected features at threshold *k*, where F*_s,k_ ⊂* F*_s_*, and F*_r_* indicates the set of reference features. Reference features were defined as either (1) the ground truth differentially expressed features within a cluster or along a differentiation path from the single-cell RNA sequencing simulation study (see *Splatter simulation*) or (2) the features determined to be useful for classifying cells according to cell cycle phase using a random forest classifier trained on ground truth phase annotations from time-lapse imaging for the protein immunofluorescence imaging datasets (See *Random forest*, *RPE analysis*, *PDAC analysis*).

#### Unsupervised clustering

To evaluate feature selection method performance on retaining features that are informative for identifying canonical cell types, we performed unsupervised clustering on the data defined by the top *p* ranked features from a feature selection strategy. More specifically, for each feature selection approach, clustering was performed on the selected data using either the KMeans++ algorithm [103] with the number of centroids set as the same number of ground truth cell cycle phase labels for the protein immunofluorescence imaging datasets (RPE: *c* = 4, PDAC: *c* = 5).

To assess the accuracy of clustering assignments, we quantified a normalized mutual information (NMI) score between the predicted cluster labels and the ground truth cell type labels. Normalized mutual information [131] is a clustering quality metric that measures the amount of shared information between two cell-to-cluster partitions (**u** and **v**, such that the *i*th entry *u_i_* gives the cluster assignment of cell *i*) as,

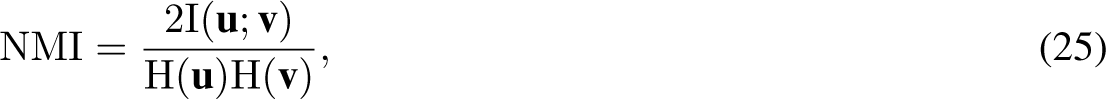

where, I(**u**; **v**) measures the mutual information between ground truth cell type labels **u** and cluster labels **v**, and H(**u**) or H(**v**) indicates the Shannon entropy or the amount of uncertainty for a given set of labels. Here, a score of 1 indicates that clustering on the selected features perfectly recovers the ground truth cell type labels. KMeans++ clustering was implemented using the KMeans function in sklearn v0.23.4.

#### Protein-protein interaction networks

In this work, we aimed to test whether features within DELVE dynamic clusters had experimental evidence of co-regulation as compared to random assignment. The STRING (search tool for the retrieval of interacting genes/proteins) database [121] is a relational database that computes protein association scores according to information derived from several *evidence* channels, including computational predictions (e.g. neighborhood, fusion, co-occurance), co-expression, experimental assays, pathway databases, and literature text mining. To assess the significance of protein interactions amongst features within a DELVE cluster, we performed a permutation test with a test statistic derived from STRING association scores using experimental evidence as follows.

Let *G_p_* = (*N_p_, E_p_*) denote a graph of *p* proteins from a DELVE cluster comprising the nodes *N_p_*, and *E_p_* denote the set of edges, where edge weights encode the association scores of experimentally-derived protein-protein interaction evidence from the STRING database. Moreover, let *G_r_* = (*N_r_, E_r_*) denote a graph of *r* proteins randomly sampled without replacement from the full feature space *d* such that *r* = *p* comprising the nodes *N_r_*, and *E_r_* denote the set of edges encoding the experimentally-derived association scores between those *r* proteins from the STRING database. We compute the permutation *p*-value as described previously in ref. [132] as,

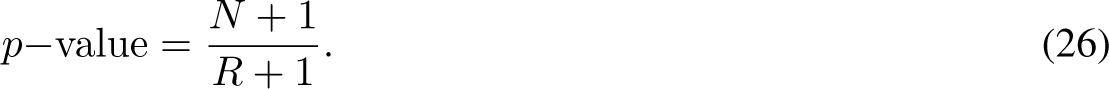

Here *N* indicates the number of times that *T*_r_ *≥ T*_obs_ out of *R* random permutations (*R* = 1000), where *T*_r_ is the average degree of a STRING association network from randomly permuted features as 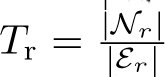, and *T*_obs_ is the average degree of a STRING association network from the features identified within a DELVE cluster as 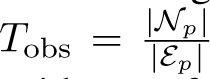. Of note, networks with higher degree are more connected, and thus show greater experimental evidence of protein-protein interactions. Experimental evidence-based association scores were obtained from the STRING database using the stringdb v0.1.5 package in python and networks were generated using networkx v2.5.1.

#### Trajectory inference and analysis

To evaluate how well feature selection methods could identify features that recapitulate the underlying cellular trajectory and can be used for trajectory analysis, we computed three metrics to assess trajectory preservation at different stages of inference: accuracy of the inferred trajectory graph, correlation of estimated pseudotime to the ground truth cell progression measurements, and the significance of dynamic features identified following regression analysis.

To obtain predicted trajectories, we performed trajectory inference using the diffusion pseudotime algorithm [83] based on 20 diffusion map components generated from a *k*-nearest neighbor graph (*k* = 10), where edge weights were determined by pairwise Euclidean distances between cells according to selected feature expression. Inference was performed for the following lineages – simulated trajectories: all cells, arrested trajectory: cells with G0 phase annotation, proliferative trajectory: cells with G1, S, G2, or M phase annotation, CD8 T cell memory lineage: cells following day 7 of infection with a memory score. Moreover, for each feature selection approach, we estimated pseudotime using ten random root cells according to *a priori* biological knowledge. Root cells were chosen as either (1) cells with the smallest ground truth pseudotime annotation for the simulated datasets, (2) cells with the youngest molecular age for 4i cell cycle datasets, or (3) cells from the day 7 population along the memory lineage for the CD8 T cell differentiation dataset. Feature selection trajectory performance was subsequently assessed as follows.

1. *Trajectory graph similarity:* Partition-based graph abstraction (PAGA) [104] performs trajectory inference by constructing a coarse grained trajectory graph of the data. First cell populations are determined either through unsupervised clustering, graph partitioning, or *a prior* experimental annotations. Next, a statistical measure of edge connectivity is computed between cell populations to estimate the confidence of a cell population transition. To assess if feature selection methods retain features that represent coarse cell type transitions, we compared predicted PAGA trajectory graphs to ground truth cell cycle reference trajectories curated from the literature [92]. First, PAGA connectivity was estimated between ground truth cell cycle phase groups using the *k*-nearest neighbor graph (*k* = 10) based on pairwise Euclidean distances between cells according to selected feature expression. We then computed the Jaccard distance between predicted and reference trajectories as,

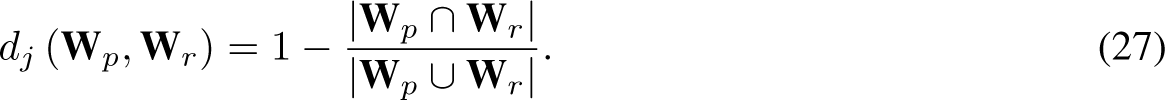

**W***_p_* indicates the predicted cell type transition adjacency matrix, where each entry W*_p,ij_* represents the connectivity strength between cell populations *i* and *j* from PAGA and **W***_r_* indicates the reference trajectory adjacency matrix with entries encoding ground truth cell type transitions curated from the literature. Here, a lower Jaccard distance indicates that predicted trajectories better capture known cellular transitions.

1. *Pseudotime correlation:* To evaluate if feature selection methods retain features that accurately represent a cell’s progression through a biological trajectory, we computed the Spearman rank correlation coefficient between estimated pseudotime following feature selection and ground truth cell progression annotations (e.g. the ground truth pseudotime labels generated from simulations, time-lapse imaging molecular age measurements).
2. *Regression analysis:* To identify genes associated with the inferred CD8+ T cell differentiation trajectory following feature selection, we performed regression analysis for each gene (*d* = 500) along estimated pseudotime using a negative binomial generalized additive model (GAM). Genes were considered to be differentially expressed along the memory lineage if they had a *q−*value *<* 0.05 following Benjamini-Hochberg false discovery rate correction [122].
3. *Gene Ontology:* To identify the biological relevance of the differentially expressed genes along inferred CD8+ T cell differentiation trajectories specific to each feature selection strategy, we performed gene set enrichment analysis on the set difference of significant genes from either highly variable gene selection or DELVE feature selection using Enrichr [123]. Here, we considered the mouse gene sets from GO Biological Process 2021.

Diffusion pseudotime was implemented using the dpt function in Scanpy v1.8.1, PAGA was implemented using the paga function in scanpy v1.8.1, GAM regression was implemented using the statsmodels v0.12.2 package in python, and gene set enrichment analysis was performed using the enrichr function in gseapy v1.0.4 package in python.

#### PHATE visualizations

To qualitatively compare lower dimensional representations from each feature selection strategy, we performed non-linear dimensionality reduction using PHATE (potential of heat-diffusion for affinity-based transition embedding) [75] as this approach performs reasonably well for representing complex continuous biological trajectories. PHATE was implemented using the phate v1.0.7 package in python. Here, we used the same set of hyperparameters across all feature selection strategies (knn = 30, t = 10, decay = 40).

#### Aggregate scores

To rank feature selection methods on preserving biological trajectories in the presence of single-cell noise, we computed rank aggregate scores by taking the mean of scaled method scores across simulated single-cell RNA sequencing datasets from a trajectory type and noise condition (e.g. linear trajectory, dropout noise). More specifically, we first defined an overall method score per dataset as the median of each metric. Method scores were subsequently min-max scaled to ensure datasets were equally weighted prior to computing the average.

## Data and code availability

The raw publicly available single-cell datasets used in this study are available in the Zenodo repository https://doi.org/10.5281/zenodo.4525425 for the RPE cell cycle dataset [133], the Zenodo repository https://doi.org/10.5281/zenodo.7860332 for the PDAC cell cycle datasets [134], and the Gene Expression Omnibus (GEO) under the accession code GSE131847 for the CD8 T cell differentiation dataset [135]. The preprocessed datasets are available in the Zenodo repository https://doi.org/10.5281/zenodo.7883604 [136]. DELVE is implemented as an open-source python package and is publicly available at https://github.com/jranek/ delve. Source code including all functions for benchmarking feature selection methods including preprocessing, feature selection, evaluation, and plotting are publicly available at: https://github.com/jranek/delve_benchmark.

## Funding

This work was supported by the National Institutes of Health, F31-HL156433 (JSR), 5T32-GM067553 (JSR), R01-GM138834 (JEP), NSF CAREER Award 1845796 (JEP), and NSF Award 2242980 (JEP).

### Authors’ contributions

JSR and WS conceptualized the study. JSR, NS, and JEP designed the method and computational analyses. JSR performed the data preprocessing, benchmarking, evaluation, and analysis. WS provided input for the RPE and PDAC cell cycle study. JM provided input for the CD8 T cell study. JSR wrote the manuscript with help from all authors. All authors read and approved of the final manuscript.

## Acknowledgements

We would like to thank Tarek Zikry, Garrett Sessions, and Alec Plotkin for their insightful discussions related to this work.

## Supplementary Information

### Supplementary Tables

**Supplementary Table 1:** Implemented feature selection method parameters

| Method name | Supervision | Parameter description | Parameters |
| --- | --- | --- | --- |
| Random Forest classifier | supervised | cell type labels<br>number of trees | cell type labels<br>n_estimators = 10, 100, 500 |
| DELVE | unsupervised | number of nearest neighbors<br>number of representative neighborhoods<br>number of modules | $k = 10$<br>$m = 1000$<br>$c = 3, 5, 10$ |
| Laplacian score | unsupervised | number of nearest neighbors<br>kernel bandwidth parameter | $k = 10$<br>$\sigma_i = 3\text{rd-nearest neighbor distance}$ |
| Neighborhood variance | unsupervised | NA | NA |
| Hotspot | unsupervised | number of nearest neighbors<br>kernel bandwidth parameter<br>number of principal components<br>model | $k = 10$<br>$\sigma_i = k/3\text{-nearest neighbor distance}$<br>n_pcs = 50<br>danb |
| MCFS | unsupervised | number of nearest neighbors<br>kernel bandwidth parameter<br>number of nonzero coefficients<br>number of eigenvectors | $k = 10$<br>$\sigma_i = 3\text{rd-nearest neighbor distance}$<br>$p$<br>$c = \text{number of known cell types}$ |
| SCMER | unsupervised | number of nearest neighbors<br>number of principal components<br>lasso regression parameter<br>ridge regression parameter | $k = 10$<br>n_pcs = 50<br>lasso = $3.87e - 4$<br>ridge = 0 |
| Max variance | unsupervised | NA | NA |
| Highly variable genes | unsupervised | flavor<br>bins<br>n_top_genes | Seurat<br>20<br>$p$ |
| All features | NA | NA | NA |
| Random features | NA | NA | NA |

### Supplementary Figures

**Supplementary Figure 1:**
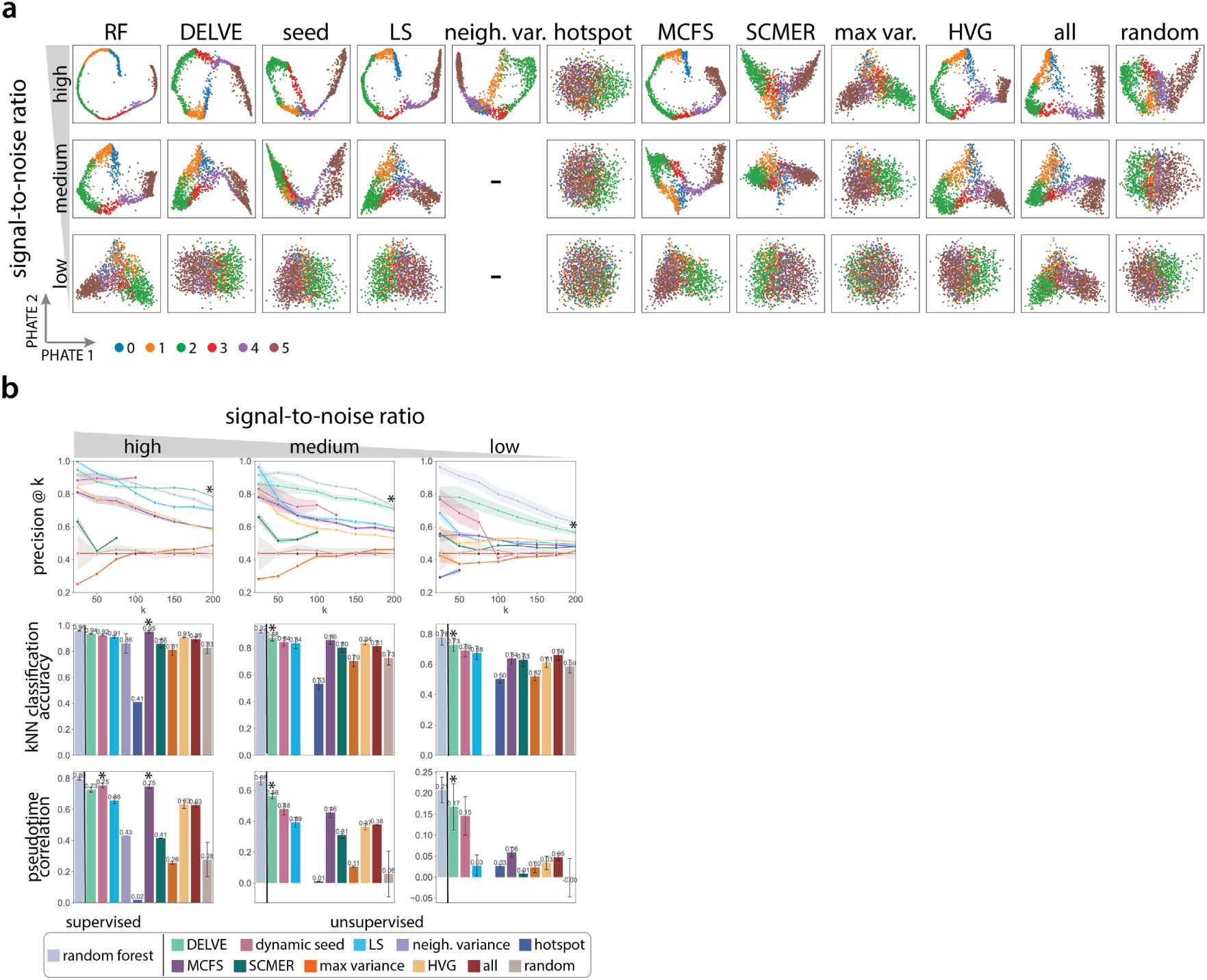
Comparison of feature selection methods on preserving linear differentiation trajectories under a reduction in the signal-to-noise ratio. (a) Example PHATE [75] visualizations of simulated linear differentiation trajectories for twelve feature selection strategies when subjected to a reduction in the signal-to-noise ratio (high, medium, low). The signal-to-noise ratio was altered by modifying the biological coefficient of variation parameter within Splatter (high: BCV = 0.1, medium: BCV = 0.25, low: BCV = 0.5). This scaling factor controls the mean-variance relationship between genes, where lowly expressed genes are more variable than highly expressed genes. *d* indicates the total number of genes (*d* = 500) and *p* indicates the number of selected genes following feature selection (*p* = 100). (b) Performance of twelve different feature selection methods when subjected to a reduction in the signal-to-noise ratio. Following feature selection, trajectory preservation was quantitatively assessed according to several metrics: the precision of differentially expressed genes at *k* selected genes (top), *k*-NN classification accuracy (middle), and pseudotime correlation (bottom) across 10 random trails. Error bars/ bands represent the standard deviation. * indicates the method with the highest median score. - indicates that the method identified no features.

**Supplementary Figure 2:**
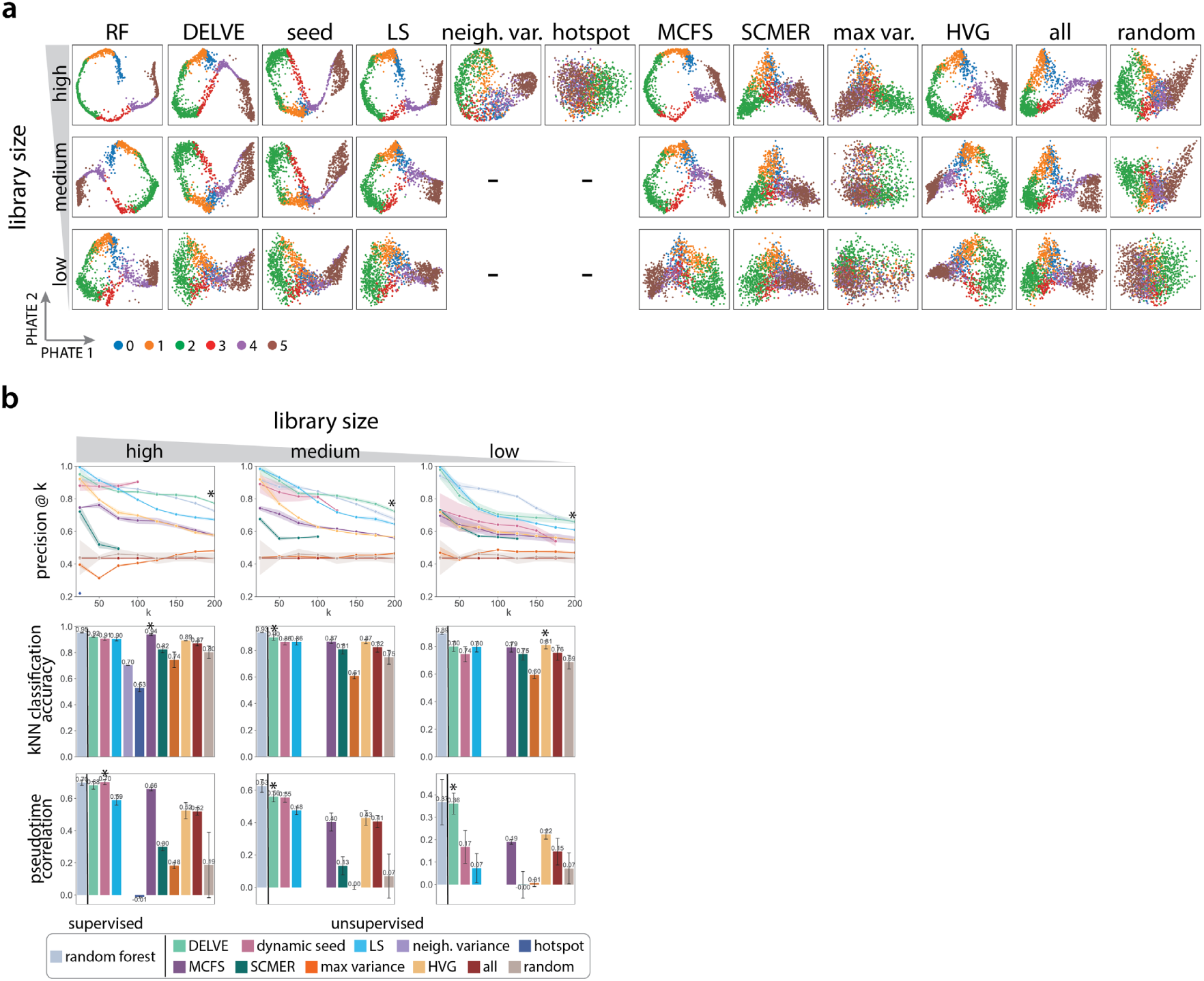
Comparison of feature selection methods on preserving linear differentiation trajectories in the presence of library size noise. (a) Example PHATE [75] visualizations of simulated linear differentiation trajectories for twelve feature selection strategies when subjected to a reduction in the total mRNA count (high, medium, low). Library size was reduced by modifying the location parameter in the log-normal distribution in Splatter [72] that specifies library size scaling factors (high: location = 12, medium: location = 11, low: location = 10). *d* indicates the total number of genes (*d* = 500) and *p* indicates the number of selected genes following feature selection (*p* = 100). (b) Performance of twelve different feature selection methods when subjected to a reduction in total mRNA count. Following feature selection, trajectory preservation was quantitatively assessed according to several metrics: the precision of differentially expressed genes at *k* selected genes (top), *k*-NN classification accuracy (middle), and pseudotime correlation (bottom) across 10 random trails. Error bars/ bands represent the standard deviation. * indicates the method with the highest median score. - indicates that the method identified no features.

**Supplementary Figure 3:**
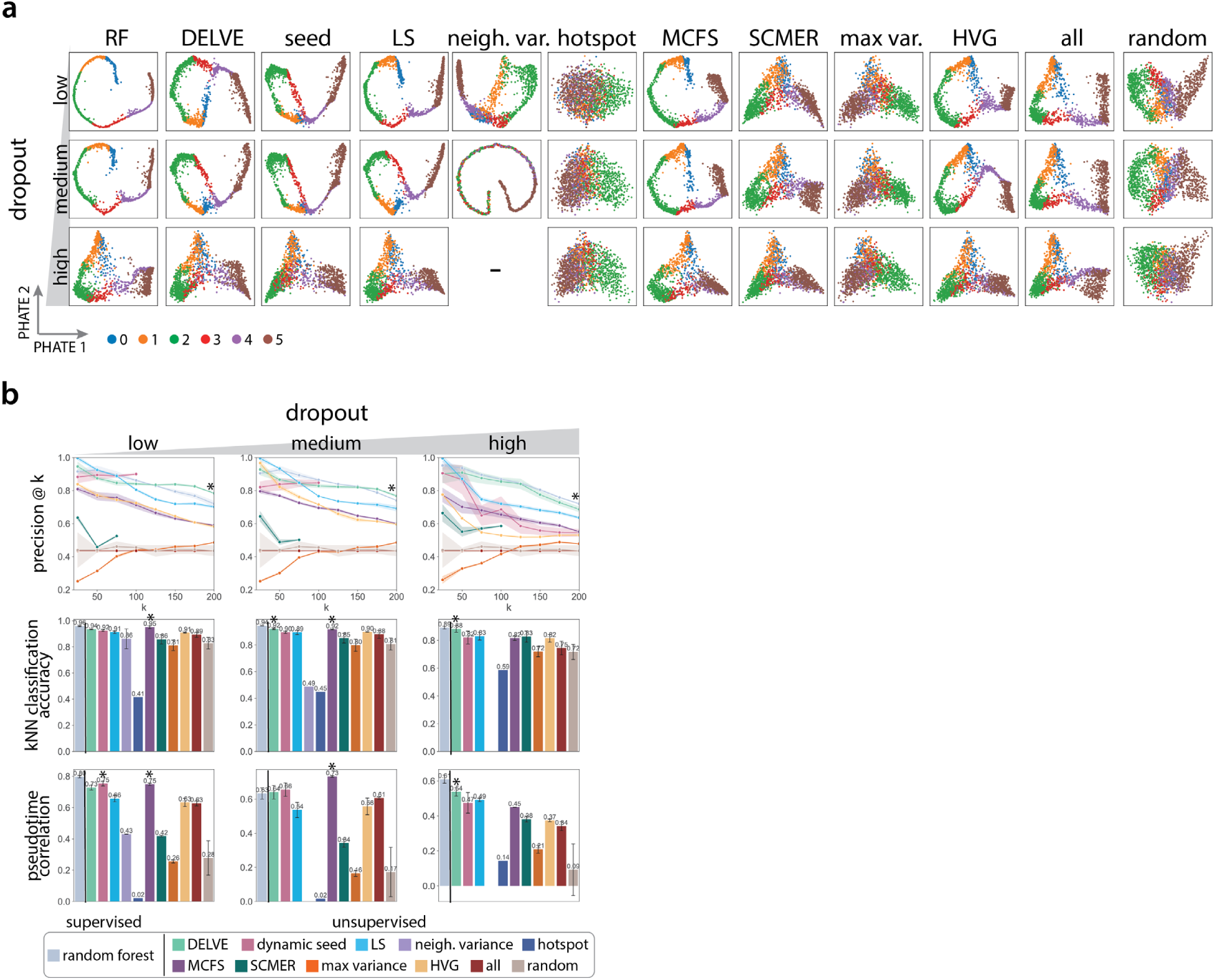
Comparison of feature selection methods on preserving linear differentiation trajectories is the presence of dropout noise. (a) Example PHATE [75] visualizations of simulated linear differentiation trajectories for twelve feature selection strategies when subjected to an increase in the amount of dropout (low, medium, high). Technical dropout was simulated by undersampling mRNA counts by sampling from a binomial distribution with the scale parameter or dropout rate proportional to the mean expression of each gene (low: *λ* = 0, medium: *λ* = 0.05, low: *λ* = 0.1). *d* indicates the total number of genes (*d* = 500) and *p* indicates the number of selected genes following feature selection (*p* = 100). (b) Performance of twelve different feature selection methods when subjected to an increase in the amount of dropout noise. Following feature selection, trajectory preservation was quantitatively assessed according to several metrics: the precision of differentially expressed genes at *k* selected genes (top), *k*-NN classification accuracy (middle), and pseudotime correlation (bottom) across 10 random trails. Error bars/ bands represent the standard deviation. * indicates the method with the highest median score. - indicates that the method identified no features.

**Supplementary Figure 4:**
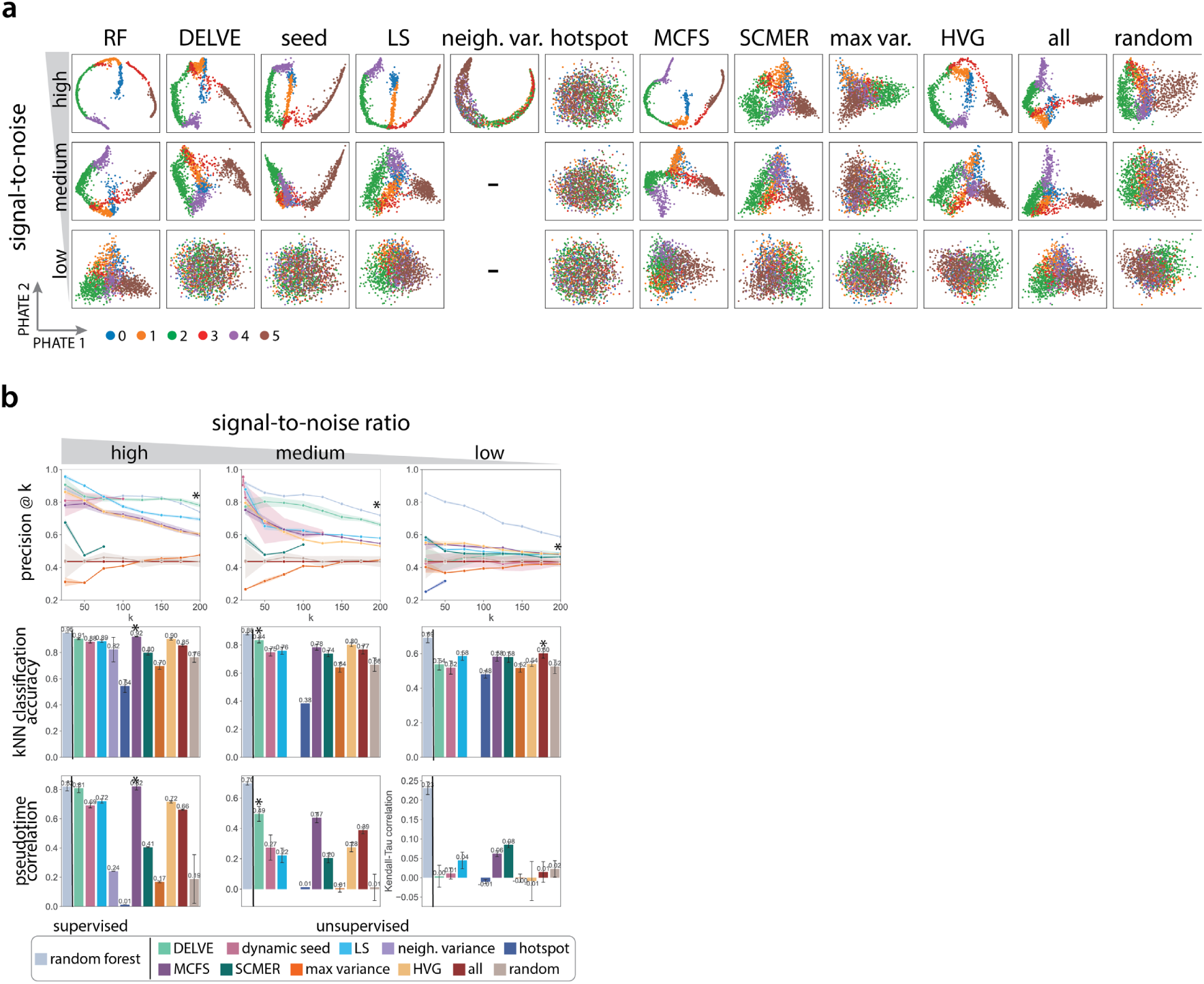
Comparison of feature selection methods on preserving bifurcation differentiation trajectories under a reduction in the signal-to-noise ratio. (a) Example PHATE [75] visualizations of simulated bifurcation differentiation trajectories for twelve feature selection strategies when subjected to a reduction in the signal-to-noise ratio (high, medium, low). The signal-to-noise ratio was altered by modifying the biological coefficient of variation parameter within Splatter (high: BCV = 0.1, medium: BCV = 0.25, low: BCV = 0.5). This scaling factor controls the mean-variance relationship between genes, where lowly expressed genes are more variable than highly expressed genes. *d* indicates the total number of genes (*d* = 500) and *p* indicates the number of selected genes following feature selection (*p* = 100). (b) Performance of twelve different feature selection methods when subjected to a reduction in the signal-to-noise ratio. Following feature selection, trajectory preservation was quantitatively assessed according to several metrics: the precision of differentially expressed genes at *k* selected genes (top), *k*-NN classification accuracy (middle), and pseudotime correlation (bottom) across 10 random trails. Error bars/ bands represent the standard deviation. * indicates the method with the highest median score. - indicates that the method identified no features.

**Supplementary Figure 5:**
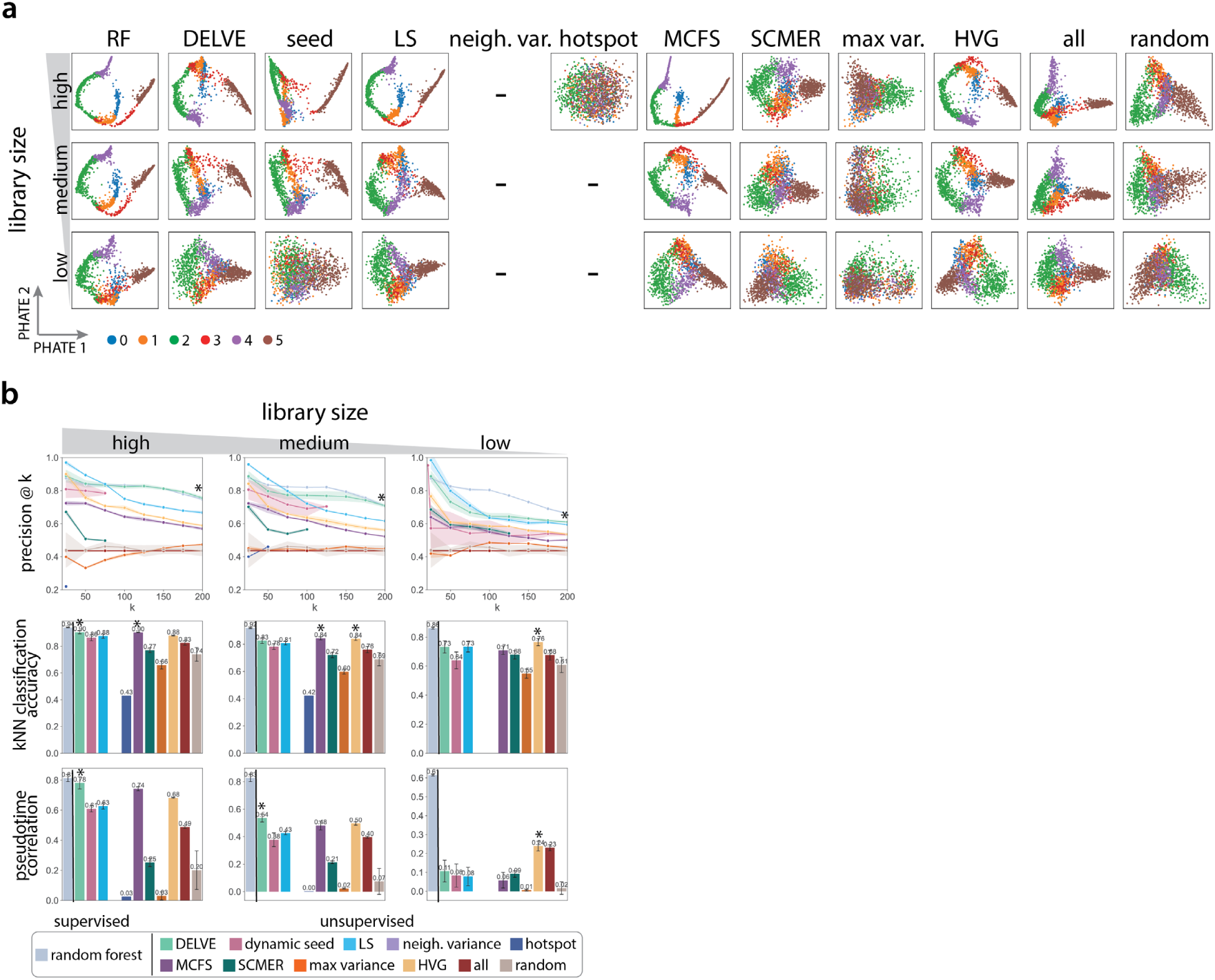
Comparison of feature selection methods on preserving bifurcation differentiation trajectories in the presence of library size noise. (a) Example PHATE [75] visualizations of simulated bifurcation differentiation trajectories for twelve feature selection strategies when subjected to a reduction in the total mRNA count (high, medium, low). Library size was reduced by modifying the location parameter in the log-normal distribution in Splatter [72] that specifies library size scaling factors (high: location = 12, medium: location = 11, low: location = 10). *d* indicates the total number of genes (*d* = 500) and *p* indicates the number of selected genes following feature selection (*p* = 100). (b) Performance of twelve different feature selection methods when subjected to a reduction in total mRNA count. Following feature selection, trajectory preservation was quantitatively assessed according to several metrics: the precision of differentially expressed genes at *k* selected genes (top), *k*-NN classification accuracy (middle), and pseudotime correlation (bottom) across 10 random trails. Error bars/ bands represent the standard deviation. * indicates the method with the highest median score. - indicates that the method identified no features.

**Supplementary Figure 6:**
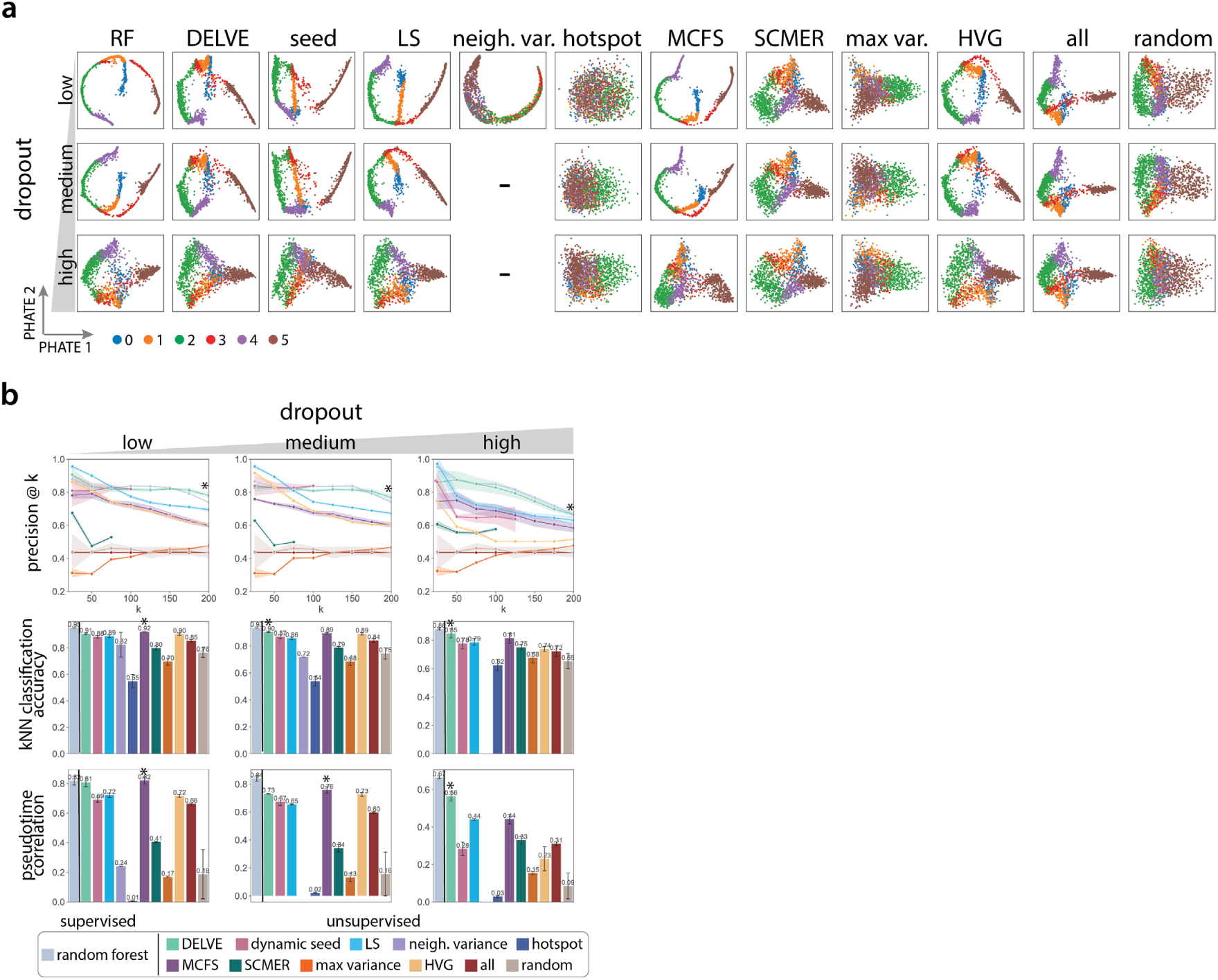
Comparison of feature selection methods on preserving bifurcation differentiation trajectories is the presence of dropout noise. (a) Example PHATE [75] visualizations of simulated bifurcation differentiation trajectories for twelve feature selection strategies when subjected to an increase in the amount of dropout (low, medium, high). Technical dropout was simulated by undersampling mRNA counts by sampling from a binomial distribution with the scale parameter or dropout rate proportional to the mean expression of each gene (low: *λ* = 0, medium: *λ* = 0.05, low: *λ* = 0.1). *d* indicates the total number of genes (*d* = 500) and *p* indicates the number of selected genes following feature selection (*p* = 100). (b) Performance of twelve different feature selection methods when subjected to an increase in the amount of dropout noise. Following feature selection, trajectory preservation was quantitatively assessed according to several metrics: the precision of differentially expressed genes at *k* selected genes (top), *k*-NN classification accuracy (middle), and pseudotime correlation (bottom) across 10 random trails. Error bars/ bands represent the standard deviation. * indicates the method with the highest median score. - indicates that the method identified no features.

**Supplementary Figure 7:**
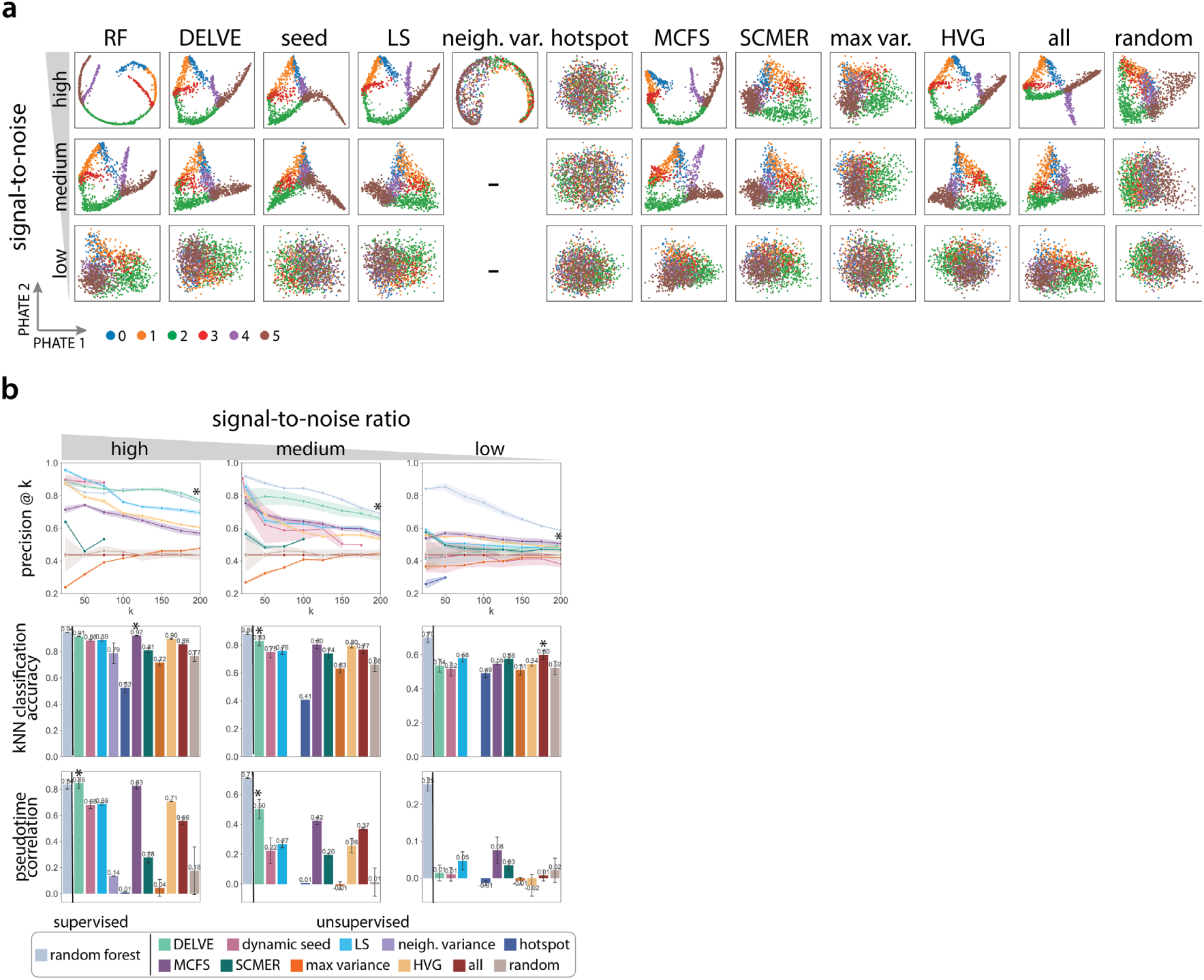
Comparison of feature selection methods on preserving tree differentiation trajectories under a reduction in the signal-to-noise ratio. (a) Example PHATE [75] visualizations of simulated tree differentiation trajectories for twelve feature selection strategies when subjected to a reduction in the signal-to-noise ratio (high, medium, low). The signal-to-noise ratio was altered by modifying the biological coefficient of variation parameter within Splatter (high: BCV = 0.1, medium: BCV = 0.25, low: BCV = 0.5). This scaling factor controls the mean-variance relationship between genes, where lowly expressed genes are more variable than highly expressed genes. *d* indicates the total number of genes (*d* = 500) and *p* indicates the number of selected genes following feature selection (*p* = 100). (b) Performance of twelve different feature selection methods when subjected to a reduction in the signal-to-noise ratio. Following feature selection, trajectory preservation was quantitatively assessed according to several metrics: the precision of differentially expressed genes at *k* selected genes (top), *k*-NN classification accuracy (middle), and pseudotime correlation (bottom) across 10 random trails. Error bars/ bands represent the standard deviation. * indicates the method with the highest median score. - indicates that the method identified no features.

**Supplementary Figure 8:**
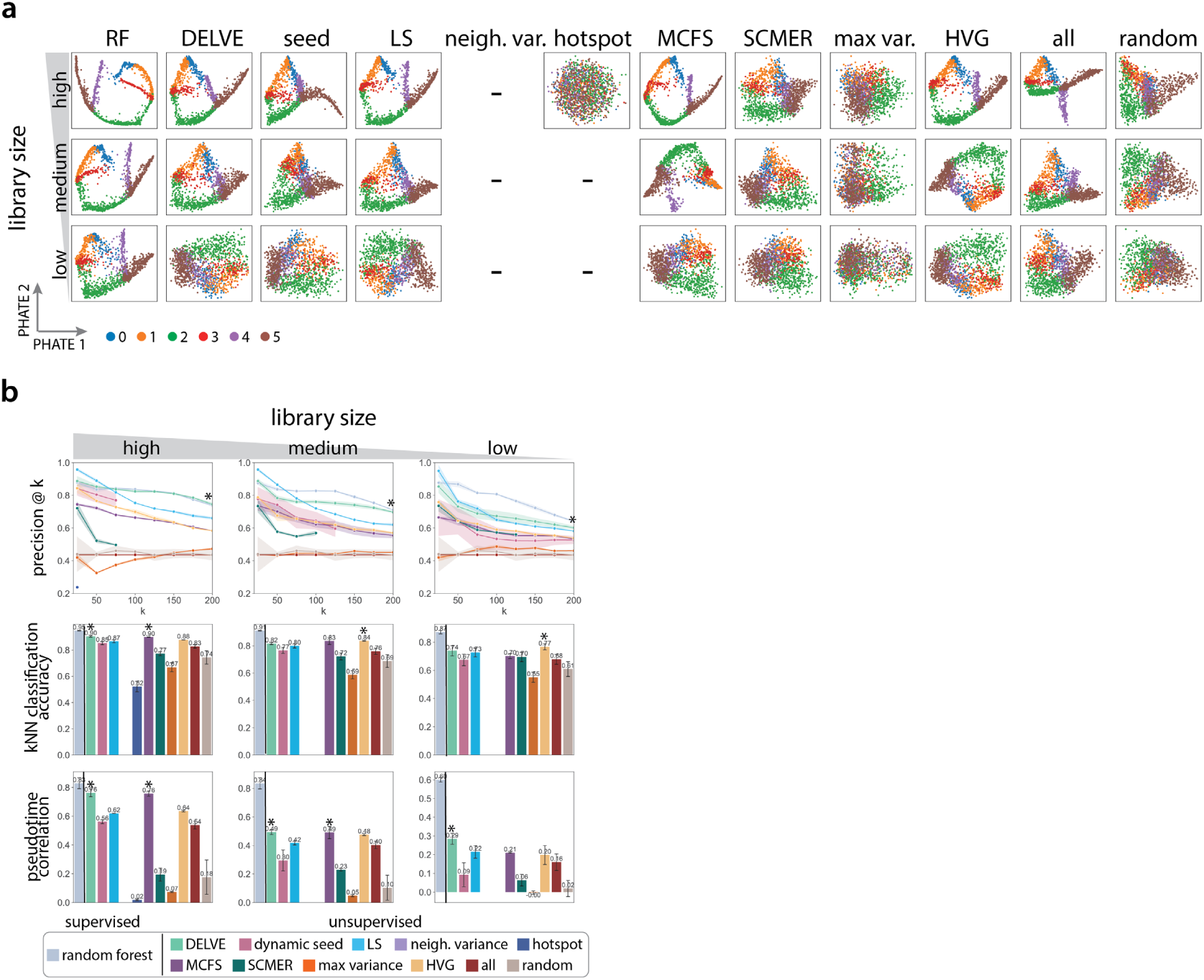
Comparison of feature selection methods on preserving tree differentiation trajectories in the presence of library size noise. (a) Example PHATE [75] visualizations of simulated tree differentiation trajectories for twelve feature selection strategies when subjected to a reduction in the total mRNA count (high, medium, low). Library size was reduced by modifying the location parameter in the log-normal distribution in Splatter [72] that specifies library size scaling factors (high: location = 12, medium: location = 11, low: location = 10). *d* indicates the total number of genes (*d* = 500) and *p* indicates the number of selected genes following feature selection (*p* = 100). (b) Performance of twelve different feature selection methods when subjected to a reduction in total mRNA count. Following feature selection, trajectory preservation was quantitatively assessed according to several metrics: the precision of differentially expressed genes at *k* selected genes (top), *k*-NN classification accuracy (middle), and pseudotime correlation (bottom) across 10 random trails. Error bars/ bands represent the standard deviation. * indicates the method with the highest median score. - indicates that the method identified no features.

**Supplementary Figure 9:**
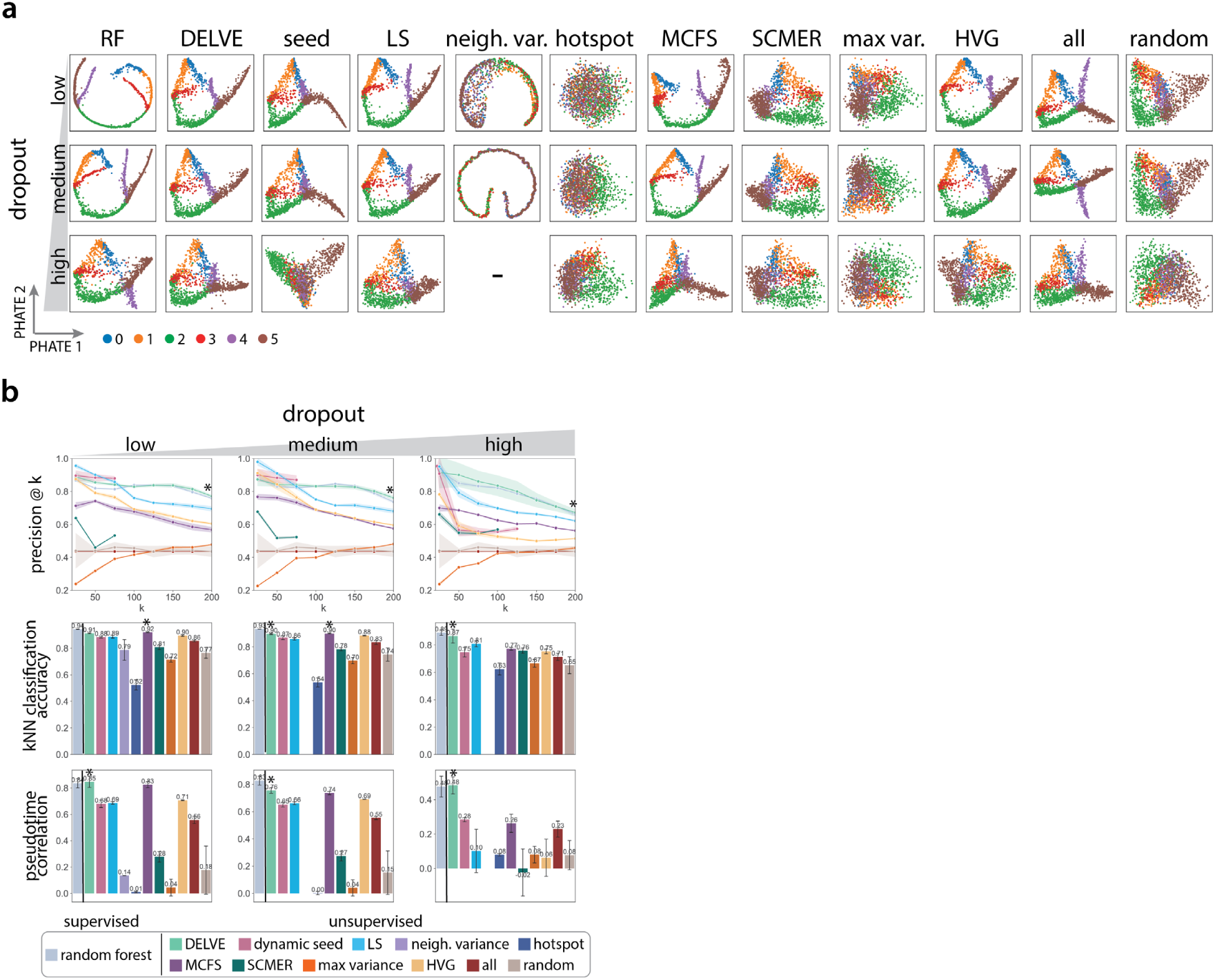
Comparison of feature selection methods on preserving tree differentiation trajec-tories is the presence of dropout noise. (a) Example PHATE [75] visualizations of simulated tree differentiation trajectories for twelve feature selection strategies when subjected to an increase in the amount of dropout (low, medium, high). Technical dropout was simulated by undersampling mRNA counts by sampling from a binomial distribution with the scale parameter or dropout rate proportional to the mean expression of each gene (low: *λ* = 0, medium: *λ* = 0.05, low: *λ* = 0.1). *d* indicates the total number of genes (*d* = 500) and *p* indicates the number of selected genes following feature selection (*p* = 100). (b) Performance of twelve different feature selection methods when subjected to an increase in the amount of dropout noise. Following feature selection, trajectory preservation was quantitatively assessed according to several metrics: the precision of differentially expressed genes at *k* selected genes (top), *k*-NN classification accuracy (middle), and pseudotime correlation (bottom) across 10 random trails. Error bars/ bands represent the standard deviation. * indicates the method with the highest median score. - indicates that the method identified no features.

**Supplementary Figure 10:**
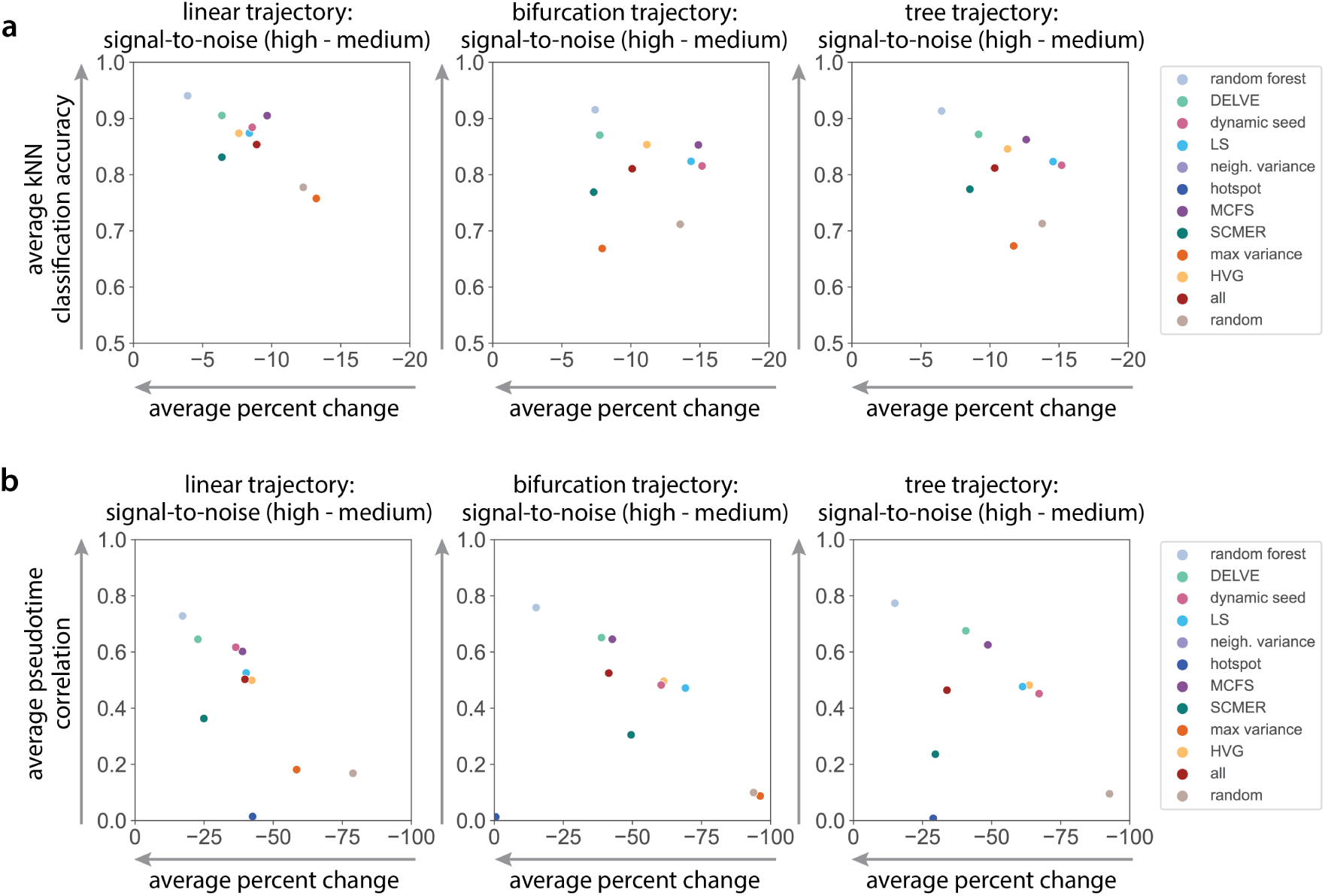
Comparison of the robustness of feature selection methods on inferring differentiation trajectories under a reduction in the signal-to-noise ratio. (a) Average *k*-NN classification accuracy vs. average percent change in *k*-NN classification accuracy as the signal-to-noise ratio decreased (high to medium) and the mean-variance relationship amongst genes increased. The signal-to-noise ratio was altered by modifying the biological coefficient of variation parameter within Splatter [72] (high: BCV = 0.1, medium: BCV = 0.25). (b) Average pseudotime correlation vs. average percent change in pseudotime correlation as the signal-to-noise ratio decreased (high to medium).

**Supplementary Figure 11:**
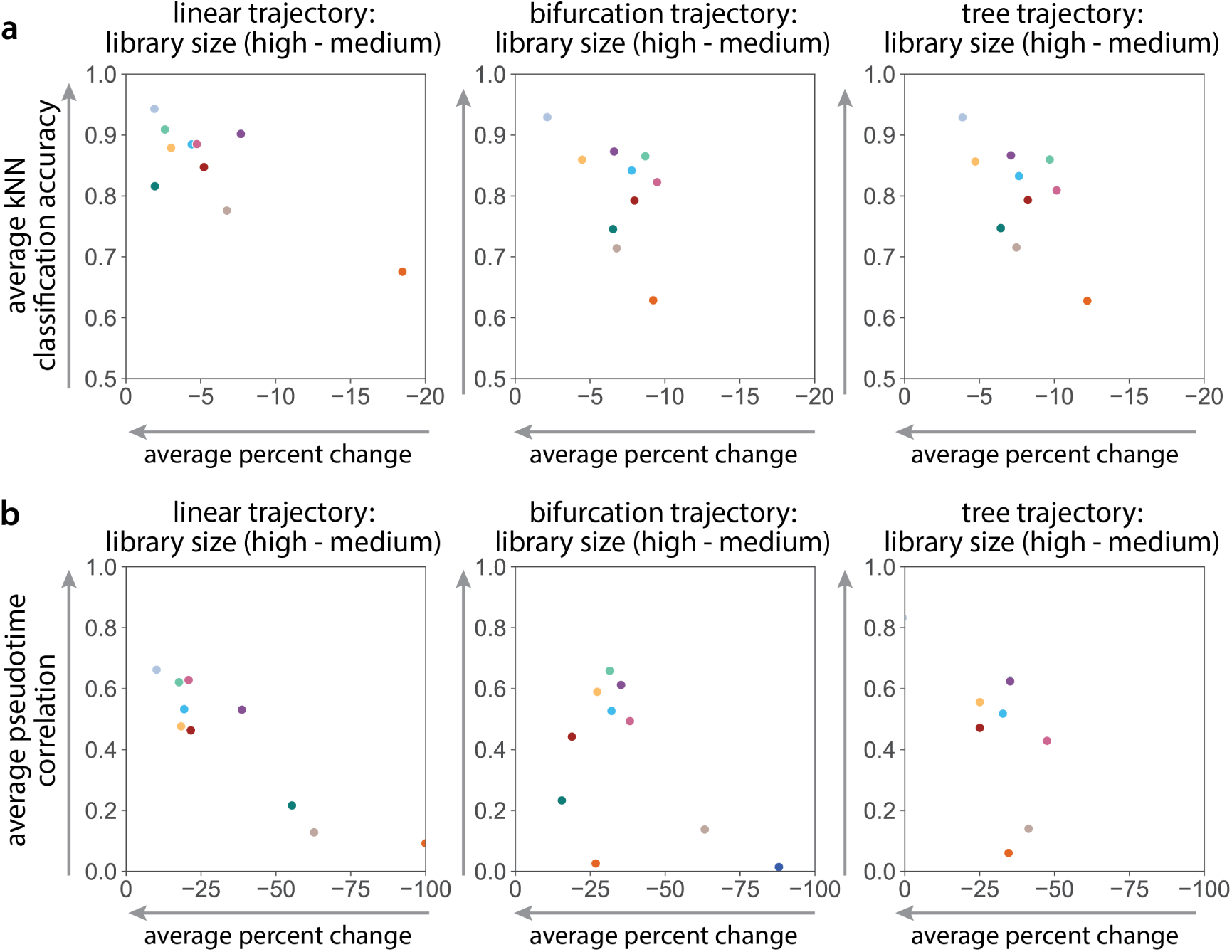
Comparison of the robustness of feature selection methods on inferring differentiation trajectories under library size noise corruption. (a) Average *k*-NN classification accuracy vs. average percent change in *k*-NN classification accuracy as the total mRNA count decreased (high to medium). Library size was reduced by modifying the location parameter in the log-normal distribution in Splatter [72] that specifies library size scaling factors (high: location = 12, medium: location = 11). (b) Average pseudotime correlation vs. average percent change in pseudotime correlation as the total mRNA count decreased (high to medium).

**Supplementary Figure 12:**
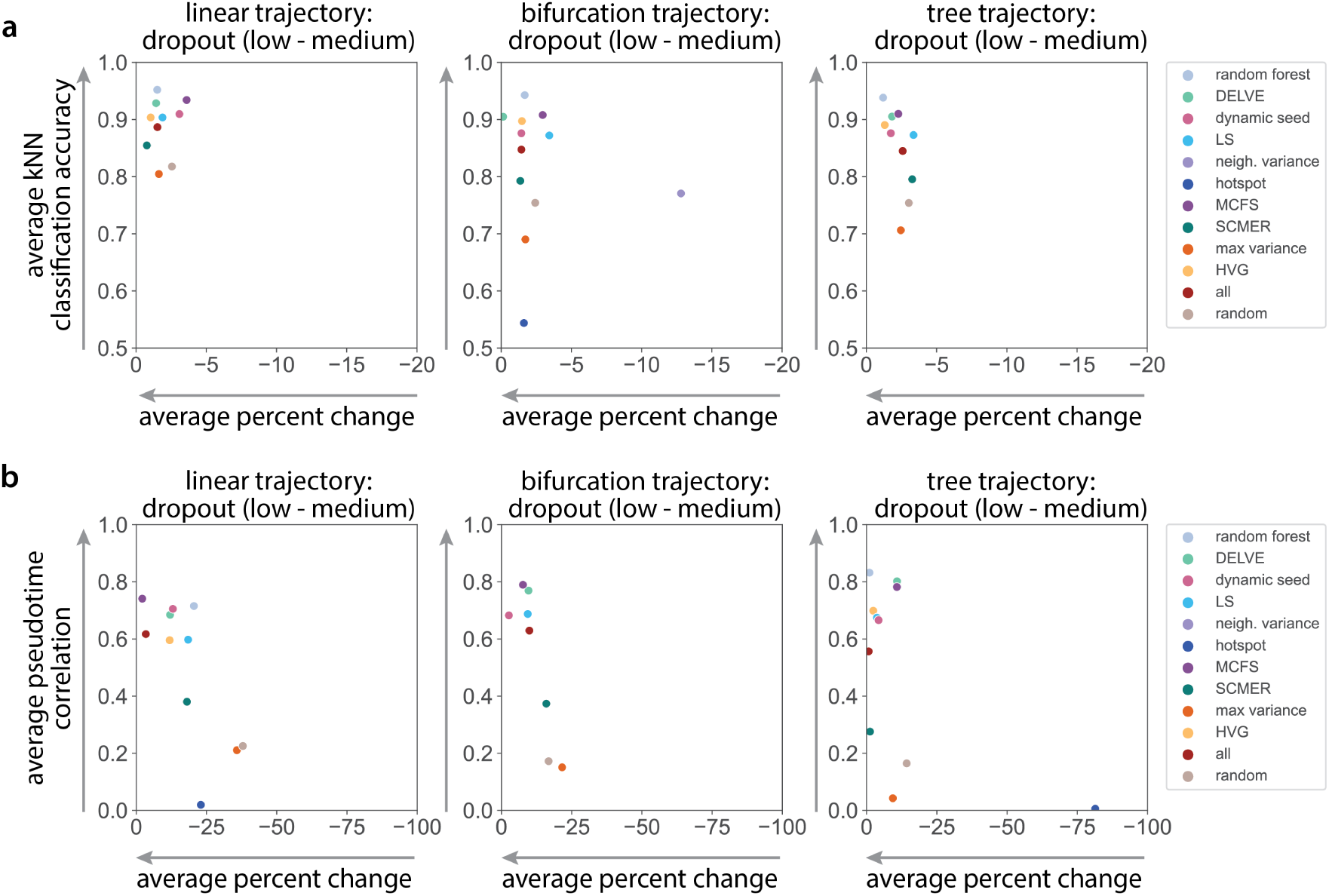
Comparison of the robustness of feature selection methods on inferring differentiation trajectories under dropout noise corruption. (a) Average *k*-NN classification accuracy vs. average percent change in *k*-NN classification accuracy as the amount of dropout or sparsity increased (low to medium). Technical dropout was simulated by undersampling mRNA counts by sampling from a binomial distribution with the scale parameter or dropout rate proportional to the mean expression of each gene (low: *λ* = 0, medium: *λ* = 0.05). (b) Average pseudotime correlation vs. average percent change in pseudotime correlation as the amount of dropout or sparsity increased (low to medium).

**Supplementary Figure 13:**
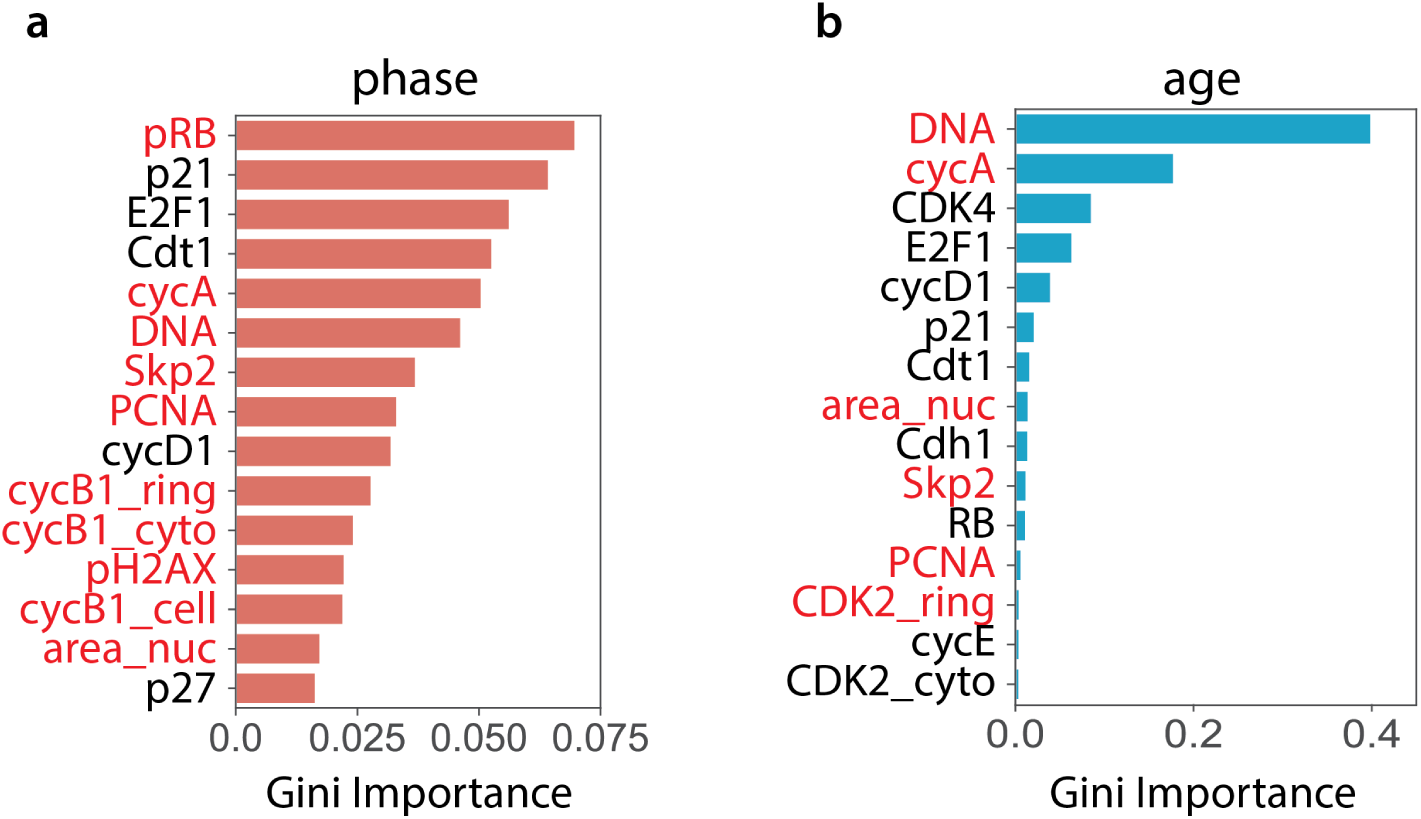
Validation of DELVE seed selection on retinal pigmented epithelial (RPE) cell cycle dataset. Top ranked features identified by a (a) random forest classifier trained on ground truth cell cycle phase annotations or (b) random forest regressor trained on ground truth cell cycle age measurements. Features highlighted in red were also identified by DELVE seed selection (See Figure 4a heatmap).

**Supplementary Figure 14:**
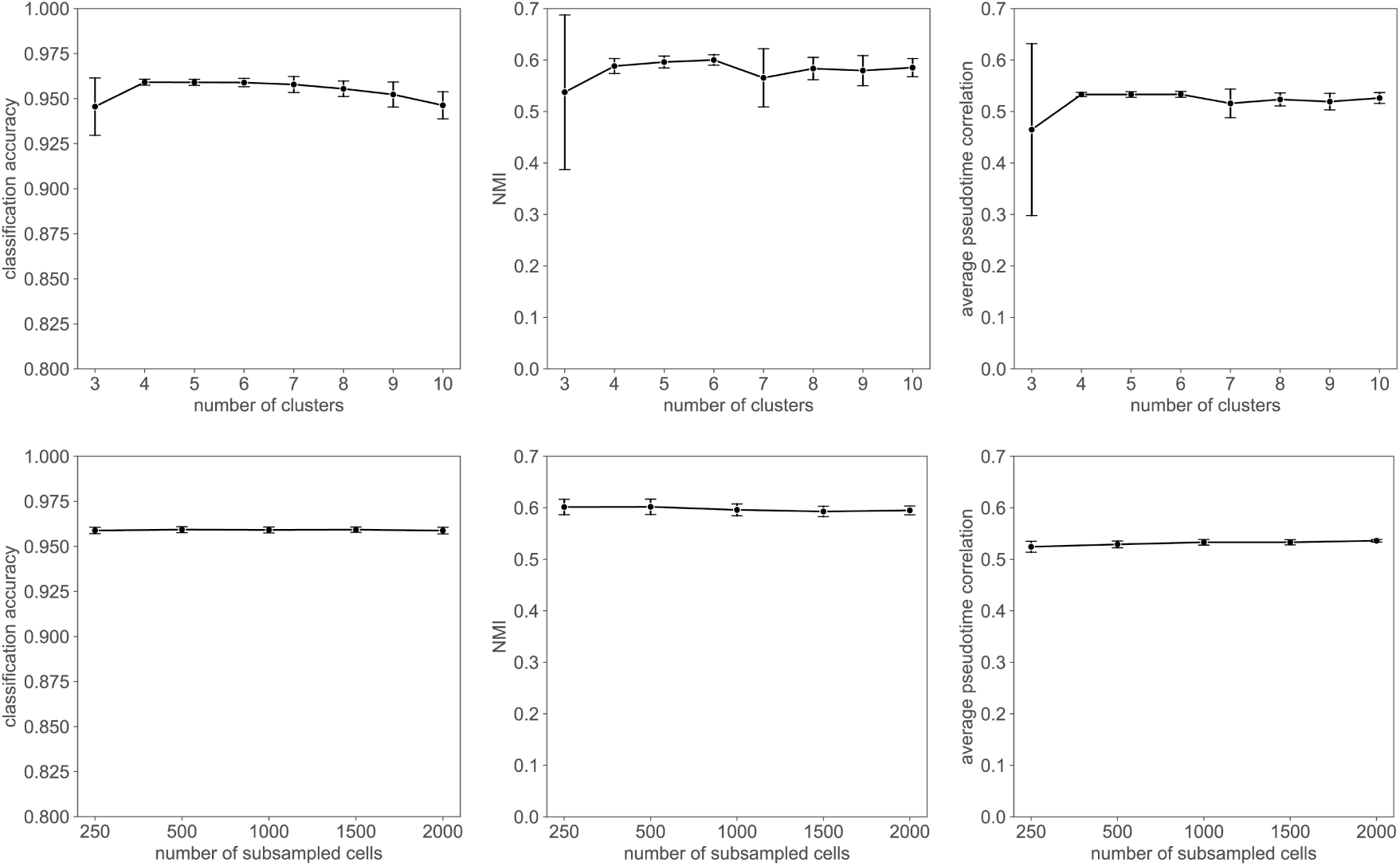
DELVE is robust to changes in hyperparameters for the retinal pigmented epithelial (RPE) cell cycle dataset. DELVE achieves similar classification accuracy, normalized mutual information (NMI) clustering score, and pseudotime correlation across a range of (top) cluster sizes and (bottom) subsampled cells. Plots show results over 20 random trials. Error bars represent the standard deviation.

**Supplementary Figure 15:**
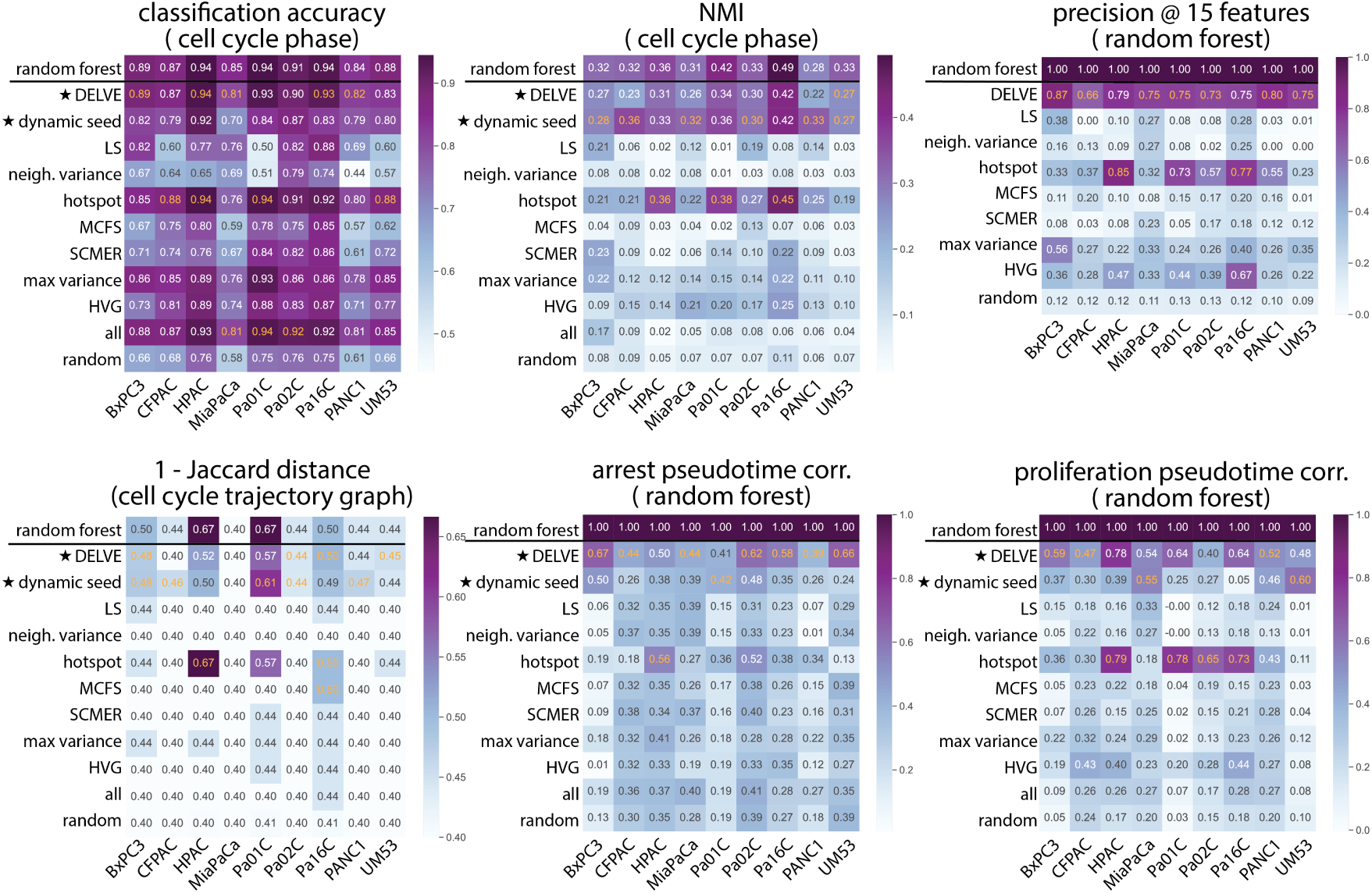
Feature selection method performance on pancreatic adenocarcinoma (PDAC) cell cycle datasets. Performance of twelve feature selection methods on preserving cell cycle trajectories from 9 PDAC cancer cell lines (BxPC3, CFPAC, HPAC, MiaPaCa, Pa02C, Pa01C, Pa16C, PANC1, and UM53) profiled with protein immunofluorescence imaging. Following feature selection, cell cycle preservation was quantitatively assessed according to several metrics including: support vector machine classification accuracy to the ground truth phase annotations, normalized mutual information (NMI) clustering score to ground truth phase annotations, precision of cell cycle phase-specific imaging-derived features as measured by a random forest classifier trained on ground truth phase annotations, Jaccard distance between predicted cell cycle trajectory graphs and a ground truth reference cell cycle trajectory curated from the literature, and the Spearman rank correlation between estimated pseudotime and the ground truth as measured by a random forest classifier trained on ground truth phase annotations. Heatmaps show the average performance. Approaches with the highest average score are highlighted in yellow.

**Supplementary Figure 16:**
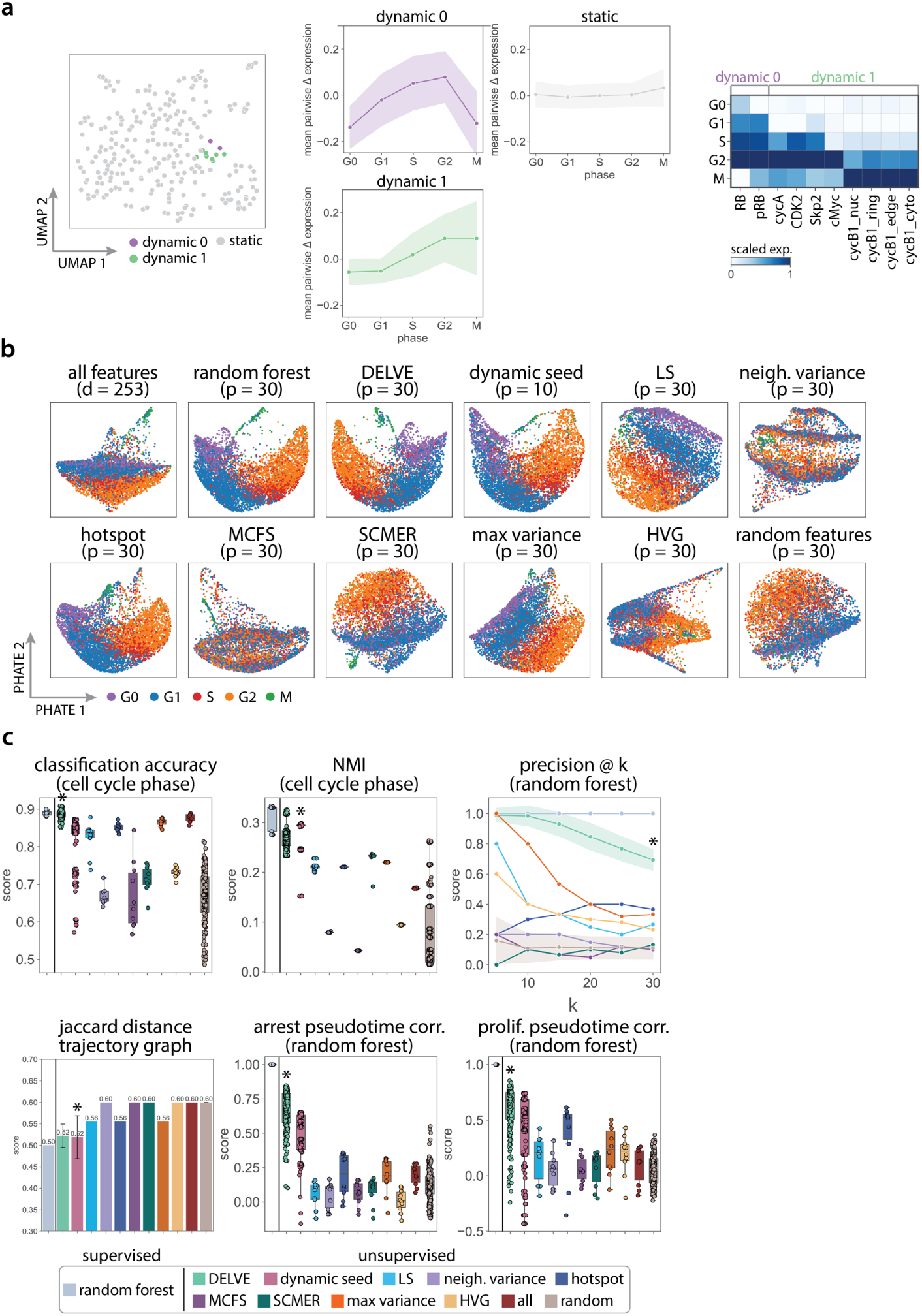
DELVE recovers BxPC3 pancreatic adenocarcinoma cell cycle trajectories in protein immunofluorescence imaging data. BxPC3 cells were profiled with protein immunofluorescence imaging to measure 63 core cell cycle effectors resulting in a dataset with *d* = 253 imaging-derived features. (a) DELVE identified two modules of dynamic features representing a minimum cell cycle. (a left) UMAP visualization of image-derived features where each point indicates a dynamic or static feature identified by the model. (a middle) The average pairwise change in expression for features within a module ordered across ground truth cell cycle phase annotations. (a right) Heatmap illustrating the standardized average expression of dynamic seed features across cell cycle phases. (b) Feature selection was performed to select the top *p* = 30 ranked features. Example PHATE [75] visualizations of cell cycle trajectories for twelve feature selection approaches. (c) Quantitative assessment of twelve feature selection methods on preserving cell cycle phases and phase transitions according to several metrics including: support vector machine classification accuracy to the ground truth phase annotations, normalized mutual information (NMI) clustering score to ground truth phase annotations, precision of cell cycle phase-specific imaging-derived features as measured by a random forest classifier trained on ground truth phase annotations, Jaccard distance between predicted cell cycle trajectory graphs and a ground truth reference cell cycle trajectory curated from the literature, and the Spearman rank correlation between estimated pseudotime and the ground truth as measured by a random forest classifier trained on ground truth phase annotations. Boxplots show the results over 10 random trials, error bands represent the standard deviation, and * indicates the method with the highest median score.

**Supplementary Figure 17:**
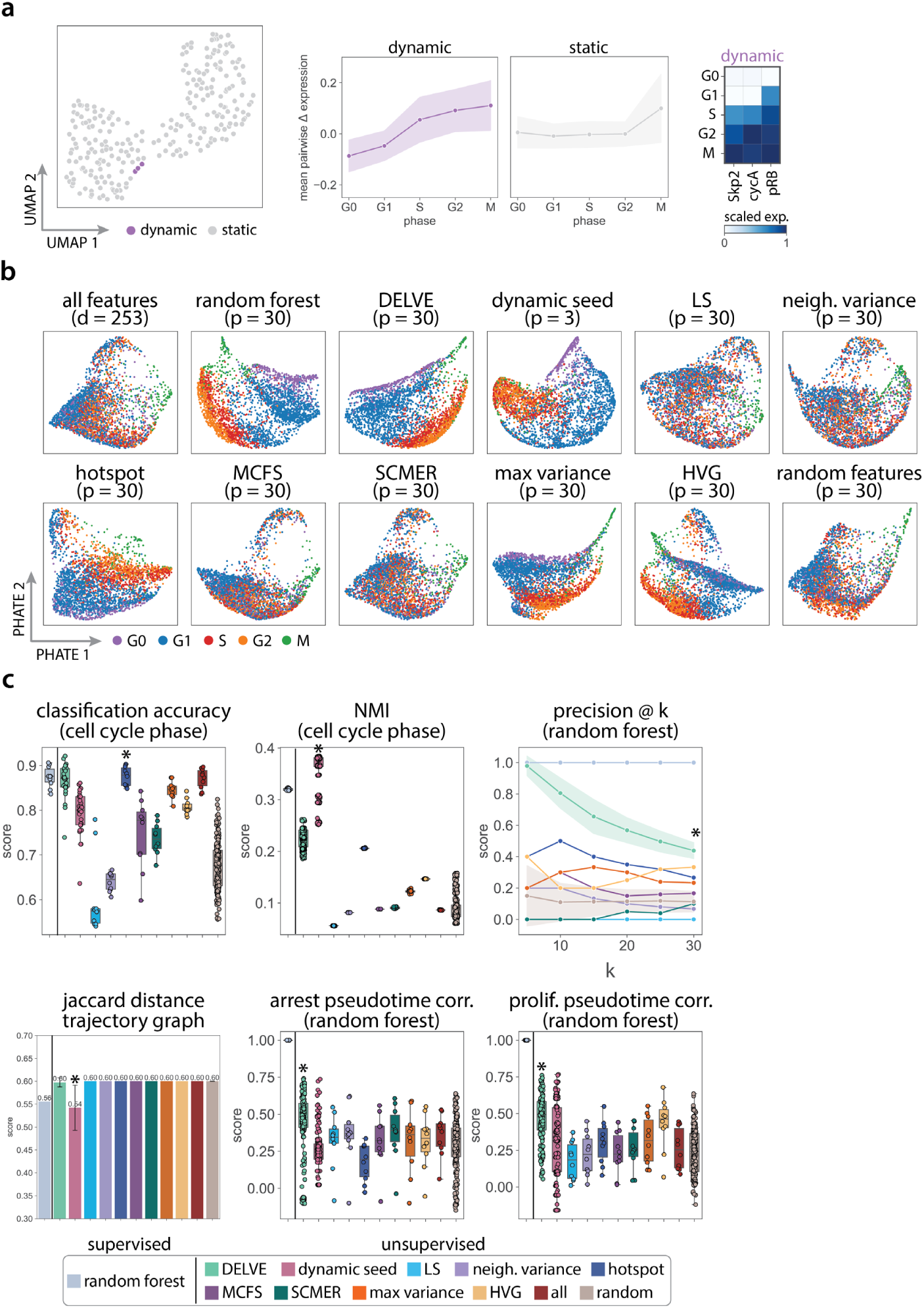
DELVE recovers CFPAC pancreatic adenocarcinoma cell cycle trajectories in protein immunofluorescence imaging data. CFPAC cells were profiled with protein immunofluorescence imaging to measure 63 core cell cycle effectors resulting in a dataset with *d* = 253 imaging-derived features. (a) DELVE identified one module of dynamic features representing a minimum cell cycle. (a left) UMAP visualization of image-derived features where each point indicates a dynamic or static feature identified by the model. (a middle) The average pairwise change in expression for features within a module ordered across ground truth cell cycle phase annotations. (a right) Heatmap illustrating the standardized average expression of dynamic seed features across cell cycle phases. (b) Feature selection was performed to select the top *p* = 30 ranked features. Example PHATE [75] visualizations of cell cycle trajectories for twelve feature selection approaches. (c) Quantitative assessment of twelve feature selection methods on preserving cell cycle phases and phase transitions according to several metrics including: support vector machine classification accuracy to the ground truth phase annotations, normalized mutual information (NMI) clustering score to ground truth phase annotations, precision of cell cycle phase-specific imaging-derived features as measured by a random forest classifier trained on ground truth phase annotations, Jaccard distance between predicted cell cycle trajectory graphs and a ground truth reference cell cycle trajectory curated from the literature, and the Spearman rank correlation between estimated pseudotime and the ground truth as measured by a random forest classifier trained on ground truth phase annotations. Boxplots show the results over 10 random trials, error bands represent the standard deviation, and * indicates the method with the highest median score.

**Supplementary Figure 18:**
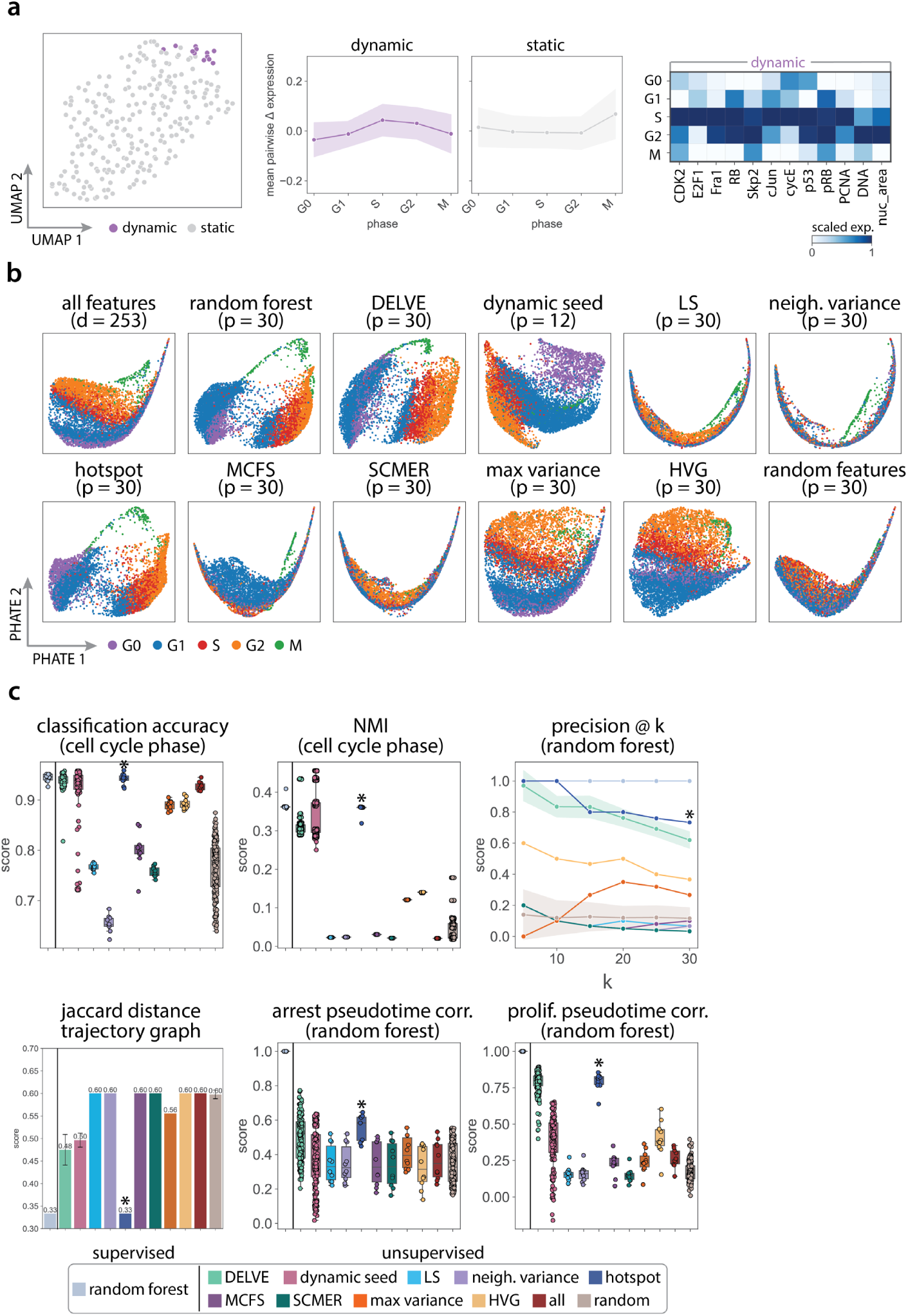
DELVE recovers HPAC pancreatic adenocarcinoma cell cycle trajectories in protein immunofluorescence imaging data. HPAC cells were profiled with protein immunofluorescence imaging to measure 63 core cell cycle effectors resulting in a dataset with *d* = 253 imaging-derived features. (a) DELVE identified one module of dynamic features representing a minimum cell cycle. (a left) UMAP visualization of image-derived features where each point indicates a dynamic or static feature identified by the model. (a middle) The average pairwise change in expression for features within a module ordered across ground truth cell cycle phase annotations. (a right) Heatmap illustrating the standardized average expression of dynamic seed features across cell cycle phases. (b) Feature selection was performed to select the top *p* = 30 ranked features. Example PHATE [75] visualizations of cell cycle trajectories for twelve feature selection approaches. (c) Quantitative assessment of twelve feature selection methods on preserving cell cycle phases and phase transitions according to several metrics including: support vector machine classification accuracy to the ground truth phase annotations, normalized mutual information (NMI) clustering score to ground truth phase annotations, precision of cell cycle phase-specific imaging-derived features as measured by a random forest classifier trained on ground truth phase annotations, Jaccard distance between predicted cell cycle trajectory graphs and a ground truth reference cell cycle trajectory curated from the literature, and the Spearman rank correlation between estimated pseudotime and the ground truth as measured by a random forest classifier trained on ground truth phase annotations. Boxplots show the results over 10 random trials, error bands represent the standard deviation, and * indicates the method with the highest median score.

**Supplementary Figure 19:**
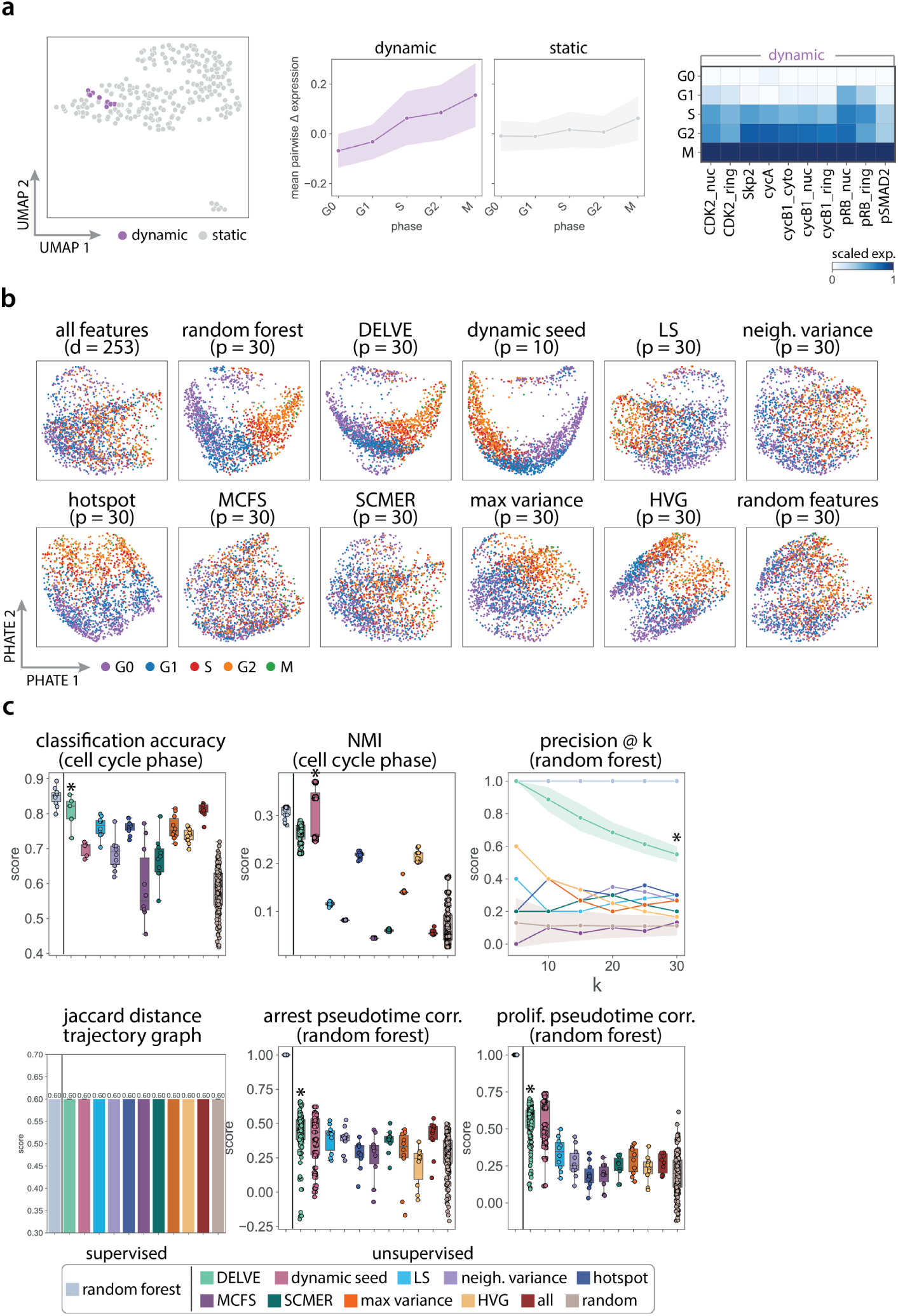
DELVE recovers MiaPaCa pancreatic adenocarcinoma cell cycle trajectories in protein immunofluorescence imaging data. MiaPaCa cells were profiled with protein immunofluorescence imaging to measure 63 core cell cycle effectors resulting in a dataset with *d* = 253 imaging-derived features. (a) DELVE identified one module of dynamic features representing a minimum cell cycle. (a left) UMAP visualization of image-derived features where each point indicates a dynamic or static feature identified by the model. (a middle) The average pairwise change in expression for features within a module ordered across ground truth cell cycle phase annotations. (a right) Heatmap illustrating the standardized average expression of dynamic seed features across cell cycle phases. (b) Feature selection was performed to select the top *p* = 30 ranked features. Example PHATE [75] visualizations of cell cycle trajectories for twelve feature selection approaches. (c) Quantitative assessment of twelve feature selection methods on preserving cell cycle phases and phase transitions according to several metrics including: support vector machine classification accuracy to the ground truth phase annotations, normalized mutual information (NMI) clustering score to ground truth phase annotations, precision of cell cycle phase-specific imaging-derived features as measured by a random forest classifier trained on ground truth phase annotations, Jaccard distance between predicted cell cycle trajectory graphs and a ground truth reference cell cycle trajectory curated from the literature, and the Spearman rank correlation between estimated pseudotime and the ground truth as measured by a random forest classifier trained on ground truth phase annotations. Boxplots show the results over 10 random trials, error bands represent the standard deviation, and * indicates the method with the highest median score.

**Supplementary Figure 20:**
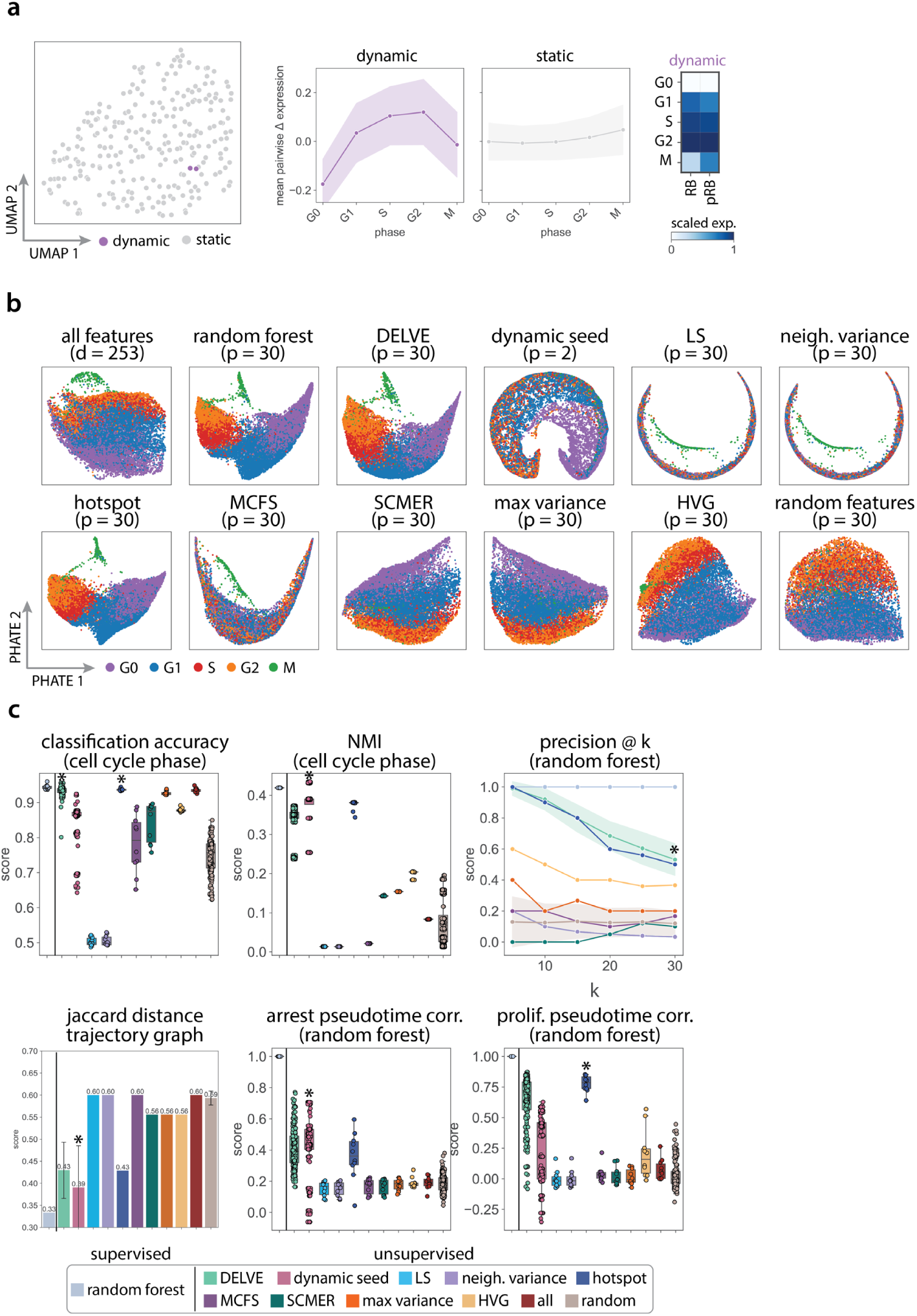
DELVE recovers Pa01C pancreatic adenocarcinoma cell cycle trajectories in protein immunofluorescence imaging data. Pa01C cells were profiled with protein immunofluorescence imaging to measure 63 core cell cycle effectors resulting in a dataset with *d* = 253 imaging-derived features. (a) DELVE identified one module of dynamic features representing a minimum cell cycle. (a left) UMAP visualization of image-derived features where each point indicates a dynamic or static feature identified by the model. (a middle) The average pairwise change in expression for features within a module ordered across ground truth cell cycle phase annotations. (a right) Heatmap illustrating the standardized average expression of dynamic seed features across cell cycle phases. (b) Feature selection was performed to select the top *p* = 30 ranked features. Example PHATE [75] visualizations of cell cycle trajectories for twelve feature selection approaches. (c) Quantitative assessment of twelve feature selection methods on preserving cell cycle phases and phase transitions according to several metrics including: support vector machine classification accuracy to the ground truth phase annotations, normalized mutual information (NMI) clustering score to ground truth phase annotations, precision of cell cycle phase-specific imaging-derived features as measured by a random forest classifier trained on ground truth phase annotations, Jaccard distance between predicted cell cycle trajectory graphs and a ground truth reference cell cycle trajectory curated from the literature, and the Spearman rank correlation between estimated pseudotime and the ground truth as measured by a random forest classifier trained on ground truth phase annotations. Boxplots show the results over 10 random trials, error bands represent the standard deviation, and * indicates the method with the highest median score.

**Supplementary Figure 21:**
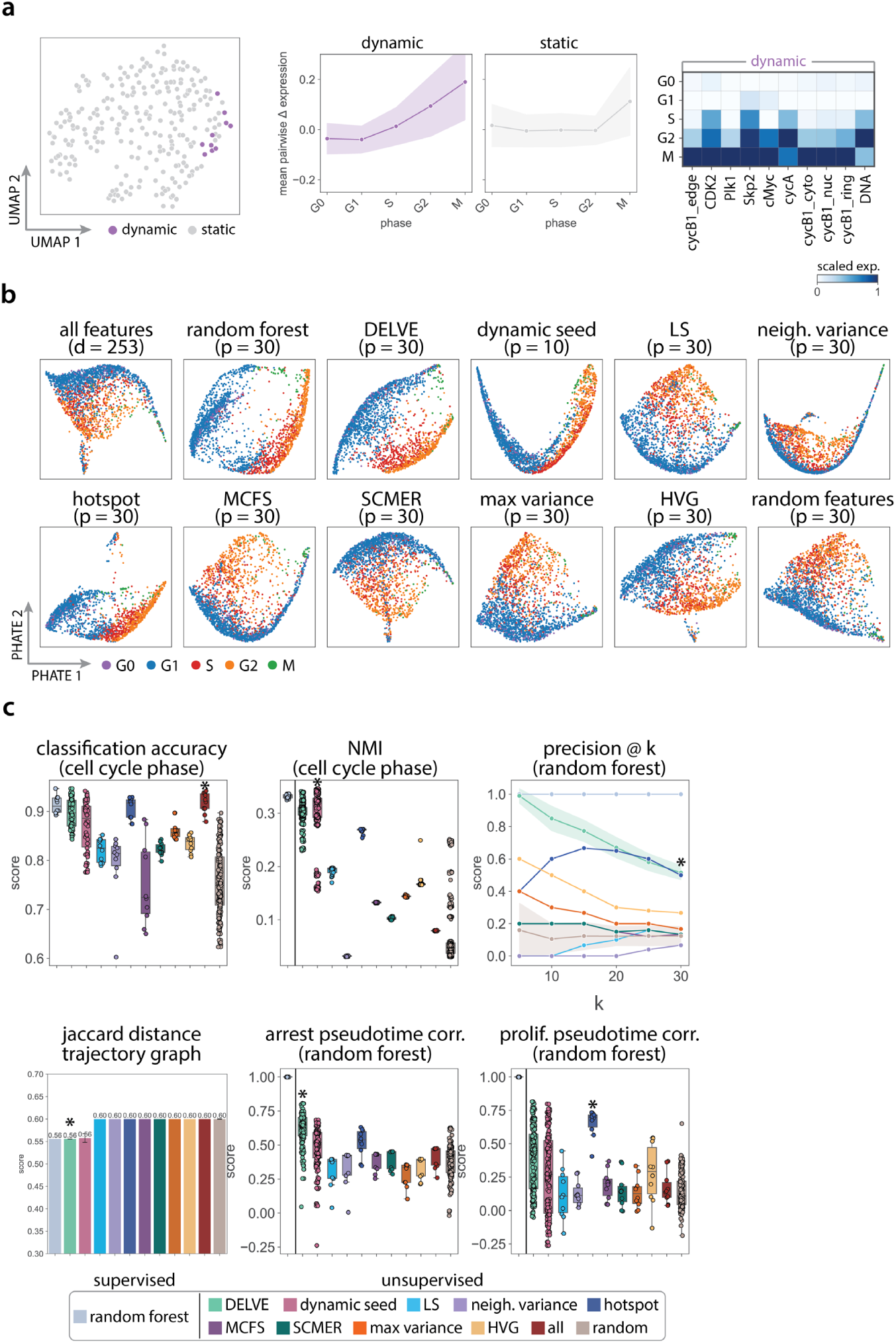
DELVE recovers Pa02C pancreatic adenocarcinoma cell cycle trajectories in protein immunofluorescence imaging data. Pa02C cells were profiled with protein immunofluorescence imaging to measure 63 core cell cycle effectors resulting in a dataset with *d* = 253 imaging-derived features. (a) DELVE identified one module of dynamic features representing a minimum cell cycle. (a left) UMAP visualization of image-derived features where each point indicates a dynamic or static feature identified by the model. (a middle) The average pairwise change in expression for features within a module ordered across ground truth cell cycle phase annotations. (a right) Heatmap illustrating the standardized average expression of dynamic seed features across cell cycle phases. (b) Feature selection was performed to select the top *p* = 30 ranked features. Example PHATE [75] visualizations of cell cycle trajectories for twelve feature selection approaches. (c) Quantitative assessment of twelve feature selection methods on preserving cell cycle phases and phase transitions according to several metrics including: support vector machine classification accuracy to the ground truth phase annotations, normalized mutual information (NMI) clustering score to ground truth phase annotations, precision of cell cycle phase-specific imaging-derived features as measured by a random forest classifier trained on ground truth phase annotations, Jaccard distance between predicted cell cycle trajectory graphs and a ground truth reference cell cycle trajectory curated from the literature, and the Spearman rank correlation between estimated pseudotime and the ground truth as measured by a random forest classifier trained on ground truth phase annotations. Boxplots show the results over 10 random trials, error bands represent the standard deviation, and * indicates the method with the highest median score.

**Supplementary Figure 22:**
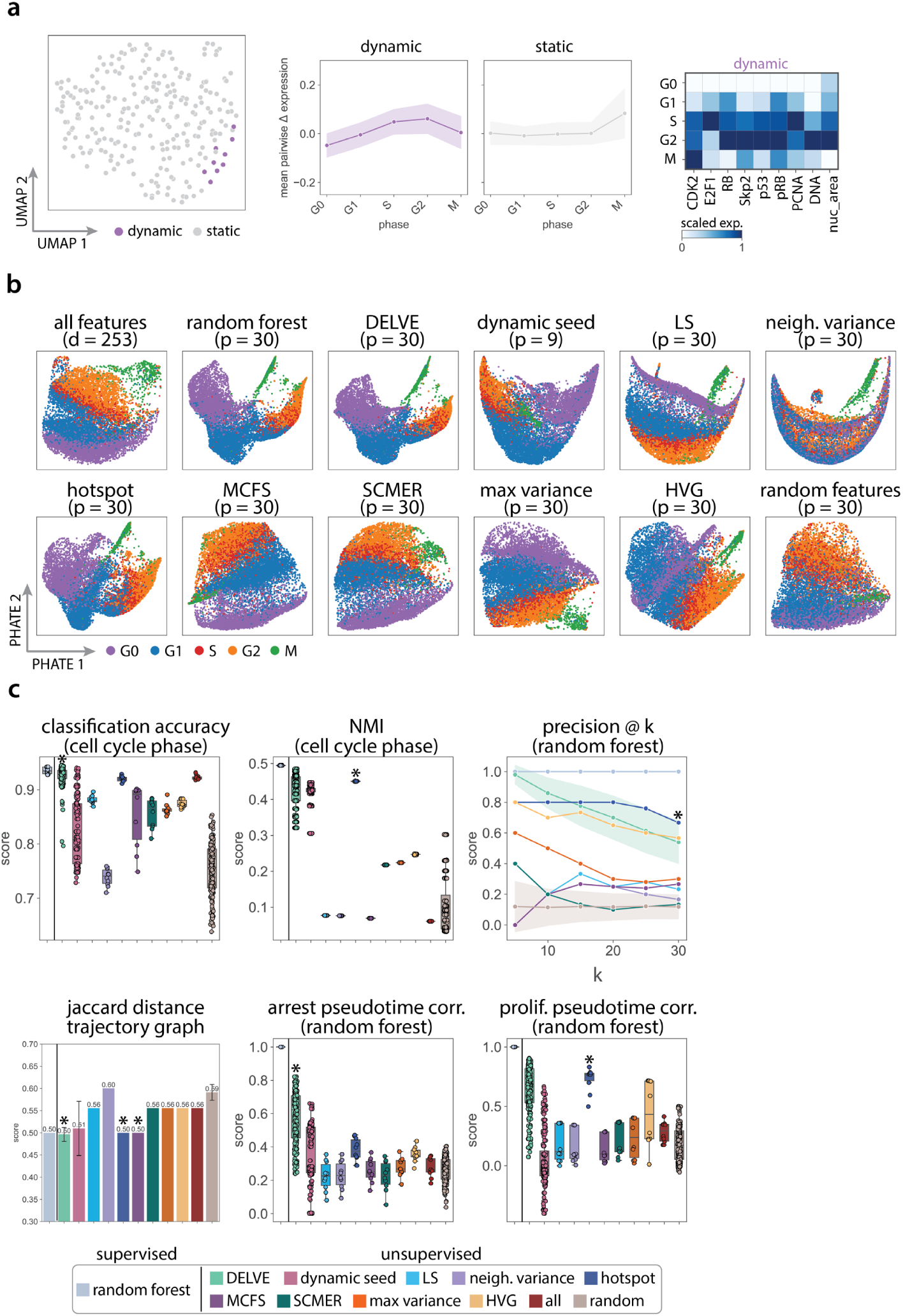
DELVE recovers Pa16C pancreatic adenocarcinoma cell cycle trajectories in protein immunofluorescence imaging data. Pa16C cells were profiled with protein immunofluorescence imaging to measure 63 core cell cycle effectors resulting in a dataset with *d* = 253 imaging-derived features. (a) DELVE identified one module of dynamic features representing a minimum cell cycle. (a left) UMAP visualization of image-derived features where each point indicates a dynamic or static feature identified by the model. (a middle) The average pairwise change in expression for features within a module ordered across ground truth cell cycle phase annotations. (a right) Heatmap illustrating the standardized average expression of dynamic seed features across cell cycle phases. (b) Feature selection was performed to select the top *p* = 30 ranked features. Example PHATE [75] visualizations of cell cycle trajectories for twelve feature selection approaches. (c) Quantitative assessment of twelve feature selection methods on preserving cell cycle phases and phase transitions according to several metrics including: support vector machine classification accuracy to the ground truth phase annotations, normalized mutual information (NMI) clustering score to ground truth phase annotations, precision of cell cycle phase-specific imaging-derived features as measured by a random forest classifier trained on ground truth phase annotations, Jaccard distance between predicted cell cycle trajectory graphs and a ground truth reference cell cycle trajectory curated from the literature, and the Spearman rank correlation between estimated pseudotime and the ground truth as measured by a random forest classifier trained on ground truth phase annotations. Boxplots show the results over 10 random trials, error bands represent the standard deviation, and * indicates the method with the highest median score.

**Supplementary Figure 23:**
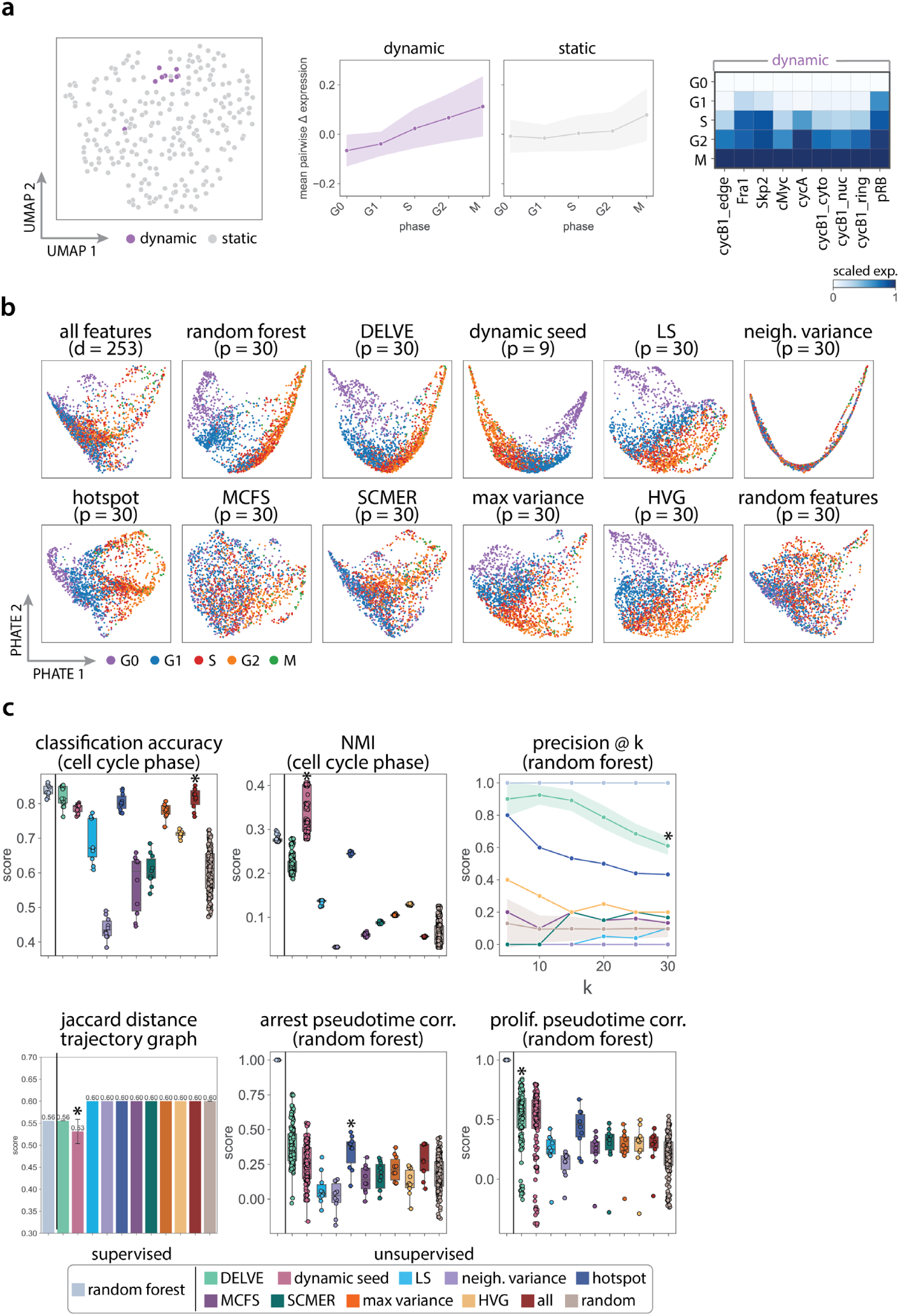
DELVE recovers PANC1 pancreatic adenocarcinoma cell cycle trajectories in protein immunofluorescence imaging data. PANC1 cells were profiled with protein immunofluorescence imaging to measure 63 core cell cycle effectors resulting in a dataset with *d* = 253 imaging-derived features. (a) DELVE identified one module of dynamic features representing a minimum cell cycle. (a left) UMAP visualization of image-derived features where each point indicates a dynamic or static feature identified by the model. (a middle) The average pairwise change in expression for features within a module ordered across ground truth cell cycle phase annotations. (a right) Heatmap illustrating the standardized average expression of dynamic seed features across cell cycle phases. (b) Feature selection was performed to select the top *p* = 30 ranked features. Example PHATE [75] visualizations of cell cycle trajectories for twelve feature selection approaches. (c) Quantitative assessment of twelve feature selection methods on preserving cell cycle phases and phase transitions according to several metrics including: support vector machine classification accuracy to the ground truth phase annotations, normalized mutual information (NMI) clustering score to ground truth phase annotations, precision of cell cycle phase-specific imaging-derived features as measured by a random forest classifier trained on ground truth phase annotations, Jaccard distance between predicted cell cycle trajectory graphs and a ground truth reference cell cycle trajectory curated from the literature, and the Spearman rank correlation between estimated pseudotime and the ground truth as measured by a random forest classifier trained on ground truth phase annotations. Boxplots show the results over 10 random trials, error bands represent the standard deviation, and * indicates the method with the highest median score.

**Supplementary Figure 24:**
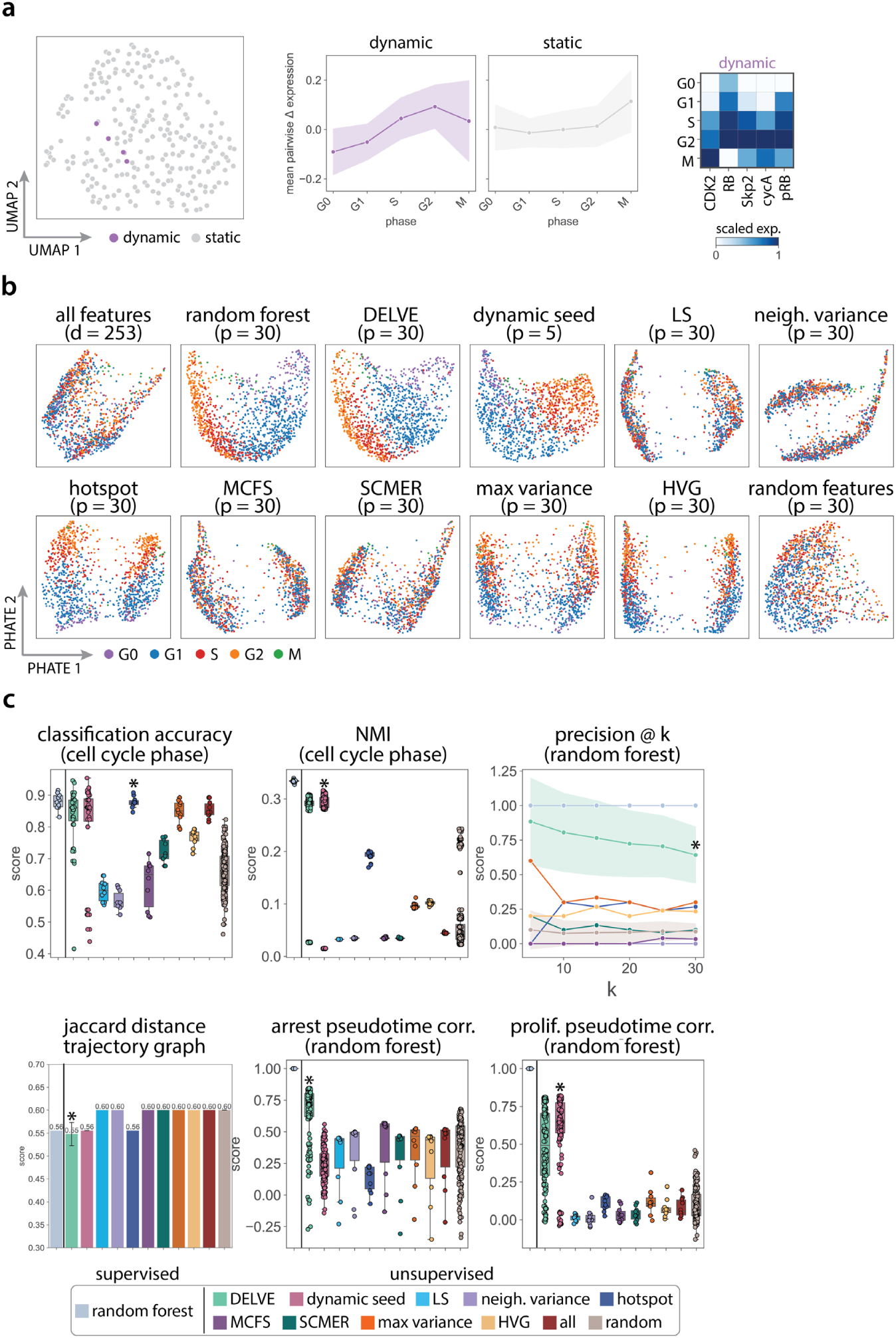
DELVE recovers UM53 pancreatic adenocarcinoma cell cycle trajectories in protein immunofluorescence imaging data. UM53 cells were profiled with protein immunofluorescence imaging to measure 63 core cell cycle effectors resulting in a dataset with *d* = 253 imaging-derived features. (a) DELVE identified one module of dynamic features representing a minimum cell cycle. (a left) UMAP visualization of image-derived features where each point indicates a dynamic or static feature identified by the model. (a middle) The average pairwise change in expression for features within a module ordered across ground truth cell cycle phase annotations. (a right) Heatmap illustrating the standardized average expression of dynamic seed features across cell cycle phases. (b) Feature selection was performed to select the top *p* = 30 ranked features. Example PHATE [75] visualizations of cell cycle trajectories for twelve feature selection approaches. (c) Quantitative assessment of twelve feature selection methods on preserving cell cycle phases and phase transitions according to several metrics including: support vector machine classification accuracy to the ground truth phase annotations, normalized mutual information (NMI) clustering score to ground truth phase annotations, precision of cell cycle phase-specific imaging-derived features as measured by a random forest classifier trained on ground truth phase annotations, Jaccard distance between predicted cell cycle trajectory graphs and a ground truth reference cell cycle trajectory curated from the literature, and the Spearman rank correlation between estimated pseudotime and the ground truth as measured by a random forest classifier trained on ground truth phase annotations. Boxplots show the results over 10 random trials, error bands represent the standard deviation, and * indicates the method with the highest median score.

